# Resolving Sub-Microsecond Conformational Dynamics of Vertical Nucleic Acids on Graphene

**DOI:** 10.64898/2026.06.06.730580

**Authors:** Lars Richter, Jakob Hartmann, Leo Christanell, Tim Schröder, Alan M. Szalai, Benjamin P. Fingerhut, Philip Tinnefeld

## Abstract

The function of nucleic acids is governed not only by their structure but also by their dynamics. At the molecular scale, transitions between functional structural states are superimposed on rapid thermal fluctuations, resulting in an intricate interplay that is challenging to resolve experimentally, particularly at the single-molecule level. Here, we introduce a novel approach for unraveling sub-microsecond dynamics in oligonucleotides, enabling direct observation of fluctuations in single DNA molecules. By immobilizing nucleic acids vertically on graphene and exploiting distance-dependent graphene energy transfer of fluorescent molecules attached to the DNA, we relate fluctuations in fluorescence intensity to biomolecular dynamics. We show that ionic strength modulates the fluctuations and that structural defects in DNA, such as nucleotide gaps or mismatches, alter the measured dynamics. The experimental findings are complemented by atomistic molecular dynamics simulations and kinetic Monte Carlo simulations, establishing a direct link between theoretical predictions of structure and dynamics and experimentally accessible fluctuation timescales. Overall, our findings advance the understanding of how thermal fluctuations affect oligonucleotides and are modulated by both external and internal stimuli.

## Introduction

In biophysical models, double-stranded deoxyribonucleic acid (dsDNA) is commonly described as a semiflexible polymer whose mechanical behavior follows the worm-like chain (WLC) model.^1^ The persistence length (l_p_) quantifies its bending stiffness and is approximately 50 nm for dsDNA molecules, corresponding to ∼150 base pairs (bp).^2,3^ Because l_p_ defines the characteristic scale of flexibility, bending DNA over significantly shorter distances requires substantial energy, making the formation of sharp bends or short loops intrinsically unfavorable.^4^ However, it has been shown that DNA can adopt strongly bent configurations, including looped structures, even at lengths as short as ∼100 bp.^5–7^ Such sharp DNA bending is not anticipated by the WLC model but such kinks can arise from localized structural distortions within the polymer.^8,9^ The conformational dynamics of DNA are further influenced by base sequence, chemical modifications and ionic environment.^5,10^ Additionally, DNA-protein interactions often create defined bending or structural deformations, making the intrinsic mechanical flexibility of DNA a central determinant for understanding the mechanisms by which these complexes realize their biological function.^11–13^

As structural changes of dsDNA at room temperature are on the nanometer scale, only a few experimental methods have been applied to reveal them. Experimentally, for example, nuclear magnetic resonance spectroscopy or temperature-jump spectroscopy provide access to the subtle spatial scale combined with fast, sub-millisecond dynamics.^14,15^ On the level of single molecules, bendability of short strands was visualized by trapping rare, looped states. Fast conformational fluctuations are expected in the nano-to microsecond range prohibiting their real-time visualization with realistic single-molecule fluorescence count rates.^16,17^ Beyond real-time measurements, intensity fluctuations for correlation analysis were induced by fluorescence resonance energy transfer (FRET) when the dynamics are correlated with distance changes between a donor and an acceptor fluorophore.^18^ However, FRET is limited to selectively report on DNA dynamics as its distance range is small, the dyes are linked *via* flexible linkers, and it relies on a single energy transfer acceptor that causes FRET to depend on torsional as well as bending dynamics.^19,20^ Further, the use of two dye molecules in FRET comes along with photophysical amplitudes in the correlation signal, complicating the clear assignment of biomolecular dynamics.^21,22^

Recently, we introduced Graphene Energy Transfer with vertical Nucleic Acids (GETvNA).^23^ On the one hand, GETvNA uses the property of graphene to act as an energy transfer acceptor with a d^-4^ scaling law and a critical distance of ∼18 nm for 50% energy transfer (Figure 1A).^24,25^ This energy transfer reports the distance of a single dye molecule to the graphene surface by intensity or fluorescence lifetime measurements. On the other hand, it was discovered that dsDNA adopts a perpendicular orientation on graphene when immobilized with a single-stranded DNA overhang that serves as glue through π-stacking interactions between graphene and nucleobases (Figure 1B). GETvNA was used to study DNA conformations with sub-nanometer precision on a confocal microscope.^26^ Structural modifications of dsDNA such as poly-A bulges and A-tracts, or external factors such as DNA-binding proteins were visualized with a time resolution in the millisecond range. However, biomolecular dynamics are not constrained to these timescales.^27,28^

**Figure 1.**
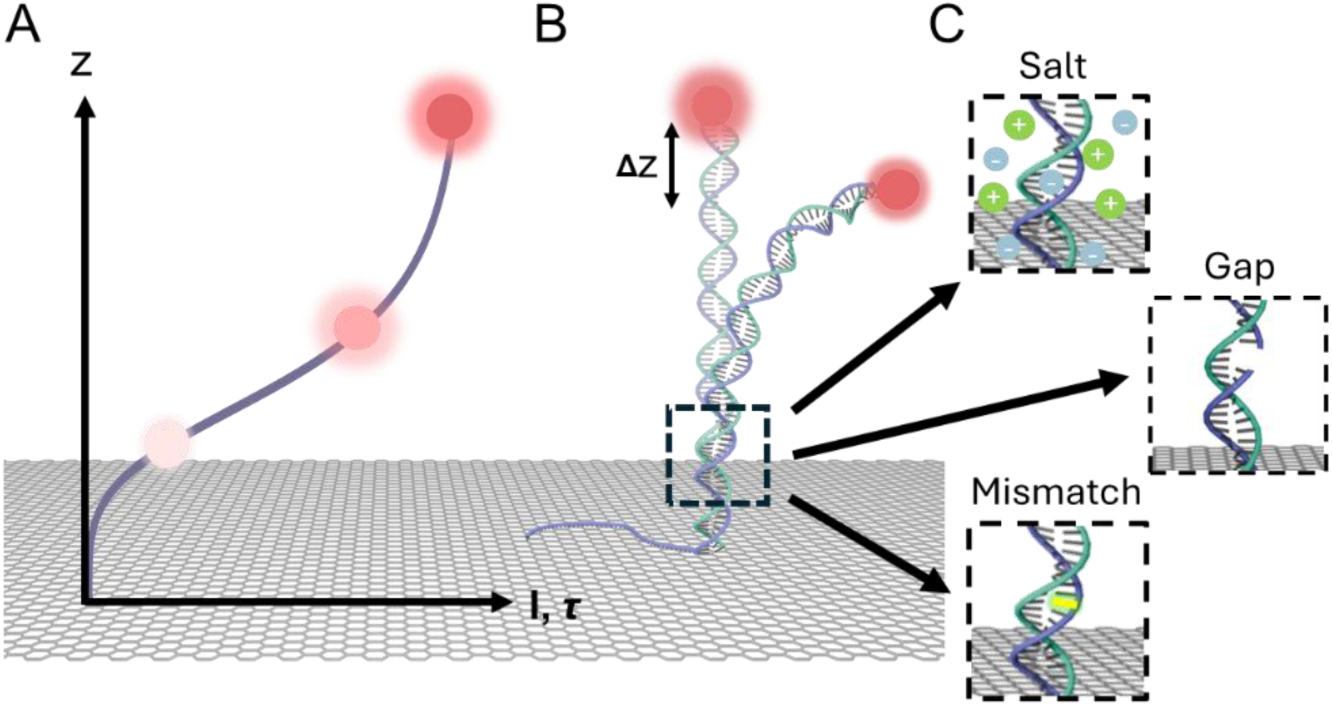
Conformational dynamics in DNA by GETvNA. **(A)** Distance dependency for the energy transfer from fluorescent molecules (three glowing circles) and graphene. **(B)** The bending of DNA leads to a change in the distance between the attached fluorescent molecule and graphene. **(C)** Sketches of the three exemplary modifications to the investigated DNA representing a variation in salt concentration, introduction of a gap in one of the DNA strands or the application to sense DNA mismatches (from top to bottom).

In this work, we combine GETvNA with autocorrelation analysis to monitor the rapid, sub-microsecond motion of dsDNA tethered to graphene. Nucleic acids are immobilized on graphene by employing a hybrid DNA system containing a dsDNA segment and a single-stranded DNA (ssDNA) overhang. We clearly distinguish conformational dynamics from photophysical intensity fluctuations using a combination of time-correlated single-photon counting and shrinking-gate fluorescence correlation spectroscopy (sgFCS).^29^ This reveals how ionic strength influences the mechanical properties of DNA on the sub-microsecond timescale. Secondly, we study several structural variants including dsDNA with gaps of 1-3 nucleotides and dsDNA with 1-3 successive mismatches (Figure 1C). The results are rationalized with complementary molecular dynamics (MD) simulations providing an atomistic view of the fluctuation changes induced by the variation of ionic strength or gaps in dsDNA. The developed experimental tools and analysis open a new window to visualize subtle biomolecular dynamics bridging the long-existing timescale gap between atomistic simulations and experimental approaches at room temperature.^30^

## Materials and Methods

Unless explicitly mentioned, laboratory products were purchased from Carl Roth and chemicals were purchased from Sigma Aldrich. Dye-modified DNA strands were purchased HPLC-purified from Ella Biotech GmbH, and unmodified DNA strands were procured from Integrated DNA Technologies, Inc.

### Graphene Preparation

60 mm × 40 mm monolayer graphene on a copper substrate with an overlying poly(methyl methacrylate) (PMMA) layer was purchased from ACS Materials LLC, USA or Graphenea Inc., Spain. The graphene foil was stored in a vacuum-sealed desiccator. All coverslips were cleaned in a 1% v/v Hellmanex™ III (Hellma GmbH & Co. KG, Germany) solution for 15 min with ultrasound at 37 °C and additional three rounds of washing for 15 min with MilliQ water were performed.

Graphene was cut from the copper/graphene/PMMA foil into pieces of ∼5 mm and placed on the surface of a 0.2 M aqueous ammonium persulfate solution for four hours to etch the copper away. Using cleaned coverslips, the floating PMMA/graphene film was scooped from the solution and transferred to a bath of Milli-Q water to remove residual etchant. To remove remaining ammonium persulfate, the foil was washed twice with MilliQ water.^31,32^ The PMMA/graphene coverslip was then dried with nitrogen. To cure the graphene, the PMMA is temporarily dissolved in a PMMA-containing chlorobenzene solution and the solvent is evaporated after 1 h.^33^ The resulting PMMA coating on the graphene monolayer is removed by two baths of the entire sample in acetone for 10 min. To remove any remaining PMMA residue, the coverslip was placed in clean toluene for 10 min. The solvent was evaporated with a nitrogen stream, and the sample was heated on activated carbon and placed on a heating plate at 230 °C for 30 min. The cooled coverslip with graphene monolayer was used for the following immobilizations. For a graphical overview of the whole process, the reader is referred to supplementary figure S1 in ref. ^23^.

### DNA Preparation

ssDNA strands exceeding 50 nucleotides in length were purified on an 8 M urea polyacrylamide gel (12% v/v acrylamide, 90 mM Tris-HCl, 90 mM boric acid) for 1 h (100 V, 15-8 mA) using 1× TBE buffer (90 mM Tris-HCl, 90 mM boric acid, 1.1 mM EDTA). Post-electrophoresis, the DNA was visualized without further staining by exposing to UV light at 254 nm (Analytik Jena GmbH + Co. KG, Germany) or a blue LED transilluminator (IO Rodeo, USA) in 400 μL elution buffer, and frozen at –20 °C. The solution was agitated for 3 h, and the gel debris was separated using a Spin-X filter (FavorPrep™ GEL/ PCR Purification, Favorgen Biotech Corp., Austria) for 2 min at 10,000 g, room temperature. Subsequently, 100 μL elution buffer (500 mM ammonium acetate, 10 mM magnesium acetate) was added to the gel debris and centrifuged again at 10,000 g for 2 min. The DNA strands were desalted through dialysis. The sample was concentrated using a 3K Amicon Ultra Filter (Merck Millipore, Merck KGaA, Germany) to a concentration of 1.0 μM or higher. DNA quantification was performed by measuring UV absorbance at 260 nm or the absorbance of the specific dye with a photospectrometer (Thermo Scientific NanoDrop 2000, Thermo Fisher Scientific Inc., USA). Samples were loaded onto a polyacrylamide gel in 1×TBE buffer (60 min at 100 V) and stained with Sybr™Gold (Invitrogen, Thermo Fisher Scientific Inc., USA) for the final visualization of the purified DNA strand. Throughout the entire purification process, ssDNA strands labeled with a fluorescent dye were handled separately from those without.

For GETvNA experiments, complementary ssDNA strands were annealed in 1×TAE buffer (40 mM Tris-HCl, 20 mM acetic acid, 1 mM EDTA, 500 mM sodium chloride). The mixing process involved a 10× excess of the longer ssDNA strand, which served as a toehold to bind to graphene. Following premixing, the solution was agitated at 300 rpm and 37 °C on a thermomixer (Eppendorf ThermoMixer® C, Germany) for 2 h, thereby inducing the formation of dsDNA. To ensure complete hybridization, hybrid DNA constructs were always assembled at 500 mM sodium chloride concentration. The variation of the sodium chloride concentration was realized in the subsequent step of GETvNA sample preparation.

### Preparation of DNA Origami Nanostructures

DNA origami nanostructures were designed using caDNAno^34^, employing the p8064 scaffold derived from M13mp18 bacteriophages (staple sequences as in Supplementary Table S5). The folding was performed in folding buffer (40 mM Tris-HCl, 20 mM acetic acid, 1 mM EDTA, 19 mM magnesium chloride, 5 mM sodium chloride) using a 10-fold molar excess of unmodified and labeled oligonucleotides relative to the scaffold strand. The thermal annealing program used is listed in Supplementary Table S6.^35^ After folding, 1× Blue Juice gel loading buffer (peqGREEN (VWR)) was added to the DNA origami solution, followed by purification through ice cooled agarose-gel electrophoresis with 1.5 % agarose gel in 50 mL of 1× TAE buffer (with 12.5 mM magnesium chloride) at 70 V for 2 h. The band corresponding to the nanostructure was excised from the gel. The purified DNA origami solution was adjusted to a final concentration of 50 pM in 1× TAE buffer and incubated on the graphene surface for 1 min. The surface was subsequently washed three times with 1× TAE buffer, after which 147 μL solution of Tx/Tq-PCA buffer (20 mM acetic acid, 1 mM EDTA, 100 mM sodium chloride, 2 mM trolox/troloxquinone (pre-dissolved in 0.5 mM methanol), 12 mM protocatechuic acid (PCA)) and 3 μL 50×PCD (2.8 mM Protocatechuate 3,4-Dioxygenase from *Pseudomonas* sp. (PCD, 25U/mL), 50 % v/v glycerol, 50 mM KCl, 100 mM Tris-HCl, 1 mM EDTA) buffer was mixed and added to the sample. The entire chamber was then hermetically sealed.

### GETvNA Sample Preparation

Subsequent to the preparation of DNA or DNA origami nanostructures, GETvNA samples were prepared. A chamber (150 μL volume, SecureSeal™ hybridization chambers, Grace Bio-Labs, USA) was attached around the graphene sheet on the glass coverslip. The graphene surface was incubated with the annealed DNA oligonucleotides around 100 pM for 1 min. The chamber was carefully washed three times with 1× TAE buffer. To achieve photostabilization by removing oxygen and suppressing triplet states, a 147 μL solution of Tx/Tq-PCA buffer (20 mM acetic acid, 1 mM EDTA, 100 mM sodium chloride, 2 mM trolox/troloxquinone (pre-dissolved in 0.5 mM methanol), 12 mM protocatechuic acid (PCA)) and 3 μL 50× PCD (2.8 mM 3,4-Dioxygenase from *Pseudomonas* sp. (PCD, 25U/mL), 50 % v/v glycerol, 50 mM KCl, 100 mM Tris-HCl, 1 mM EDTA) buffer was mixed and added to the sample. The potential variation of the sodium chloride concentration was realized by washing the sample 5× with 1× TAE buffer but adjusted concentration. The buffer for photostabilization was concentration-adjusted, too. The entire chamber was then hermetically sealed with Microtube Tough-Spots® labels or equivalent adhesive labels.^23^

### Optical Setup

The single-molecule experiments were carried out on the commercial confocal microscope Luminosa (PicoQuant GmbH, Figure 2A). In transmission mode, the area with graphene was identified on the microscope slide and used for the following measurement. The fluorescent molecules were excited using a pulsed laser at a wavelength of 640 nm, operated at 80 MHz with an excitation power of 4 μW. The laser light was coupled into the epi-illuminated confocal microscope via a dichroic beam splitter (DB532/640) and focused into the sample by an oil immersion objective (100x, NA 1.45, UPLXAPO, Olympus). The emitted fluorescence was collected through the same objective and spatially filtered using a pinhole with a diameter of 50 μm. The fluorescence signal was divided into two detection channels by a non-polarizing, 50:50 beam splitter (BS 50/50). The fluorescence was filtered in both channels by a bandpass filter (BP690/70) and focused on two separate single-photon counting modules. Immobilized molecules were located by performing a surface scan with the piezo-scanner. Molecules were automatically selected using PicoQuant’s Luminosa software, and fluorescence traces were recorded for two minutes or until irreversible photobleaching of one dye occurred.

**Figure 2.**
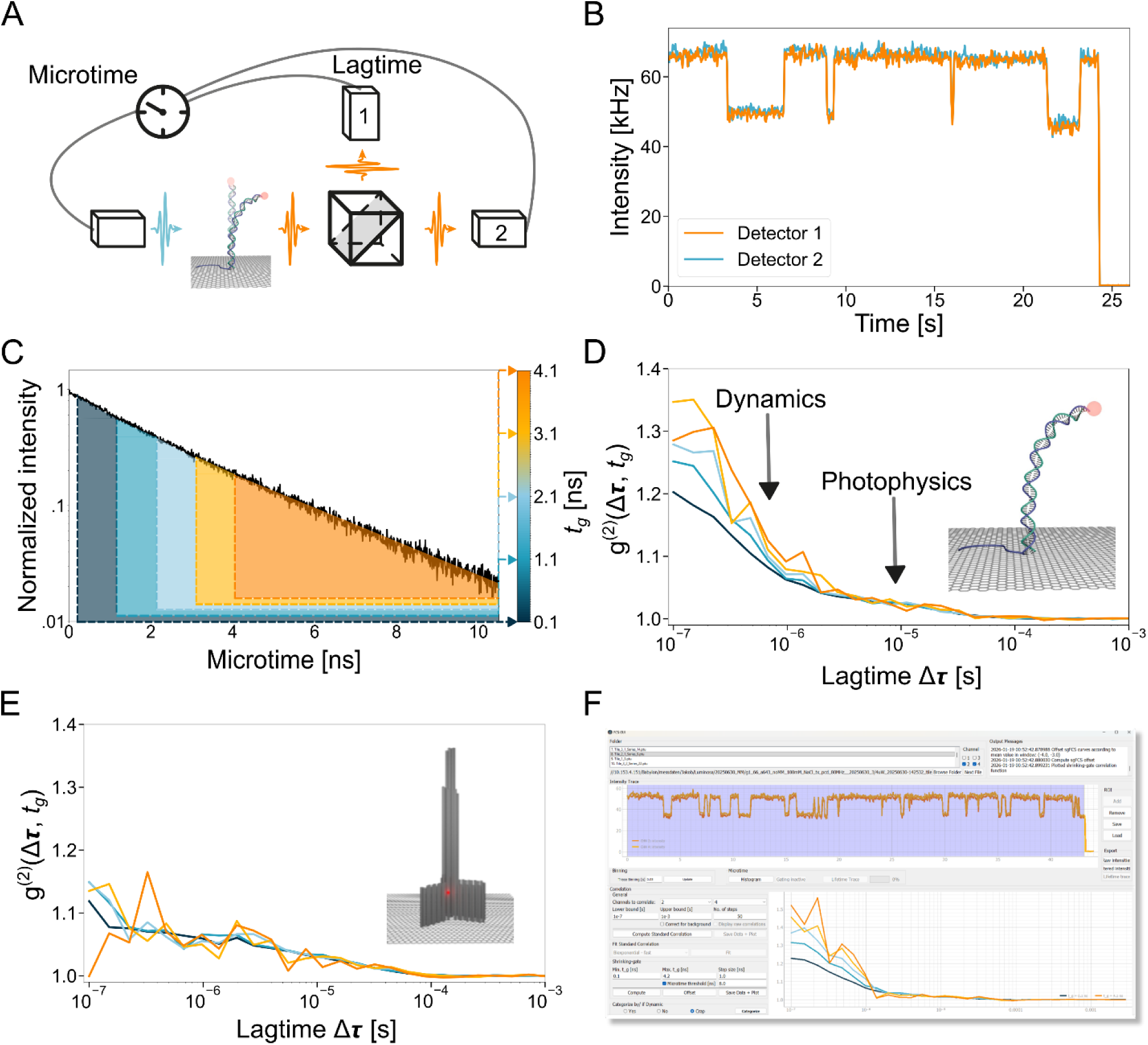
Measurement of sub-microsecond fluctuations in DNA using single-molecule fluorescence. **(A)** Sketch depicting the experimental setup based on pulsed laser excitation of the sample and the detection of the emitted fluorescence photons by two avalanche photodiodes after splitting the signal using a non-polarizing beamsplitter. The photon arrival times are cross-correlated to overcome the dead time of the detectors and monitor photon bunching for lagtimes Δ_τ_ in the sub-microsecond regime. **(B)** Exemplary intensity-time trace from a single fluorescent molecule with a single bleaching step at 24 seconds. The acquired photon counts at the detectors 1 (orange) and 2 (blue) are binned at 50 ms. Due to the high-power as well as pulsed excitation the majority of the time-intensity traces exhibit well-known spectral shifts on the second time scale leading to a temporarily reduced fluorescence intensity (Supplementary Figures S1-S10). **(C)** Fluorescence decay histogram from detector 1. Overlaid to the decay are five exemplary, color-coded areas each representing a microtime threshold according to the color bar on the right and indicated by the respective labels. **(D)** Resulting intensity cross-correlation of the fluorescence signals in (B). As in (C), the microtime thresholds are color-coded from dark blue (0.1 ns) to orange (4.1 ns). Inset: rendering of bent dsDNA being immobilized on graphene. **(E)** Exemplary correlation function for the same parameters as in (B) for the fluorescence signal from a DNA origami structure as depicted in the inset. The absence of a microtime-dependent amplitude exemplifies the fixed distance of the fluorescent molecule in the rigid DNA origami structure. As in (C), the corresponding microtime thresholds are color-coded from dark blue (0.1 ns) to orange (4.1 ns). **(F)** Publicly available user interface for graphical analysis of single-molecule fluorescence signals (https://gitlab.lrz.de/tinnefeldlab/fcs-gui).

### Correlation Analysis

The respective second-order intensity correlations are time-binned according to a multi-tau photon correlation algorithm.^36^ Correlations from two detectors, *e*.*g*. 1 and 2, are calculated by averaging *g*^*(2)*^*(1*⊗*2)* and *g*^*(2)*^*(2*⊗*1)*. To reveal lifetime correlated intensity fluctuations, we used the shrinking-gate fluorescence correlation spectroscopy (sgFCS) algorithm.^37^ Here, macrotime stamps were filtered according to their photon arrival time following pulsed laser excitation. We marked the maximum of the TCSPC histogram as 0 ns. The applied microtime thresholds range from 0.1 ns to 3.1 ns and are applied at step sizes of 0.5 ns. Here, the lower limit of 0.1 ns was chosen in order to discard scattered photons from the excitation laser pulse. Furthermore, a cut-off threshold of 8 ns was used for the microtime to increase the signal-to-noise ratio. Additionally, the visibility was enhanced by offsetting the microtime-thresholded correlation functions to a common value. This removes long timescale correlations related to common spectral jumps of the dyes. The shrinking-gate correlation from one measurement was visually inspected. The dataset was classified as ‘dynamic’, if the microtime thresholding leads to an increasing correlation amplitude with respect to smaller thresholds for at least four consecutive data points.

Once dynamics are identified, the respective correlation function using all photons was computed. For this, a background correction is performed according to ^29^. The data is fitted in the interval between 100 ns and 3 ms using a biexponential decay model,

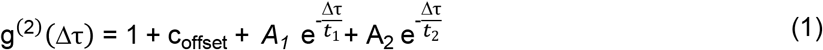

with a variable offset parameter c_offset_ and correlation times *t*_*1*_ and *t*_*2*_. Depending on the visibility of the respective dynamics and to avoid fitting noise, the start of the fit is adjusted accordingly, but never exceeds 300 ns. The fit is carried out using a least-squares algorithm, yielding the corresponding correlation amplitude values *A*_*1,2*_ and correlation times *t*_*1,2*_. In our experiments, the fit shows microtime-independent photon bunching with correlation time on the order of 10 μs, that we assign to triplet state formation, and a dynamic component with a correlation time on the order of 0.5 µs correlation time. For the dynamic component, we drop the subscript for the correlation amplitude and write *t*_*c*_ instead of *t*_*2*_ for the correlation time.

### Molecular Dynamics Simulations

All simulations were carried out with the Amber24^38^ software package employing the bsc1 DNA forcefield^39^ together with the TIP4P-FB^40^ water model and the corresponding 12-6-4 Lennard-Jones parameters^41^ for sodium and chloride ions. Fully solvated DNA systems and a DNA-graphene system were considered. The solvated DNA starting structures were generated with the NAB and LEaP software included in AmberTools24^38^. Two DNA strands were considered, one 36 base pair DNA duplex with varying counter ion concentrations, and one 36 base pair DNA duplex with two nucleotides missing in positions 17 and 18 (3’-5’) together with its mismatch-free counterpart. A summary of all systems investigated is given in the Supplementary Information in Table S3. The 2 nt-gap system was built analogously to the mismatch-free DNA system by deleting the missing bases and adding terminal hydrogens *via* the LEaP program. For simplicity, single strand parts were omitted from the simulations. Analysis and visualization of the trajectories was performed with cpptraj^38^, pytraj^38^ and VMD^42^. The three graphene sheets (15 x 15 nm) in the DNA-graphene system were built using the VMD nanotube plugin with the carbon atoms modeled as ca-type atoms of the gaff force field (gaff2-ca).^43^

The DNA-water simulations were set up in a truncated octahedron box with a minimum buffer zone of 2 nm. Sodium cations were added to the system to reach 50 mM (1 equivalent [eq.], relative to the number of DNA phosphate groups), 250 mM (5 eq.) and 550 mM (11 eq.) Na^+^ concentrations. The DNA-graphene-water system was constructed in an orthogonal box with a minimum buffer zone of 1 and 2 nm (in the *z* direction), respectively. To achieve a similar neutralization ratio (52%) within 0.7 nm of the DNA surface, we used a 25 mM sodium solution in the DNA-graphene-water system, which provides the same effective Na^+^ equivalence as the 50 mM Na^+^ concentration in the DNA-water system. The 2 nt-gap system and its mismatch-free counterpart were submerged in 100 mM (2 eq.) Na^+^. Chloride ions were added to the systems to reach charge neutrality.

A cut-off criterion of 1 nm was imposed on the Lennard-Jones interactions. Long-range Coulomb interactions were treated with the Particle Mesh Ewald Summation^44,45^ method with a real space cut-off of 1 nm. A time step of 2 fs was used for the integration of motions while all bonds including hydrogen and all water molecules were constrained with the SHAKE^46^ and SETTLE^47^ algorithms respectively. Coordinates were saved every 10 ps.

All systems were optimized in a two-step approach by first applying harmonic positional constraints of 500 kcal mol^-1^ Å^-2^ on the solute atoms to allow relaxation of the solvent (5000 steps) followed by a constraint free second optimization (10000 steps). After half of the respective optimization steps, the algorithm was switched from the steepest descent to the conjugated gradient method.

The systems were heated to 300 K over a period of 400 ps using the Langevin thermostat^48^ employing a collision frequency of 1 ps^-1^ while keeping all solute atoms fixed with harmonic constraints of 10 kcal mol^-1^ Å^-2^. Due to the inherent instability of the 2 nt-gap system, harmonic restraints of 100 kcal mol^-1^ Å^-2^ were applied to the solute atoms over 400 ps and removed over the following 5 ns, this procedure was also applied to the corresponding mismatch-free DNA sequence for consistency (see Supplementary Information, Supplementary Table S1 for sequence information). Subsequent flexible equilibration in the NpT ensemble at 1 bar was performed for at least 500 ns. The pressure was controlled with the Berendsen barostat^49^ using a pressure coupling constant of 2 ps. Graphene was constrained throughout all MD steps to its initial positions with harmonic constraints of 1 kcal mol^-1^ Å^-2^. Production sampling runs cover a timescale up to 3 μs, depending on system size (see Supplementary Information & Supplementary Table S3).

After initial equilibration, a positional restraint of 100 kcal mol^-1^ Å^-2^ was imposed on the heavy atoms of the first two base pairs to mimic the immobilization of the DNA on a virtual graphene sheet with subsequent re-equilibration of at least 500 ns. The effect of immobilization was later compared to the simulated dynamics of the DNA-graphene-water system (Supplementary Information, Supplementary Figure S16 & Supplementary Table S7).

### Kinetic Monte Carlo Simulations

Kinetic Monte Carlo simulations are based on a probability distribution that is used as an input. This distribution describes the probability of residence of a single photon emitter being connected to the end of dsDNA that is pointing away from graphene. For a polymer of contour length *b* and persistence length *l*_*p*_, we assume that it is well described by the Worm-like chain (WLC) model. Further, we assume that in GETvNA, the polymer stands perpendicular on graphene at the single-to-double stranded junction (that is the nucleobases closest to graphene). If unbent, the measured graphene distance *z* of the polymer will equal the contour length *b* plus 0.4 nm offset since the dsDNA is not in direct contact with graphene. Assuming a continuous bending along the contour length, each bending angle *θ* (defined as in ^26^) relates to a unique graphene distance *z* that is in quadratic approximation: *θ* = sqrt(6(1-*z/b*)). For contour lengths smaller than the persistence length,^50^ each bending at angle *θ* means a bending energy *U*_*B*_ = 0.5 κ_*B*_ / *b* * *θ*^*2*^ with the bending stiffness κ_*B*_. The first quadratic approximation is valid up to bending angles of *θ ≤* 75° and allows equating κ_*B*_ *= l*_*p*_ *\* k*_*B*_ *\* T*. Based on the assumption that the unbent state has a probability that is equal to unity, we calculate a probability distribution according to Boltzmann as *p(θ) = exp(-0*.*5 * l*_*p*_ */ b * θ*^*2*^).

The above consideration relates a probability *p* to a certain bending angle *θ* and with this to a graphene distance *z*. However, in GETvNA for each bending angle *θ*, there are several possible torsional configurations that lead to the same measured graphene distance *z*. We model this entropic component by calculating the accessible circumference for a dsDNA with the corresponding bending angle *θ*. Based on the ascribed enthalpic and entropic components, we found a probability for each graphene distance that only depends on the contour length of the dsDNA and its persistence length. Following, we performed kinetic Monte Carlo simulations. Here, we assume that the single-photon emitter moves according to the derived probability distribution and is described by normal diffusion with a diffusion constant *D*. Further, assuming this single-photon emitter is excited at a given repetition rate *k*_*rep*_ and emits photons depending on its current graphene distance, and given a certain quantum yield as well as fluorescence lifetime. Diffusion and photon emission are interconnected by the underlying probability of diffusion *p*_*Diff*_ *= 2 * D / dD*^*2*^ */ k*_*rep*_ with *dD* defining the diffusion distance (equal to the mean square displacement) during one laser cycle.

We repeated the simulation for a given set of parameters at least three times for simulated ‘experiment durations’ of 2.5 s to 20 s. The obtained photon macro- and microtime stamps were analyzed for dynamics, equivalent to the experimental data. However, since our simulation neglects photophysical processes such as, for example, triplet transitions, we fitted the data for times larger than 0.1 µs using a single exponential fit in combination with a least-squares fitting algorithm.

## Results

### Measuring sub-microsecond fluctuations in DNA

In our experiments, we immobilized hybrid DNA constructs consisting of a single- and double-stranded part on graphene.^26^ The ssDNA adsorbs on graphene, whereas the dsDNA stands perpendicular. Here, the ssDNA was 30 nucleotides (nt) long and the dsDNA 66 nt, except for the experiment presented in Figure 4 (see Supplementary Table S1 for sequences). To maintain the advantage of single-molecule detection, that is the ability to characterize subpopulations, we aimed to quantify the conformational dynamics from individual molecules. With expected correlation times down to the nanosecond range, this required high count rates and long single-molecule survival times before photobleaching.^51,52^ To this end, we tagged the end of the dsDNA with a photostable and hydrophilic ATTO 643 molecule.^53^ We employed comparatively high excitation intensity of 4 µW at 80 MHz repetition rate (measured in front of the dichroic mirror), yielding intensity traces with count rates of 100-160 kHz over a few to several tens of seconds that often exhibited reversible, intensity jumps on the second time scale that we assigned to spectral jumps (Figure 2B), a known phenomenon for carborhodamine dyes such as ATTO 643 and ATTO 647N (Supplementary Figures S1-S10).^53^ The spectral jumps led to a change in fluorescence lifetime of less than 10% and related autocorrelation amplitudes were filtered out by subtracting long timescale amplitudes.

We used second-order correlation analysis to detect fluctuations in intensity. In our specific case, we made use of the second-order intensity correlation g^(2)^,

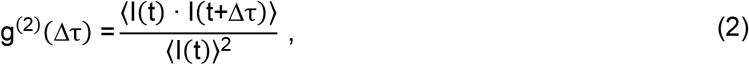

which correlates the measured fluorescence intensity *I(t)* at time *t* with the measured intensity after the additional lagtime *Δ*τ. Experimentally, the correlation of single photons is compromised by electronic artefacts from after-pulsing or the dead time of the detector. They are circumvented by introducing a non-polarizing, 50:50 beamsplitter in the detection path that splits the signal equally on two detectors.^54^ The corresponding cross-correlation of the intensities measured at both detectors, thereby, is no longer subject to electronic artefacts of a single detector and the g^(2)^ correlation reveals intensity fluctuations down to the time scale of the fluorescence lifetime, where photon antibunching occurs (Figure 2A). Using time-correlated single photon counting allows for assigning to each photon detection event an absolute time with respect to the beginning of the experimental acquisition, the so-called macrotime (Figure 2B), and a relative time that elapsed since the last laser pulse, *i*.*e*. the microtime. Further, the monoexponential fit of the detected and histogrammed microtimes yielded the corresponding fluorescence lifetime (Figure 2C).

Correlation analysis revealed different correlation amplitudes that can represent conformational dynamics of interest but also photophysical processes such as triplet or redox blinking. In our case, conformational dynamics were associated with changes of the fluorescence lifetime due to dynamic distance variations between the dye and the graphene surface. In contrast, on-off blinking e.g. induced by triplet state transition that typically occurs on the microsecond timescale has no influence on the fluorescence lifetime. Analysis of the intensity correlation only showed the time scale of the underlying fluctuations but not if the fluorescence lifetime is affected correlated to the intensity fluctuation. sgFCS allows to separate intensity fluctuations that correlate with a change in fluorescence lifetime.^37^ sgFCS uses multiple thresholds *t*_*g*_ to filter the macrotime stamps by the respective microtime. The filtering by *t*_*g*_ values increases the intensity contrast for fluorescence lifetime-dependent intensity fluctuations and, thus, the corresponding correlation amplitude. Importantly, photophysical processes like triplet blinking do not alter the fluorescence lifetime and the bunching amplitude of this process stays constant for any *t*_*g*_. Five exemplary thresholds (*t*_*g*_ = 0.1, 1.1, 2.1, 3.1 & 4.1 ns) represented by dashed lines and colored areas are shown in dark blue to orange in Figure 2C. In Figure 2D, the corresponding cross-correlations are depicted according to the same color code. The autocorrelation with the smallest threshold of 0.1 ns essentially represents the common correlation function with two visible correlation amplitudes at ∼0.1 µs and ∼70 µs. Interestingly, the increased thresholds yielded larger correlation amplitudes for the short component whereas the long component stayed constant. We accordingly assigned the short amplitude to originate from biomolecular dynamics whereas the long component was related to on-off processes such as triplet state formation.

The correlation amplitude rises for larger thresholds, indicating fluorescence intensity fluctuations that correlate with changes in the lifetime on the time scale of 100 ns. Biexponential fits are used to extract the correlation amplitudes and times for quantifying the dynamic processes (see Methods section for details). For comparison, we immobilized an ATTO 643 at a comparable height above graphene in a rigid DNA origami structure that prevents similar biomolecular dynamics. Corresponding sgFCS analysis of fluorescence intensity traces only exhibits the photophysical component of ∼ 70 µs correlation time and a lack of the faster threshold dependent component, supporting our hypothesis that thermal motion of dsDNA can be studied on single molecules using GETvNA (Figure 2E, Supplementary Figure S1).

For analysis of the acquired fluorescence transients, a home-written user interface was employed for screening and categorizing the acquired single-molecule data efficiently (Figure 2F, open-source software written in Python, available at https://gitlab.lrz.de/tinnefeldlab/fcs-gui). Among 159 molecules of the 66 bp-long dsDNA sample at 100 mM sodium chloride concentration 46% showed the dynamics as exemplified in Supplementary Figures S2-S8 whereas for the remaining 54% an insufficient number of fluorescence photons or count rate, respectively, prevented a clear characterization with quantitative fitting (Supplementary Figures S9 & S10). The analysis of the dsDNA reveals a homogeneous distribution of the fast correlation component with a mean correlation time of (0.44 ± 0.02) µs and an amplitude of 0.22 ± 0.01 (Supplementary Figure S11). For the reference DNA origami structure, this short component is not observed at all.

Being able to unveil rapid dsDNA dynamics, we first studied the effect of different ion concentrations on the fluctuation strength and time scale (Figure 3A). We recorded single-molecule intensity traces and fitted the fluorescence lifetime decays obtained from all photons by a monoexponential decay model with reconvolution of the instrument response function, providing a fluorescence lifetime τ. This value was converted to a graphene distance *z* according to the equation,

**Figure 3.**
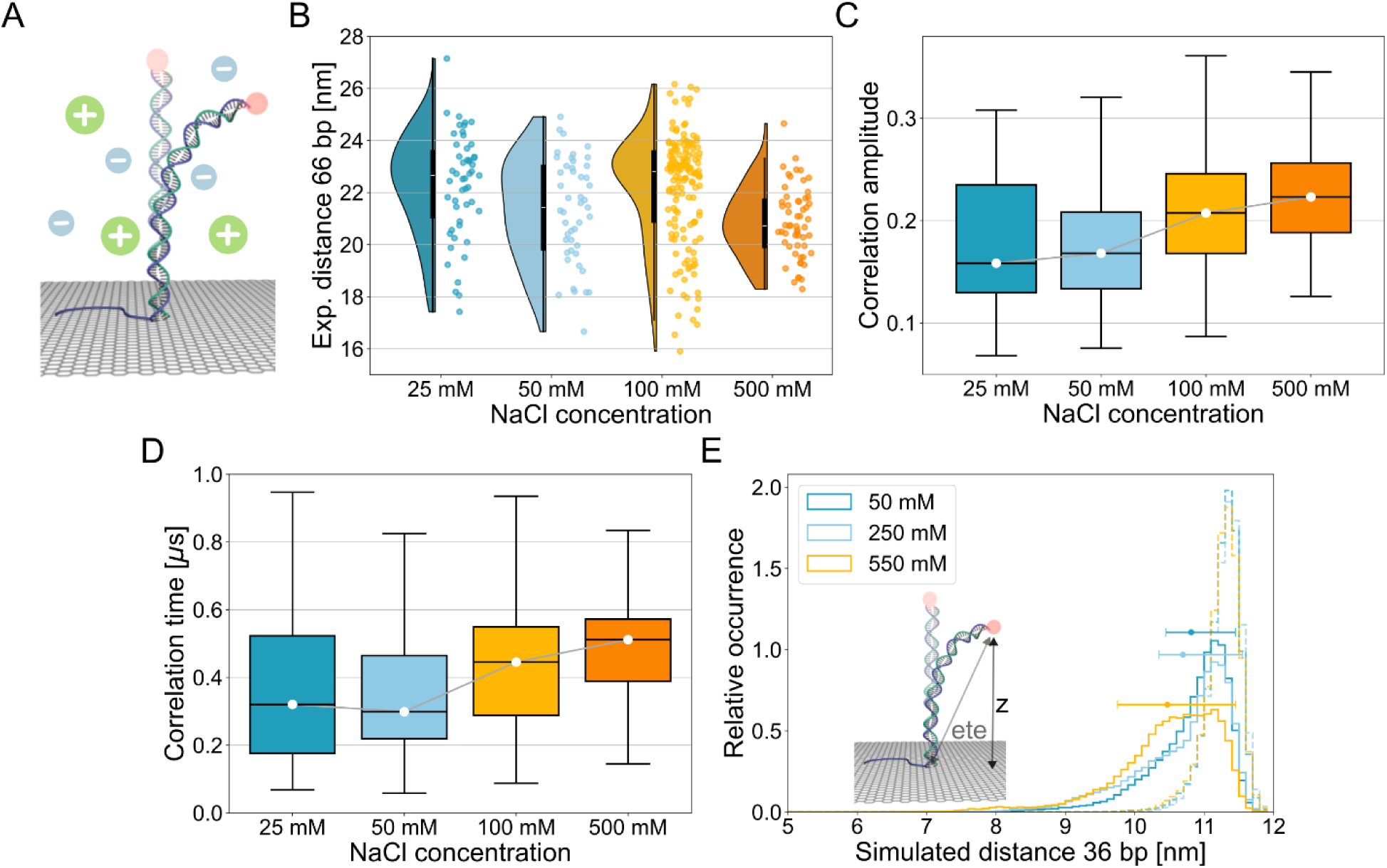
Ion-dependent fluctuation dynamics of dsDNA. **(A)** Schematic of the GETvNA sample. **(B)** Experimentally determined values of the graphene distance derived from the fluorescence lifetime for a 66 bp long dsDNA construct at four sodium chloride concentrations (*n* = 52, 47, 155, 57). These results correspond to DNA molecules exhibiting dynamics in the sgFCS. Within the violinplot, the white dot indicates the median, the box marks the first to third quartile and the whiskers range from minimum to maximum, excluding outliers. **(C)** Boxplot of the experimentally determined correlation amplitudes. **(D)** Corresponding correlation times. The box plots in (C) and (D) display each distribution excluding outliers with the whiskers ranging from minimum to maximum. The respective boxes visualize the first and third quartile separated by the median. The median values are connected by a grey line to guide the eye. Individual data points including outliers are depicted in Supplementary Figures S13-S15. **(E)** Histograms of *in silico* data from 3 µs-long MD simulations for a 36 bp dsDNA strand at the indicated sodium concentrations. The graphs show the distances from the *z*-projection in solid whereas the transparent, dashed lines correspond to the histograms of the respective end-to-end distances (ete; see Supplementary Figure S22 for details). The colors encode the respective ion concentration as indicated in the legend. The dots and error bars represent the mean distance and fluctuation range, measured by the distance that each distribution has a value that is larger than *1/e* times its maximum value.

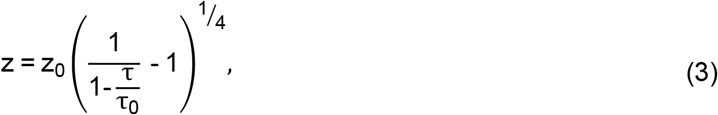

with the distance of 50% energy transfer efficiency, *z*_*0*_, as well as the fluorescence lifetime in the absence of graphene, τ_0_ = 3.91 ns (Supplementary Figure S12). The calculated graphene distances of all molecules exhibiting a clear threshold-dependent correlation amplitude are presented in Figure 3B, indicating that higher ion concentrations lead to increased height fluctuations and, on average, smaller graphene distances.^55^ The mean distances and their standard error are (22.3 ± 0.3) nm, (21.3 ± 0.3) nm, (22.2 ± 0.2) nm and (20.9 ± 0.2) nm for 25, 50, 100, and 500 mM sodium chloride, respectively. Comparing the data of molecules with threshold-dependent correlation amplitude to those without (Supplementary Figure S13) exemplifies that the attribution of dynamics according to the sgFCS does not act strictly as a filter for specific graphene distances.

In Figure 3C, the experimentally measured correlation amplitudes are sketched. We find mean values and standard errors of 0.19 ± 0.02, 0.17 ± 0.01, 0.22 ± 0.01 and 0.23 ± 0.01 for the four samples ranging from 25 mM to 500 mM, indicating stronger intensity fluctuations enabled by higher salt concentrations which shield the negative backbone charges more efficiently and, thus, cause a shorter persistence length.^56,57^ For salt concentrations higher than 50 mM, the measured correlation times display a similar trend to the amplitudes (Figure 3D) with mean value and standard error of (0.37 ± 0.04) µs, (0.36 ± 0.03) µs, (0.44 ± 0.02) µs and (0.51 ± 0.03) µs. This indicates that the fluctuation speed decreases and suggests a stabilization of bent states by the cations, in agreement with a previous report.^58^

To rationalize our idea of visualizing thermal dsDNA fluctuations, we performed molecular dynamics (MD) simulations. To ensure an efficient use of the available computational resources, the MD simulations were realized for a 36 bp-long dsDNA with the same sequence as the dye-facing dsDNA part in the experiment (Figure 3E). First, we found that the effect of the graphene immobilization on the intrinsic DNA structure is negligible. We observed only 1.6% deviation for the projected elongation of the DNA to the graphene plane in comparison to the DNA without graphene (but with restrained ending base pairs) or reference plane, respectively, which is smaller than the associated error (see Section 1.1 in Supplementary Information and Supplementary Figure S16). For this reason, simulations for varying ion concentrations were carried out with restrained ending base pairs but without explicit graphene yet requiring an adjusted definition of the *z*-projection of the DNA (Supplementary Figure S17-S19).

Interestingly, the MD simulation of the 36 bp dsDNA and the experimentally employed 66 bp dsDNA show a similar dependency of the projected dye-graphene distance on the ion concentration (Supplementary Figure S20). As annotated in Figure 3E, the mean value and fluctuation range (measured by the distance that each distribution has a value that is larger than *1/e* of its maximum value) reveal a decrease in distance but an increase in the fluctuation range with a larger number of ions in the simulation volume. Further, the simulations establish that this effect results purely from the increased flexibility of DNA since there is no change observed in the average helix rise or DNA groove width (Supplementary Table S2). The findings are in agreement with earlier reports showing a decrease in persistence length with ion concentration.^59,60^ Further, the simulation results are in line with the experimental data, which show a decreased height fluctuation time in the autocorrelation analysis of the simulated trajectories (Supplementary Figure S21).

Importantly, the MD simulations enable us to compare the graphene distance that is accessible in our experiments to the end-to-end distance that would have been determined in a corresponding FRET experiment (Figure 3E). The main geometric difference is that GET measures the projection to the surface, whereas FRET would measure the end-to-end distance of the DNA. Strikingly, for DNA bending, measuring the GET projection is more sensitive to fluctuations at different salt concentrations and the distance changes are more pronounced. Therefore, GETvNA might become the method of choice to study fluctuations in B-DNA (Supplementary Figure S22).^61–63^

As molecular dynamics simulations were carried out with shorter DNA than the experimental one, we faced the challenge of directly comparing experimental sgFCS results and molecular dynamics simulations. For translating the end-to-graphene distribution from the 36 bp molecular dynamics simulation to the 66 bp experimental DNA, we fitted the persistence and contour length to optimally match the MD distance distribution (Supplementary Figures S23 & S24). Based on the resulting parameters, we calculated the expected distance distribution for the 66 bp DNA using the WLC model. Furthermore, the timescales of the MD simulations are similar to the time scales of the biomolecular dynamics but much shorter than the total duration of the time traces used for correlation analysis. To bridge molecular dynamics simulations with experiments, we emulated the experiment with kinetic Monte Carlo simulations. In our simplified model, we assumed that DNA bending occurs equally throughout the molecule; that is, the DNA-end moves at a constant speed of 0.6 nm per time step. The time steps were adapted to match experimental diffusion constants from earlier measurements.^64,65^ The directionality of the movement was biased to obtain the distance distribution of the 66 nt DNA. At each time point, there is a probability of photon detection adjusted to match realistic average photon streams. With each photon detection event, the fluorescence lifetime information reflecting the graphene distance was recorded.

The resulting photon stream of the Monte Carlo simulations is analyzed in the same manner as the experimental data and, hence, provides an estimate for the expected fluorescence correlation amplitude as well as the fluctuation time scale. For a 66 bp-long dsDNA with a diffusion constant of *D* = 10 µm^2^/s and 50 nm persistence length, the simulation yields a correlation amplitude of 0.10 ± 0.01 and a correlation time of (1.56 ± 0.22) µs (Supplementary Figures S25 & S26). Both values are in reasonable agreement with the experimental values showing that the simple analytical model can rationalize our experimental data (Supplementary Figure S11). However, we emphasize that the WLC model is an ideal model and that dsDNA with short contour lengths, partially, might not follow this model.^5^ Hence, the simulated correlation time might be higher than the experimental result, potentially, because the dsDNA fluctuates faster than was previously reported, and assumed in the simulation. Furthermore, the simulation neglects influences, as for example, due to the present fluid dynamics or the graphene surface.^66–68^ Additionally, the simulated amplitude value might be lower than measured if few but strong bending events occur as suggested by some models.^69^ Yet, the analyses indicate that our method allows for unraveling the rapid, thermally induced bending of dsDNA by combining sgFCS with GETvNA.

We close the analyses of the influence of the ion concentration by combining evidence from the experiments as well as the atomistic MD simulations. The fluctuating dsDNA that was immobilized on graphene undergoes confined diffusion. The MD simulation shows a decrease in the respective diffusion coefficient when analyzing the dynamics on the order of ten picoseconds time delay compared to a time scale of ten nanoseconds indicating that the motion of the tethered DNA tends towards confined diffusion on longer time scales (Supplementary Figure S27). This effect is reduced for higher concentrations since the cations effectively shield the negatively charged phosphate backbone of the DNA, allowing for more flexibility (Supplementary Figure S28 & S29). Thus, more ions mean stronger bending of the DNA leading to an increased entropic contribution, hence there is more randomness in the motion of the system. For this, the diffusion will be less confined. This trend is reflected in the results from the MD simulations and, further, in the experimental results showing a decreasing anticorrelation between correlation amplitude and correlation time for increasing ion concentrations (Supplementary Figure S30). Again, as the MD simulation showed, this subtle concentration dependency becomes only experimentally accessible thanks to the unique geometry underlying GETvNA.

Introducing gaps of different sizes to the dsDNA induces increased fluctuations to the system as proven by nuclear magnetic resonance or gel electrophoresis experiments.^70,71^ Previously, birefringence experiments demonstrated that for dsDNA with a length of less than 101 bp, these dynamics lie in the microsecond regime.^72^ To examine the impact of these dynamics under our experimental conditions, we next studied DNA constructs containing missing bases (gaps) and analyze their fluctuation behavior (Figure 4A). In the experiment, the employed dsDNA that is 80 bp-long with a gap at 17 nt counting from graphene, and a 40 nt ssDNA. We switched to an 80 bp-long dsDNA due to the expected stronger bending of the gap-bearing DNA constructs and, consequently, lower signals when the dye is closer to graphene. Again, we calculate the graphene distance based on the fitted fluorescence lifetimes for four different gap sizes (Figure 4B). The data show decreasing graphene distances with mean values and standard error of (25.3 ± 0.2) nm, (23.7 ± 0.5) nm and (21.0 ± 0.5) nm for 0 to 2 nt gap size, respectively. Supposedly, the increasing disruption of the base stacking with a larger gap yields shorter average distances. However, for a 3 nt gap size, with the distance does not decrease further with a measured value of (21.7 ± 0.3) nm.

**Figure 4.**
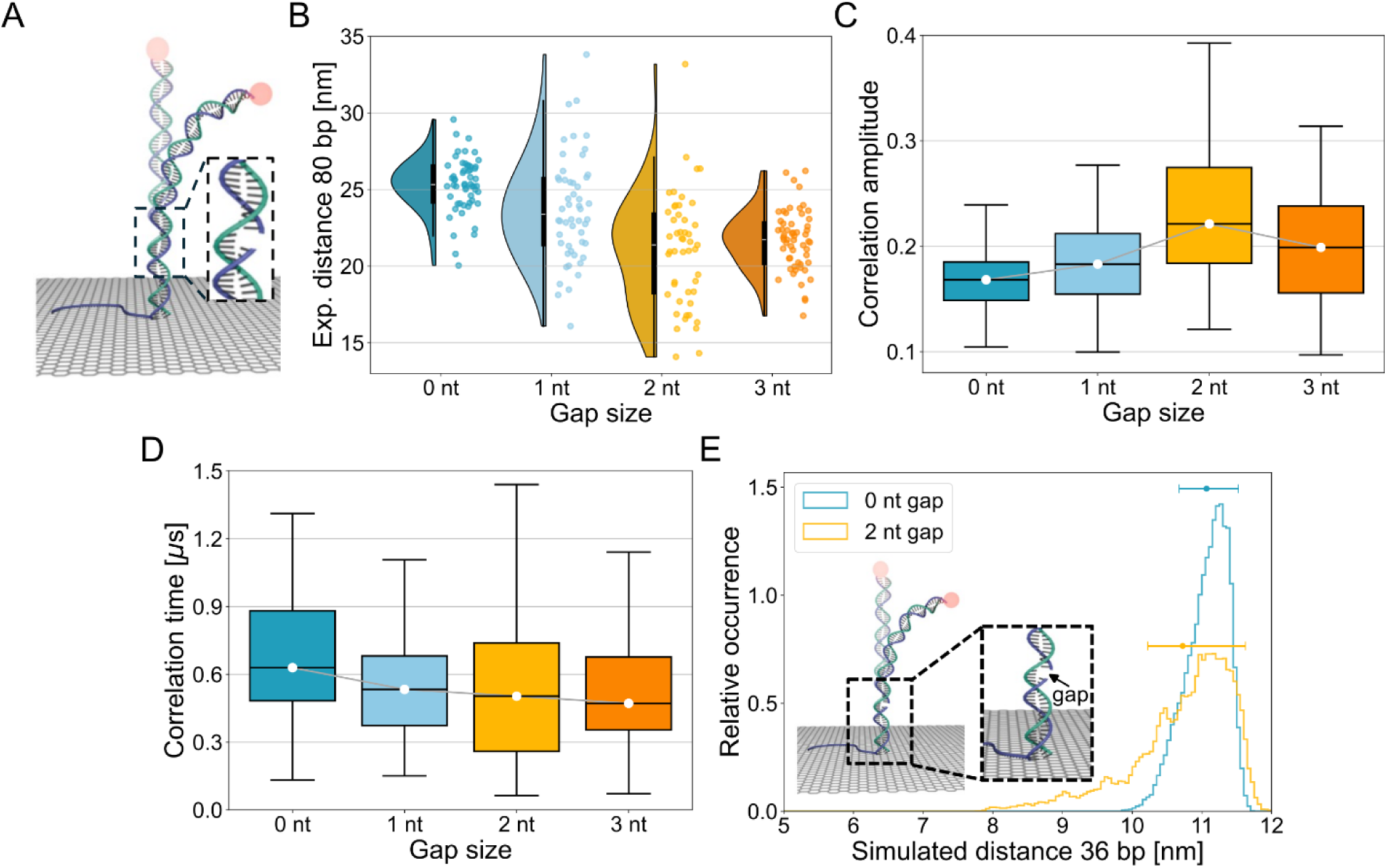
Altering fluctuations by introducing varying gaps to the DNA. **(A)** Schematic of GETvNA sample without (smaller, intact) and with a gap in the dsDNA (zoom-in). **(B)** Experimentally determined distances deviated from the measured fluorescence lifetime for 80 bp dsDNA constructs with different gap sizes (*n* = 56, 52, 49, 57). Within the violinplot, the white dot indicates the median, the box marks the first to third quartiles and the whiskers range from minimum to maximum, excluding outliers. **(C)** Box plot of experimentally determined correlation amplitudes. **(D)** Correlation times corresponding to the amplitude values and gap sizes in (B). The box plots in (C) and (D) display each distribution excluding outliers with the whiskers ranging from minimum to maximum. The respective boxes visualize the first and third quartile separated by the median. The median values are connected by a grey line to guide the eye. See Supplementary Figures S31-S33 for all data points including outliers. **(E)** Histogram from MD simulations depicting the probability of residence for an intact, 0 nt gap and a 2 nt gapped 36 bp-long DNA strand as a function of the distance *z* and as shown by the sketch. The dot and error bar represent the mean distance and the respective *1/e*-fluctuation range for each probability distribution.

The correlation analysis shows sub-microsecond fluctuations and an increase in correlation amplitude from 0.17 for 0 nt gap size up to 0.25 for 2 nt gap size, suggesting an increase in fluctuation range *Δz* (Figure 4C). Notably, the respective standard deviation of the three samples increases in a similar fashion from 0.04 to 0.13. For the 3 nt gap, the determined correlation amplitude (mean and standard error) of 0.21 ± 0.01 is slightly smaller than in the 2 nt case, potentially, a sign of a saturation in the gap-induced displacement.^73^ The corresponding correlation times exhibit a slight decrease with increasing gap size with (0.69 ± 0.04) µs for 0 nt gap size towards (0.58 ± 0.07) µs for 2 nt gap size (Figure 4D). For the 3 nt gap, the data show (0.58 ± 0.08) µs correlation time for mean and standard error.

The results are corroborated by MD simulations. Again, the MD simulations are carried out for a 36 bp-long dsDNA with a 2 nt gap (missing bp at position 17 & 18 from graphene) that is compared to the respective complete DNA strand (Supplementary Table S3). The results reveal a slightly decreased mean distance and strongly increased fluctuation range for dsDNA with a 2 nt gap, highlighting the prominent role of base stacking for the structural stability of the DNA double helix (Figure 4E and Supplementary Figure S34). Besides the increased bending, the helix’ rise, minor and major groove are preserved upon the introduction of the 2-nt gap (Supplementary Table S4), in agreement with earlier reports that investigated the impact of phosphate backbone cleavage in nicked DNA.^74^ The distance distribution of the DNA with a 2 nt gap is generally broader also allowing states with larger graphene distance (Fig. 4E) that are related to the increased contour length of single-stranded regions both for the two unpaired nucleotides as well as for potentially fraying nucleotides next to the gap. As the gap acts as the terminus of two continuous duplexes with the adjacent bases being able to flip out of the helix on the covered microsecond timescale, the overall base stacking is reduced yielding an increased contour length which includes a state of larger distance to graphene.^75^ A:T base pairs fray more easily than G:C pairs. Because the sequence used in our analysis has two A:T pairs right next to the gap, this increased fraying noticeably affects the simulated distance.^76^ Nonetheless, it should be noted that the applied parametrization of the DNA is tailored for intact double strands and, for this, might overestimate the propensity of fraying.

Finally, we note a weak anticorrelation of correlation amplitude and distance to graphene increasing from 0 to 2 nt gap size (Supplementary Figure S35). This might indicate that molecules that tend to be preferably in a stacked conformation show less dynamics. This observation agrees with the MD simulations showing a larger diffusion exponent for the DNA including a gap (Supplementary Figure S36). Nonetheless, we note that this increase in anticorrelation was not observed in the experiment for the sample with a 3 nt gap.

Lastly, the analysis of sub-microsecond fluorescence fluctuations was employed for comparing different types of DNA mismatches (Figure 5A). To this end, a 66 bp-dsDNA with mismatches at the 10^th^ (G:T), 11^th^ (C:T), 10^th^ & 11^th^ (double), or 10^th^-12^th^ (triple) nucleotide counting from graphene was used. The determined distances show variations up to 1.5 nm for the samples bearing a single G:T or C:T mismatch (Figure 5B). Interestingly, for the G:T mismatch, the resulting distance is 0.7 nm smaller than for the respective sample without mismatch, showing mean values and standard errors of (22.2 ± 0.2) nm versus (21.5 ± 0.2) nm. In contrast, the sample with C:T mismatch shows a 0.3 nm larger graphene distance, (22.5 ± 0.3) nm. Based on the G:T mismatch, a T:T mismatch was added in the adjacent bp pointing away from graphene and a distance of (22.0 ± 0.3) nm was determined. Further, the triple mismatch shows a graphene distance of (20.9 ± 0.3) nm, i.e. a 1.3 nm decreased graphene distance compared to the sample without mismatches. Here and in the case of the C:T mismatch, it should be noted that the analysis reveals the presence of two populations indicating at least two mismatch conformations which are stable at least over many seconds.

**Figure 5.**
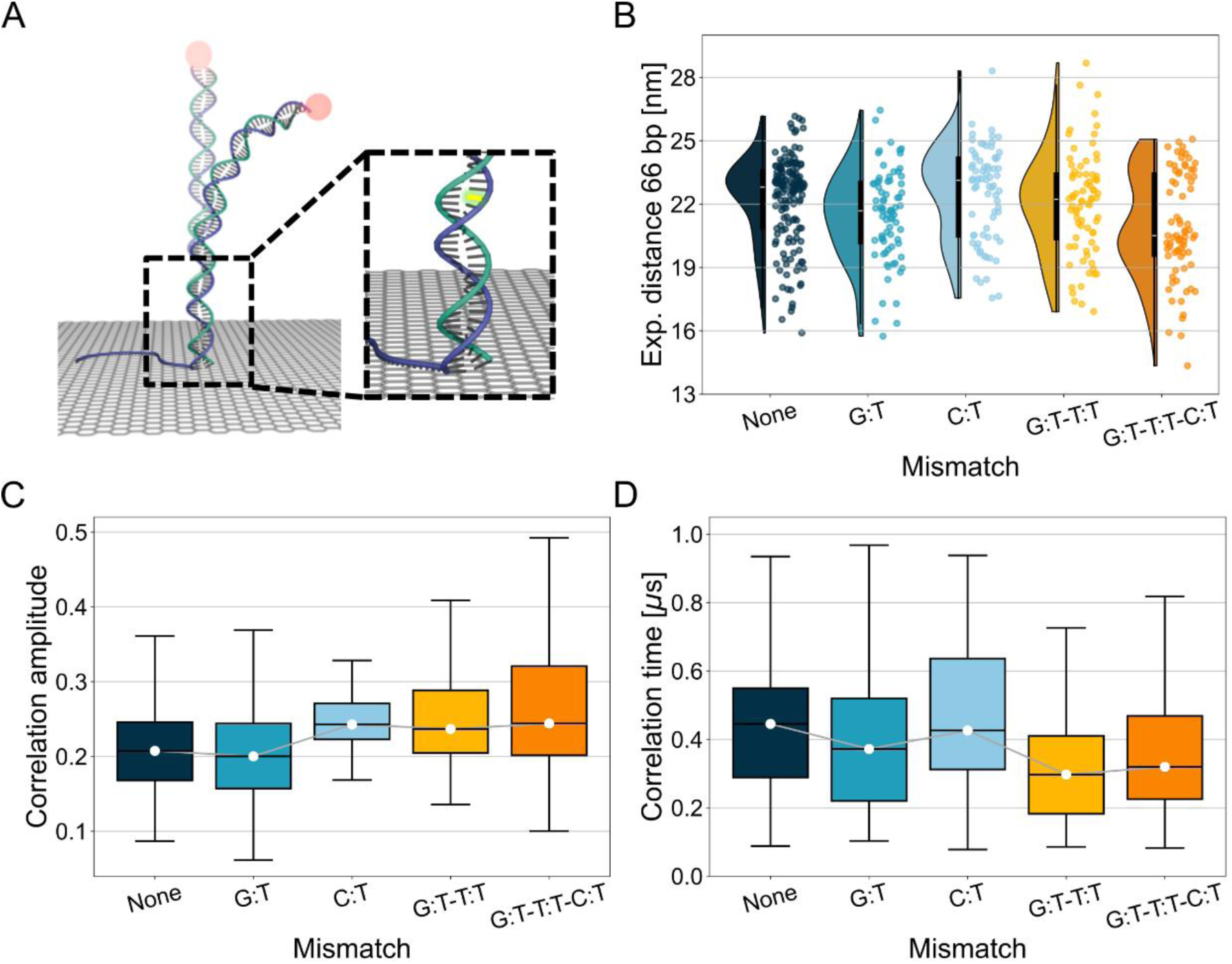
Application of measuring rapid fluctuations for distinguishing DNA mismatches. **(A)** Sketch of fluctuating DNA with a DNA mismatch highlighted in yellow. **(B)** Distances to graphene retrieved from fitting the measured fluorescence lifetime for a 66 bp-long DNA with the indicated mismatch type (*n* = 155, 76, 73, 77, 75). Within the violinplot, the white dot indicates the median, the box marks the first to third quartile and the whiskers range from minimum to maximum, excluding outliers. **(C)** Box plot of experimentally determined correlation amplitudes. **(D)** Correlation times corresponding to the measured amplitude values in (B). The box plots in (C) and (D) display each distribution excluding outliers with the whiskers ranging from minimum to maximum. The respective boxes visualize the first and third quartile separated by the median. The median values are connected by a grey line to guide the eye. See all data points including outliers in Supplementary Figures S37-S39.

When investigating fluctuations, mismatch-free and G:T-mismatched DNA show almost identical correlation amplitudes of 0.22 ± 0.01 and 0.21 ± 0.01, respectively. The C:T mismatch sample, however, showed a larger correlation amplitude of 0.25 ± 0.01, similar to the double and triple mismatch exhibiting both 0.26 ± 0.01 (Figure 5C). The discrepancy in the correlation amplitude for the single mismatches is consistent with the G:T-mismatch showing similar energetic characteristics as the Watson-Crick paired C:G nucleobases, yet, the energetic barrier is much lower for creating fluctuations in a C:T mismatch.^77^ Samples with an increasing number of mismatches also exhibit broader amplitude distributions indicating enhanced dynamic heterogeneity. The respective mean correlation time amounts to (0.44 ± 0.02) µs for the sample without mismatch (Figure 5D). It does not change substantially for the sample with G:T mismatch, exhibiting a correlation time of (0.41 ± 0.03) µs. Interestingly, for the C:T mismatch it is slightly larger with (0.49 ± 0.03) µs. A substantial difference, however, is only visible for the double or triple mismatched samples with correlation times of (0.31 ± 0.02) µs and (0.36 ± 0.02) µs. This reduction compared to the sample without or bearing a single mismatch is in line with the assumption that a mismatch will interrupt the intact double strand allowing faster intramolecular dynamics. Yet, it is surprising that the single mismatches do not cause a measurable decrease in the mean correlation time, potentially, because the nearest-neighbor interactions of a single mismatched nucleobase with its adjacent nucleobases are sufficient to maintain the stability of the DNA.^78^ Interestingly, we note an increasing anticorrelation between correlation amplitude and graphene distance for a growing number of mismatches, suggesting that mismatches similar to gaps disrupt the DNA base stacking (Supplementary Figures S40).

Overall, the results suggest that more stable base stacking and pairing ensures persistent helix properties spanning beyond the defect site. In contrast, a defect will interrupt the persistence and, *e*.*g*., lead to more intramolecular dynamics showing how chemical modifications within the DNA cause physical changes.

## Conclusion

In summary, we demonstrate that GETvNA combined with shrinking-gate fluorescence correlation spectroscopy can resolve sub-microsecond conformational dynamics in single DNA molecules on graphene. The method reports axial bending fluctuations through correlated intensity and lifetime changes and thereby separates structural dynamics from photophysical processes. We show that the fluctuation behavior of dsDNA depends on ionic strength and is sensitively altered by local structural defects such as gaps or mismatches. Especially, the application of the fluctuation analysis to DNA mismatches reveals the full potential of the method to elucidate subtle structural variations at the single-molecule level. For example, such differences in the fluctuation behavior of e.g. single C:T and G:T mismatches are revealed and could be related to the stability of the overall structure^14,77^. Here, it is noteworthy that the applied analysis is not limited to guanine- or thymine-bearing mismatches, as it is the case for the imino proton exchange used in state-of-the-art nuclear-magnetic resonance experiments^79,80^. Generally, the experiments involving gaps or mismatches reveal that with an increasing disruption of the base stacking the correlation amplitude increases but the correlation time decreases.

Molecular dynamics simulations and kinetic Monte Carlo modeling rationalize the physical interpretation of the measured correlation amplitudes and timescales. The comparison of end-to-end and end-to-graphene distributions also reveals the higher sensitivity of GETvNA compared to FRET for studying these biomolecular dynamics. Our results agree with previous experimental work, yet extend those towards shorter dsDNA.^2^ So far, our analysis is taking into account DNA bending but ignores other forms of motion like twisting.^81–84^ The resolution of GETvNA in combination with sgFCS is sufficient for distinguishing populations but does not have the resolution for the systems studied to reliably assign single molecules to the respective distributions.

Beyond the present model systems, the presented framework enables mechanistic studies of fast nucleic-acid motions and their modulation by sequence, chemical damage and biomolecular interactions.

## Acknowledgements

The authors thank the members of the Tinnefeld lab for fruitful discussions and feedback. Furthermore, we thank Gereon Brüggenthies for help with the preparation of the DNA origami structure.

## Author Contributions

The conceptualization of the project was carried out by P.T. L.R., J.H. and L.C. curated the data. Formal analysis was conducted by L.R., J.H. and L.C. The funding acquisition was led by P.T. and B.P.F. Investigations were carried out by L.R., J.H. and L.C. The methodology was developed by L.R., J.H., L.C. and T.S. The project was administered by L.R. The resources were provided by P.T. and B.P.F. The analysis software was developed by L.R. Supervision was provided by P.T. and B.P.F. Validation was performed by T.S., A.M.S., P.T. and B.P.F. L.R. and L.C. created the visualizations. The writing of the original draft was carried out by all authors as well as the review and editing.

## Data Availability

All experimental data supporting the findings of this work are available upon request.

## Code Availability

The Python-based code for the graphical user interface that was employed for analyzing the experimental single-molecule fluorescence data presented within this work is available from a public repository: https://gitlab.lrz.de/tinnefeldlab/fcs-gui.

