## Supplementary Information for "Resolving Sub-Microsecond Conformational Dynamics of Vertical Nucleic Acids on Graphene" (from another version of this preprint: version 1, 2026-06-10)

### 1 Supplementary Discussion

#### 1.1 Analysis of Data from Molecular Dynamics (MD) Simulation

*Graphene Analysis*

To assess the impact of explicitly treated graphene on the structure of the DNA, a 200 ns-simulation of the DNA on three graphene sheets, similar to (*1*), was performed. The average z-distance, end-to-end distance and helical structure parameters (Table S7) show deviations smaller than the standard deviations between the systems with and without explicitly treated graphene. It was therefore concluded that the given explicit graphene does not significantly affect the structure and dynamics of the DNA in the MD simulations and can be excluded from the calculations. The average distance between the center of mass (COM) of the first base pair and the graphene surface was calculated as 0.4 nm, and used to correct the z-distance derived from the virtual plane (Supplementary Figure S16).

*Plane Projection*

The DNA assumed a random spatial orientation in the simulation box before positional restraints were applied to the first two base pairs. Hence, a virtual plane for the calculation of the z-distance distribution of the last base pair resembling the distance measured to the graphene surface in the experiment had to be defined. A plane was defined between the COM of the first base pair and two periodic images in a way that leads to upright standing DNA molecules in the fully extended configurations. The *z*-distance was calculated as the vertical projection of the COM of the last base pair to this virtual plane. To benchmark this approach, the result was compared in the DNA-graphene system with the explicitly calculated COM-graphene distance. The benchmark showed perfect agreement between the virtual plane and the explicit graphene surface as reference with an average error of 0.2 Å (Supplementary Figure S17).

#### 2 Supplementary Figures

**Fig. S1.**

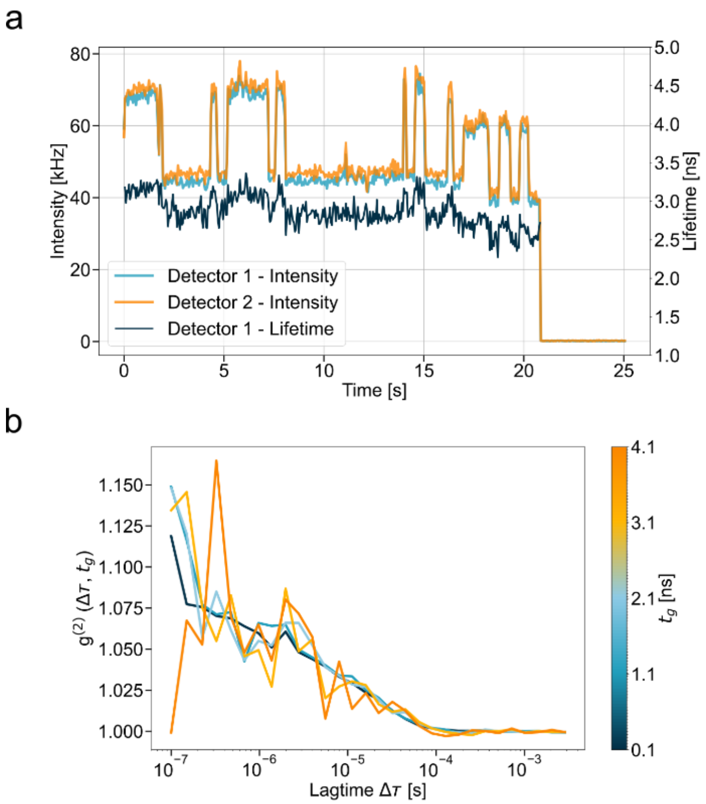

**Fig. S1. Figure 2 – Example from DNA origami measurement.** (a) Time-intensity trajectory detected at detectors 1 and 2 (in orange & blue) with time bins set to 50 ms. The black graph shows the corresponding fluorescence lifetime determined for detector 1. (b) Cross-correlation function for the detector signals 1 & 2 in the time window 0.0-20.7 s (showing a clear single-molecule signature for $g^{(2)}(0)$, see ref. (*2*)) analyzed by sgFCS using the color-coded time gates $t_g$.

**Fig. S2.**

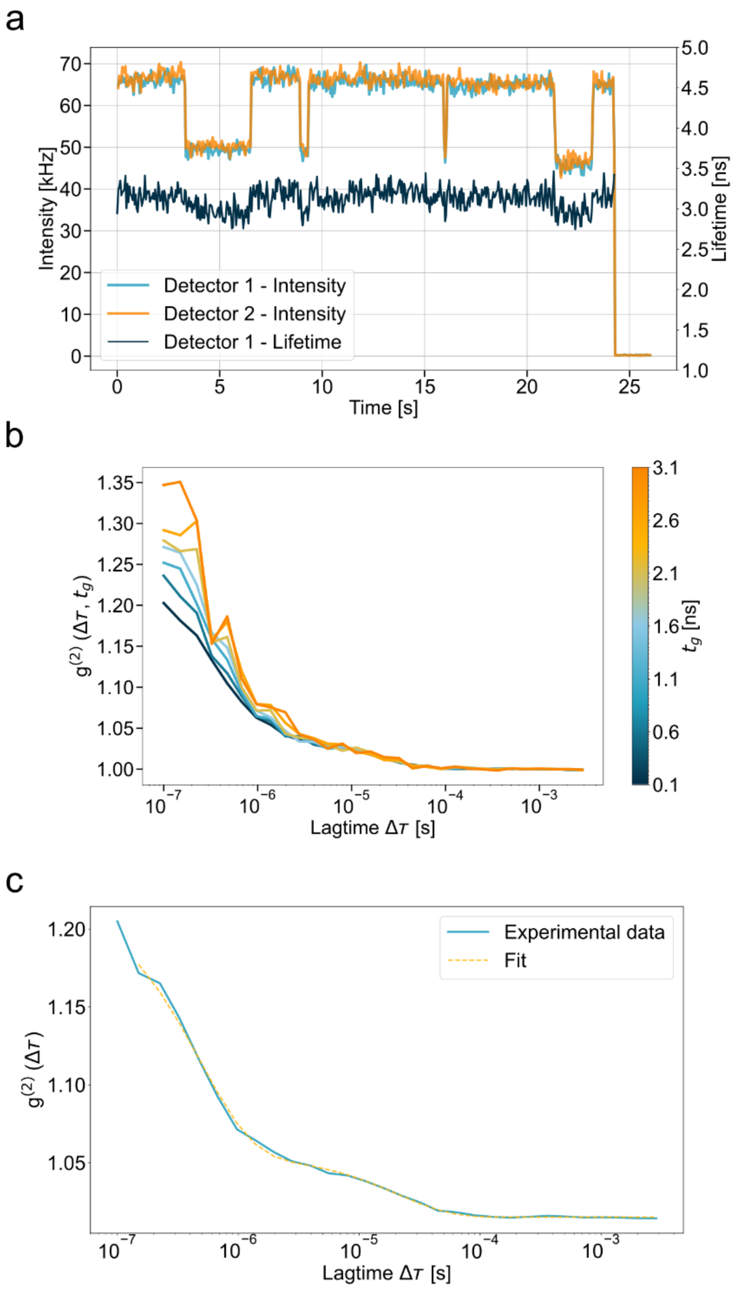

**Fig. S2. Figure 2 – Example 1.** (a) Time-intensity trajectory detected at detectors 1 and 2 (in orange & blue) with time bins set to 50 ms. The black graph shows the corresponding fluorescence lifetime determined for detector 1. (b) Cross-correlation function for the detector signals 1 & 2 in the time window 0.0-24.2 s analyzed by sgFCS using the color-coded time gates $t_g$, revealing dynamics on the sub-microsecond time scale. (c) Corresponding fluorescence cross-correlation using all detected photons and biexponential fit.

**Fig. S3.**

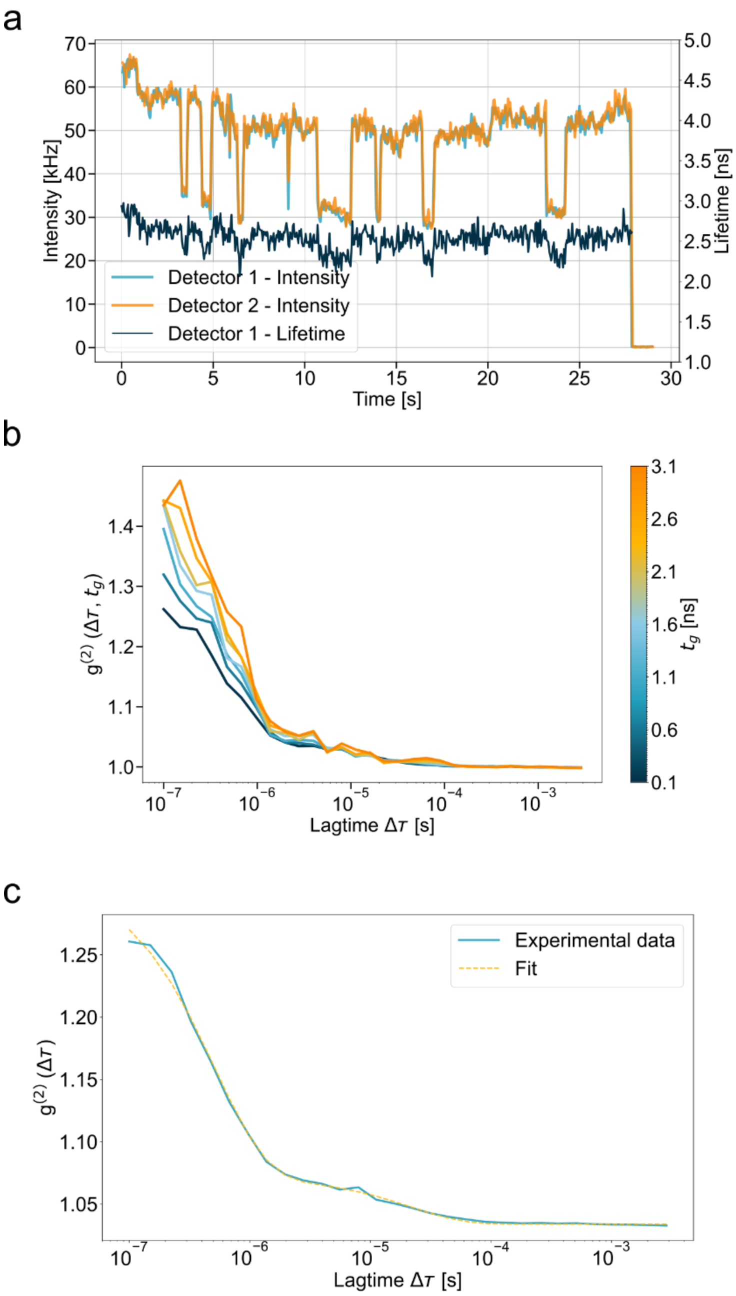

**Fig. S3. Figure 2 – Example 2.** (a) Time-intensity trajectory detected at detectors 1 and 2 (in orange & blue) with time bins set to 50 ms. The black graph shows the corresponding fluorescence lifetime determined for detector 1. (b) Cross-correlation function for the detector signals 1 & 2 in the time window 0.9-27.9 s analyzed by sgFCS using the color-coded time gates $t_g$, revealing dynamics on the sub-microsecond time scale. (c) Corresponding fluorescence cross-correlation using all detected photons and biexponential fit.

**Fig. S4.**

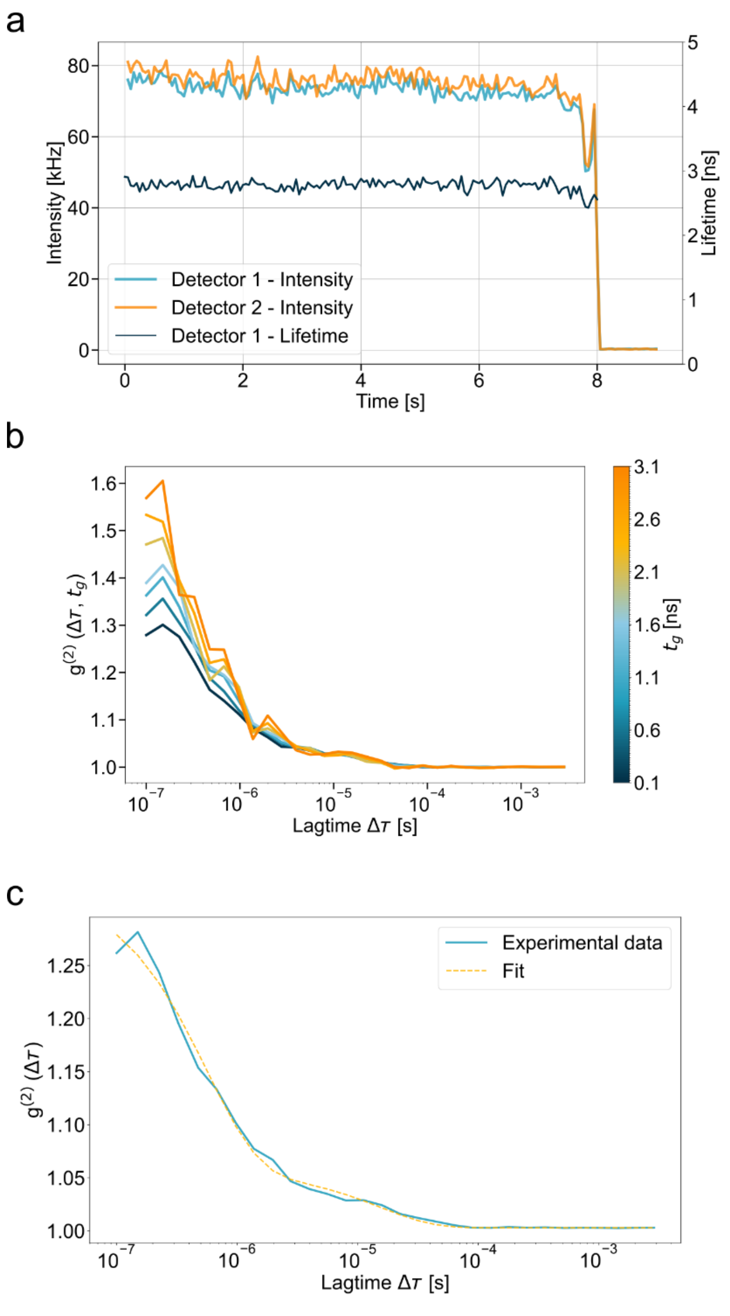

**Fig. S4. Figure 2 – Example 3.** (a) Time-intensity trajectory detected at detectors 1 and 2 (in orange & blue) with time bins set to 50 ms. The black graph shows the corresponding fluorescence lifetime determined for detector 1. (b) Cross-correlation function for the detector signals 1 & 2 in the time window 0.0-8.0 s analyzed by sgFCS using the color-coded time gates $t_g$, revealing dynamics on the sub-microsecond time scale. (c) Corresponding fluorescence cross-correlation using all detected photons and biexponential fit.

**Fig. S5.**

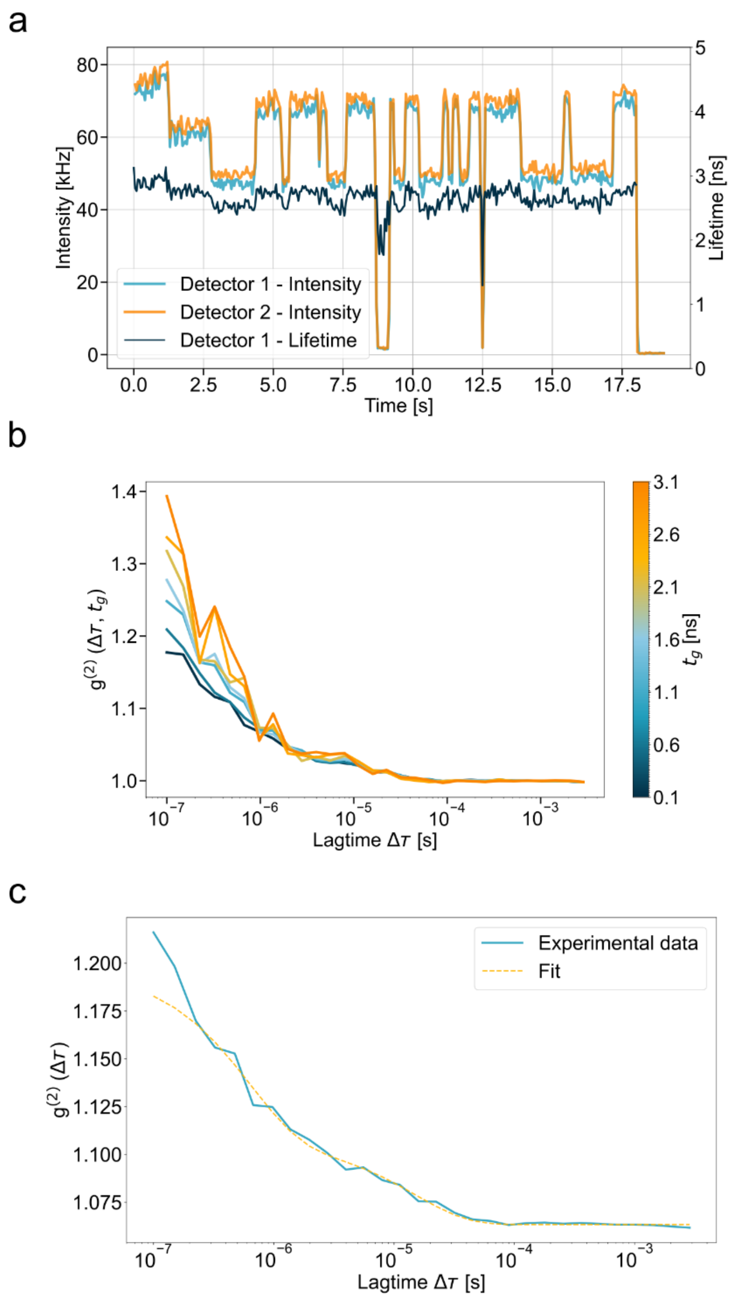

**Fig. S5. Figure 2 – Example 4.** (a) Time-intensity trajectory detected at detectors 1 and 2 (in orange & blue) with time bins set to 50 ms. The black graph shows the corresponding fluorescence lifetime determined for detector 1. (b) Cross-correlation function for the detector signals 1 & 2 in the time window 1.2-18.0 s analyzed by sgFCS using the color-coded time gates $t_g$, revealing dynamics on the sub-microsecond time scale. (c) Corresponding fluorescence cross-correlation using all detected photons and biexponential fit.

**Fig. S6.**

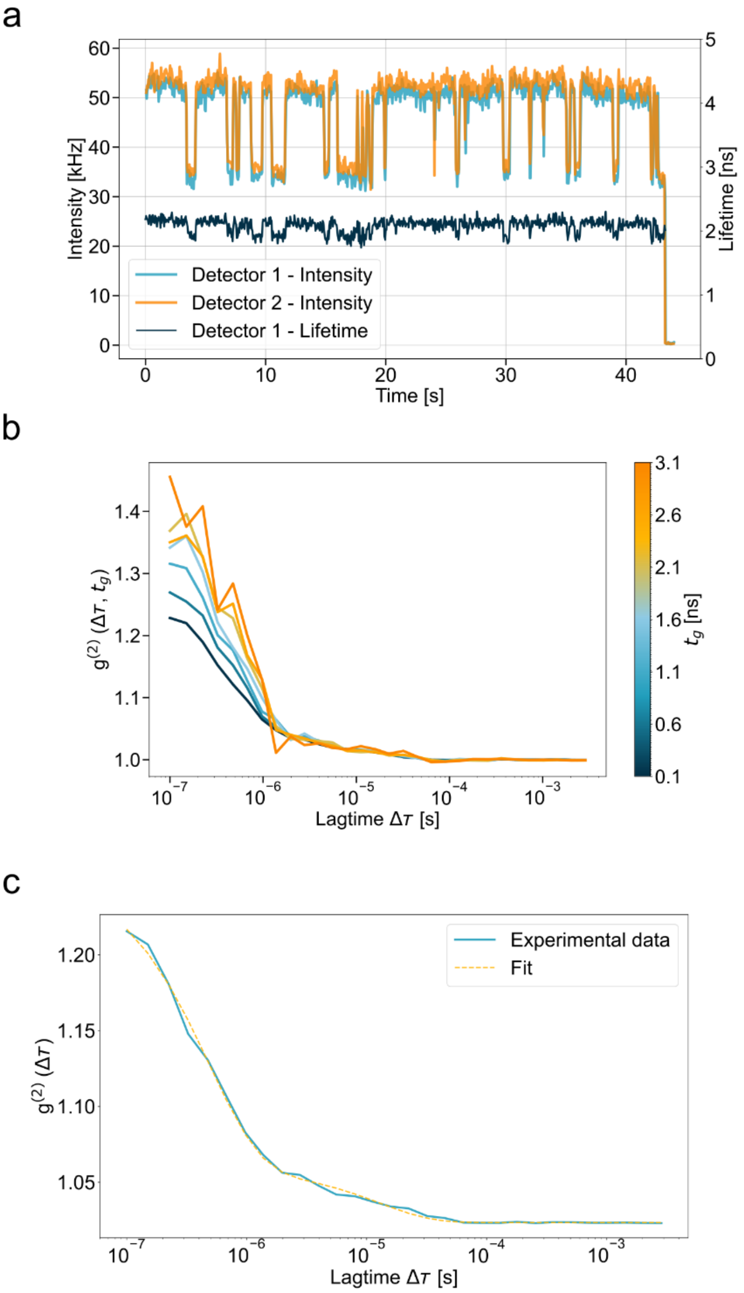

**Fig. S6. Figure 2 – Example 5.** (a) Time-intensity trajectory detected at detectors 1 and 2 (in orange & blue) with time bins set to 50 ms. The black graph shows the corresponding fluorescence lifetime determined for detector 1. (b) Cross-correlation function for the detector signals 1 & 2 in the time window 0.0-43.2 s analyzed by sgFCS using the color-coded time gates $t_g$, revealing dynamics on the sub-microsecond time scale. (c) Corresponding fluorescence cross-correlation using all detected photons and biexponential fit.

**Fig. S7.**

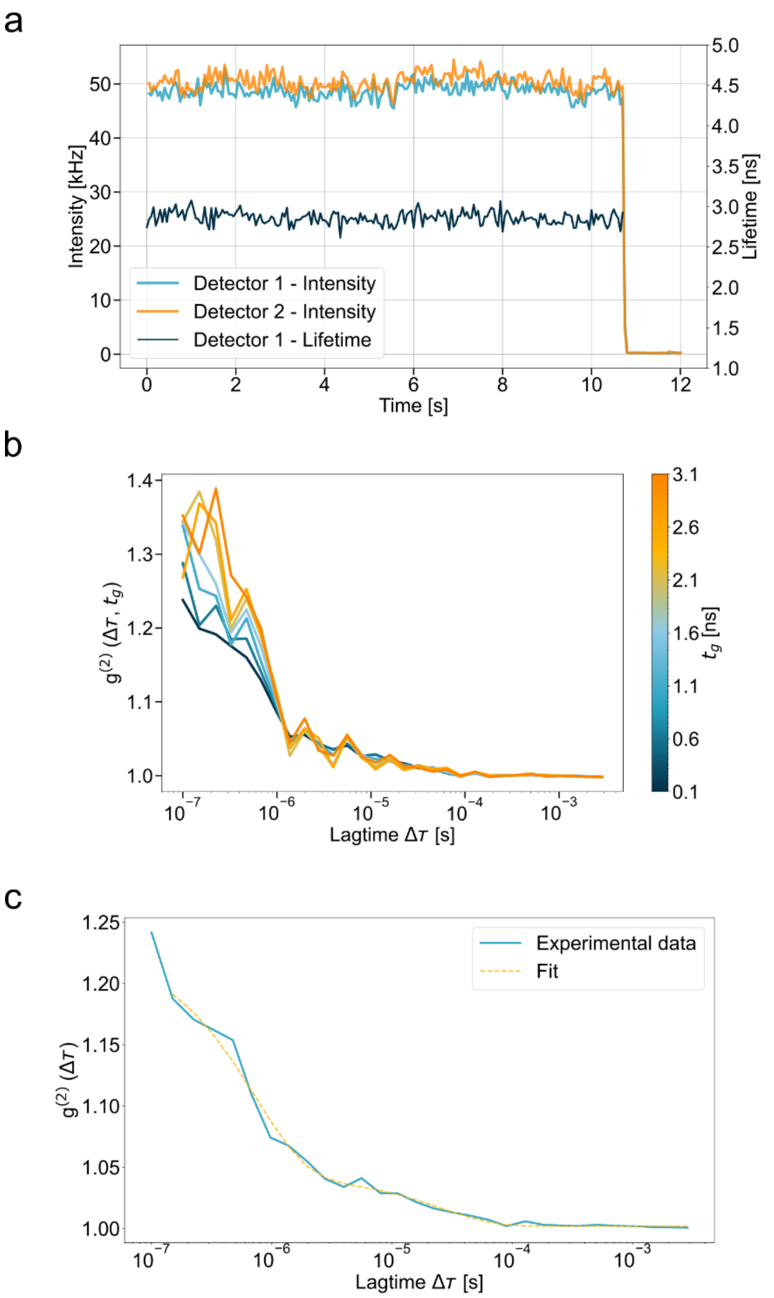

**Fig. S7. Figure 2 – Example 6.** (a) Time-intensity trajectory detected at detectors 1 and 2 (in orange & blue) with time bins set to 50 ms. The black graph shows the corresponding fluorescence lifetime determined for detector 1. (b) Cross-correlation function for the detector signals 1 & 2 in the time window 0.0-10.7 s analyzed by sgFCS using the color-coded time gates $t_g$, revealing dynamics on the sub-microsecond time scale. (c) Corresponding fluorescence cross-correlation using all detected photons and biexponential fit.

**Fig. S8.**

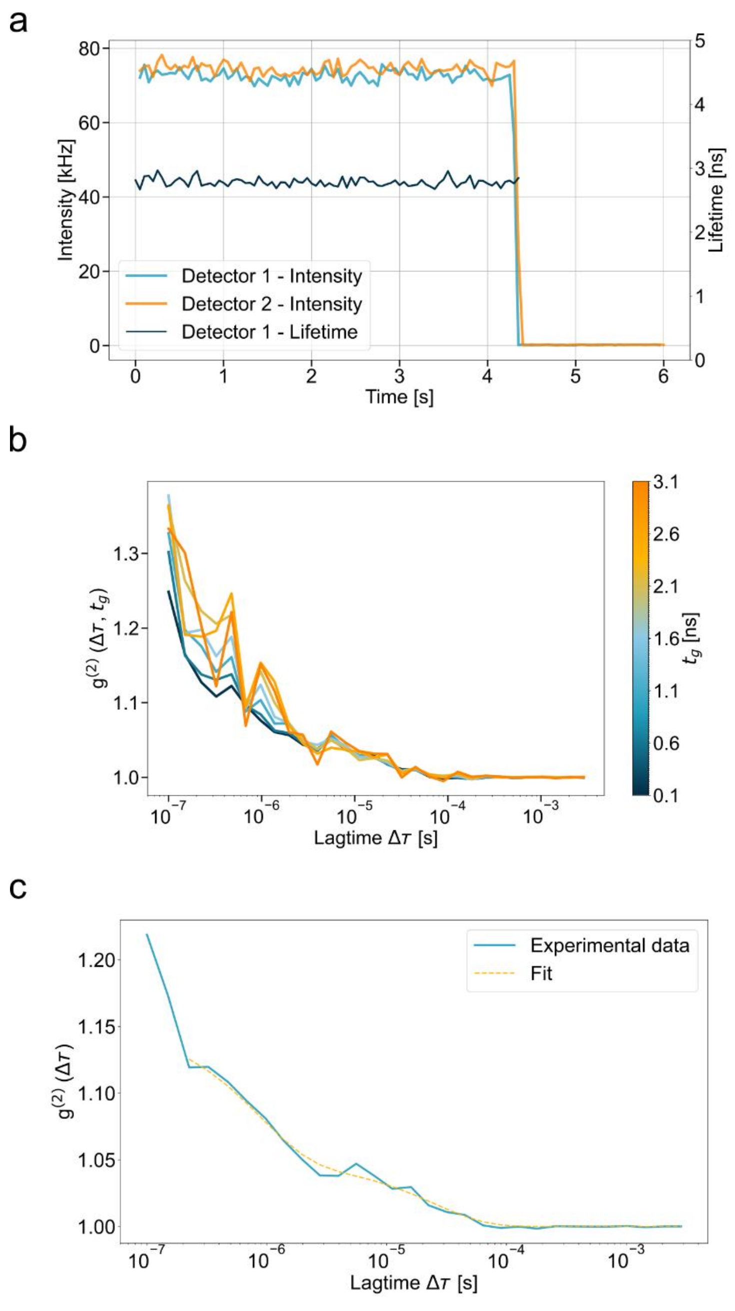

**Fig. S8. Figure 2 – Example 7.** (a) Time-intensity trajectory detected at detectors 1 and 2 (in orange & blue) with time bins set to 50 ms. The black graph shows the corresponding fluorescence lifetime determined for detector 1. (b) Cross-correlation function for the detector signals 1 & 2 in the time window 0.0-4.3 s analyzed by sgFCS using the color-coded time gates $t_g$, revealing dynamics on the sub-microsecond time scale. (c) Corresponding fluorescence cross-correlation using all detected photons and biexponential fit.

**Fig. S9.**

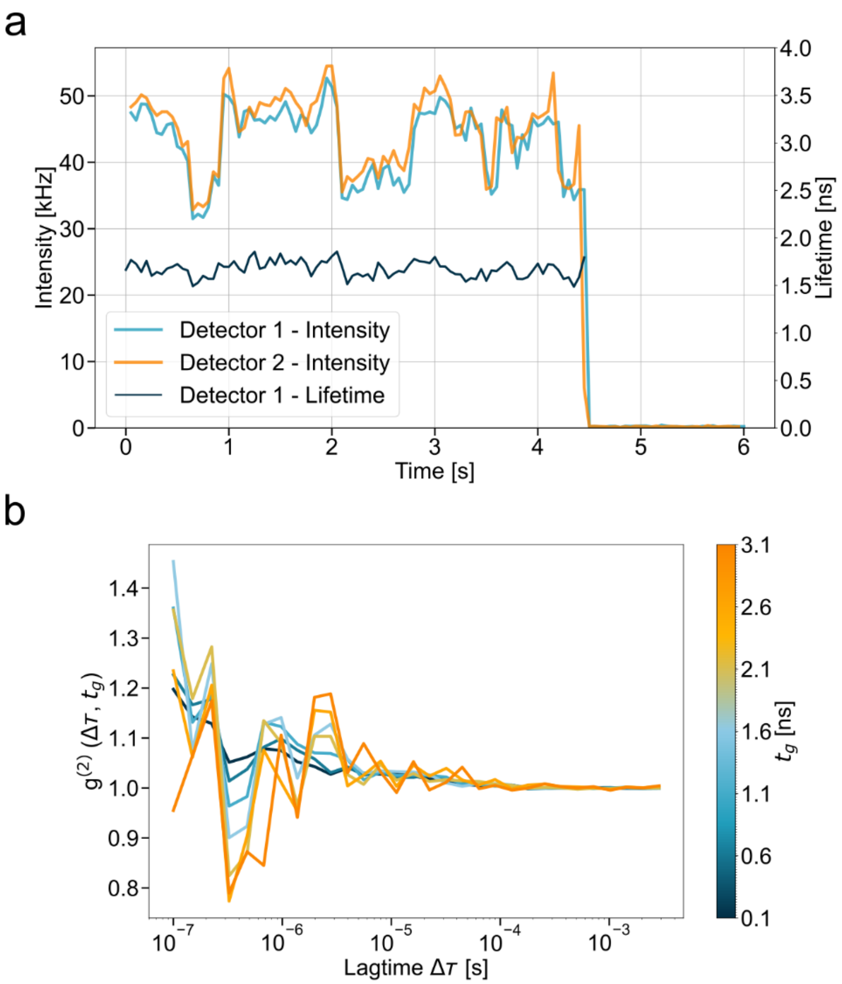

**Fig. S9. Figure 2 – Example 1 not showing dynamics.** (a) Time-intensity trajectory detected at detectors 1 and 2 (in orange & blue) with time bins set to 50 ms. The black graph shows the corresponding fluorescence lifetime determined for detector 1. (b) Cross-correlation function for the detector signals 1 & 2 in the time window 0.0-4.4 s analyzed by sgFCS using the color-coded time gates $t_g$, revealing dynamics on the sub-microsecond time scale.

**Fig. S10.**

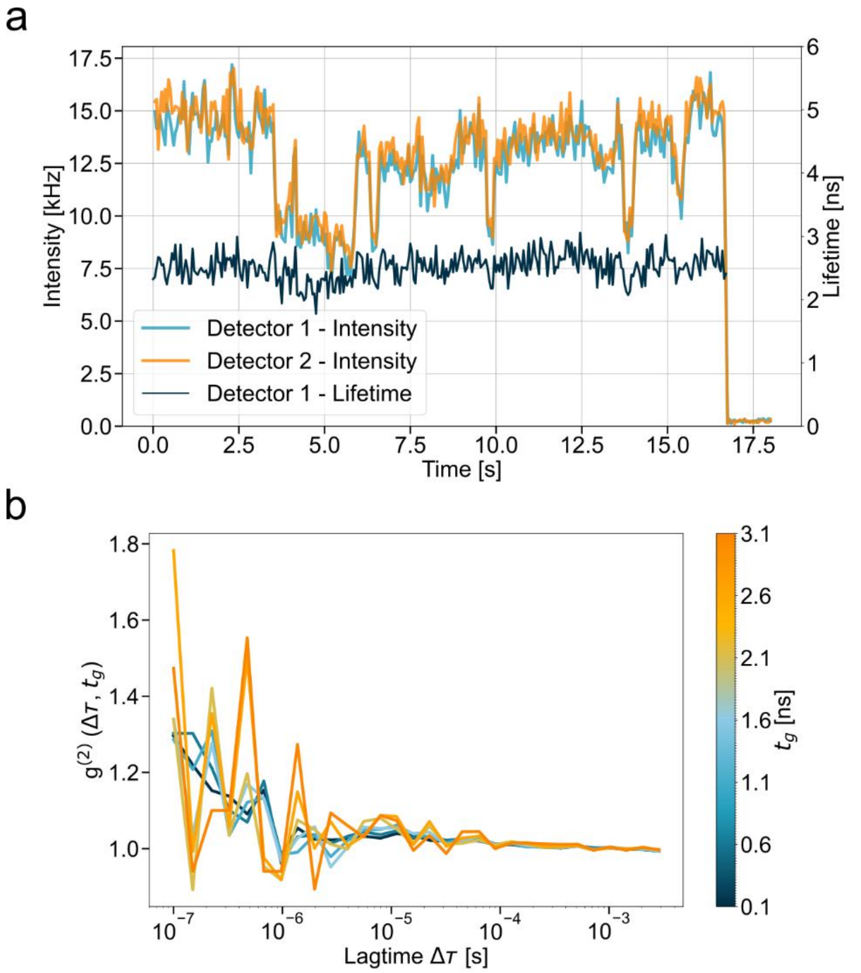

**Fig. S10. Figure 2 – Example 2 not showing dynamics.** (a) Time-intensity trajectory detected at detectors 1 and 2 (in orange & blue) with time bins set to 50 ms. The black graph shows the corresponding fluorescence lifetime determined for detector 1. (b) Cross-correlation function for the detector signals 1 & 2 in the time window 0.0-16.7 s analyzed by sgFCS using the color-coded time gates $t_g$, revealing dynamics on the sub-microsecond time scale.

**Fig. S11.**

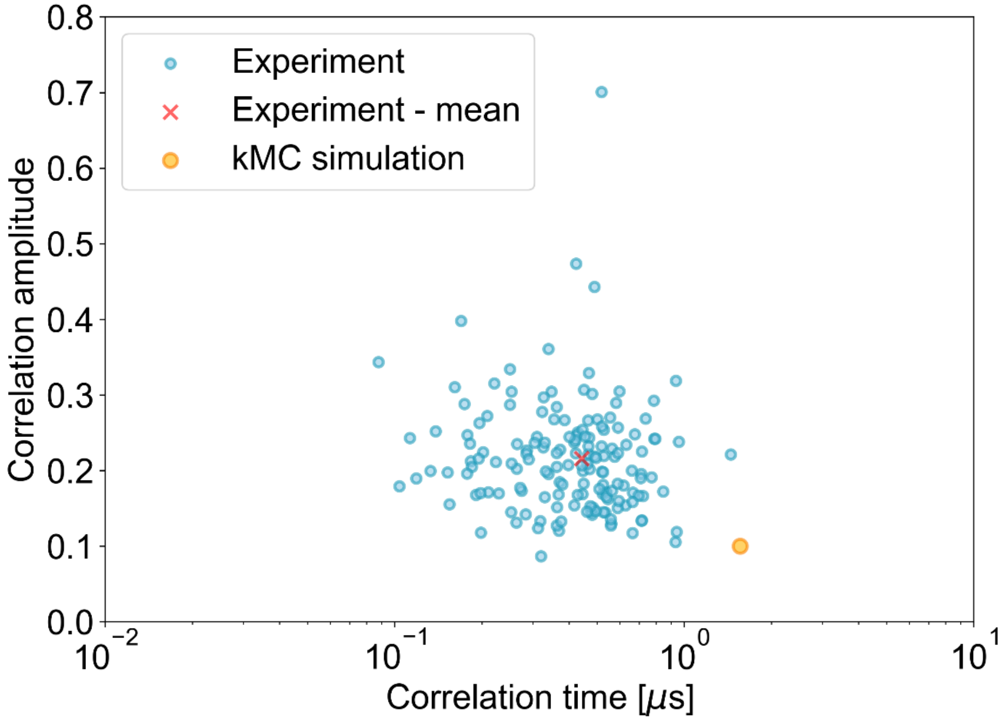

**Fig. S11. Figure 2 – Correlation time versus amplitude.** The point cloud shows the fitted correlation amplitudes and correlation times determined experimentally for the 66 bp dsDNA in 100 mM NaCl. The red cross denotes the mean of the data points. The enlarged orange dot shows the mean value of the kMC simulation runs, according to Supplementary Figure S26 for 50 nm persistence length.

**Fig. S12.**

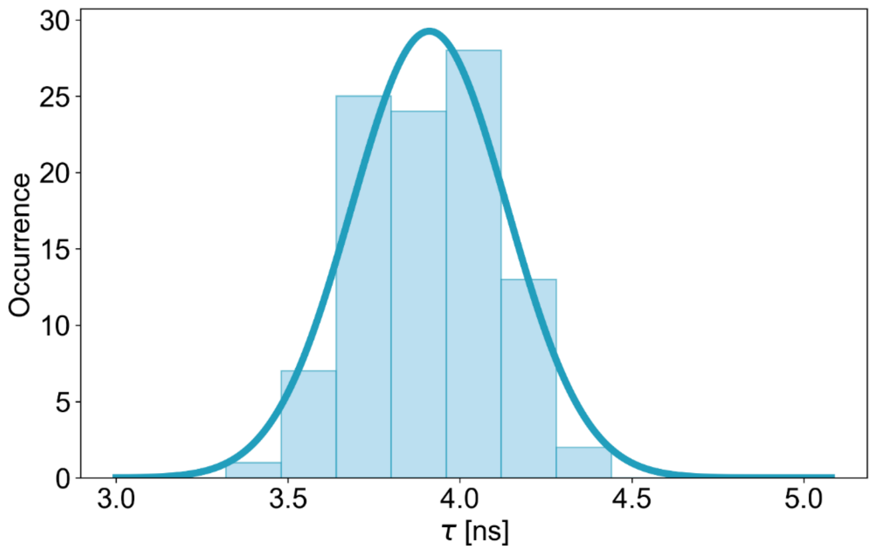

**Fig. S12. Figure 3 – Histogram of lifetimes (τ) from glass measurement.** The fit of 100 datapoints by a Gaussian function shows a mean value and standard error of (3.91 ± 0.02) ns. The measurement was performed at 100 mM NaCl concentration with the same buffer and imaging conditions as outlined in the Methods section. The fluorescent dye was attached to a 66 bp-long dsDNA that was anchored to glass by biotin and neutravidin.

**Fig. S13.**

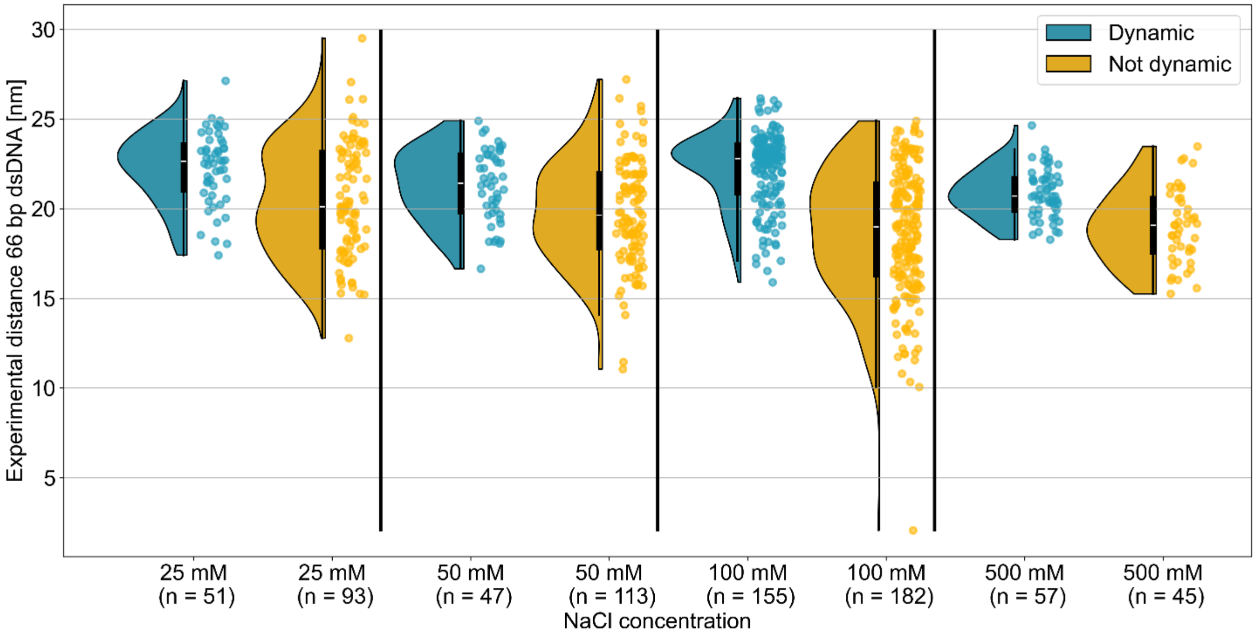

**Fig. S13. Figure 3 – Distances grouped by dynamics.** The plot displays all experimentally determined graphene distances for a 66 bp-long dsDNA at 25 mM to 500 mM sodium chloride concentration. The classification whether dynamic or not was done based on the respective shrinking-gate correlation. Each half violinplot on the left displays the distribution of the *n* datapoints as depicted by the dots on the right. The interior of the violinplot shows a box-and-whisker plot with the inter quartile range from the first to the third quartile shown as a black box separated by the median value in white. The whiskers range from minimum to maximum value. The results for each condition are separated by a vertical black line.

**Fig. S14.**

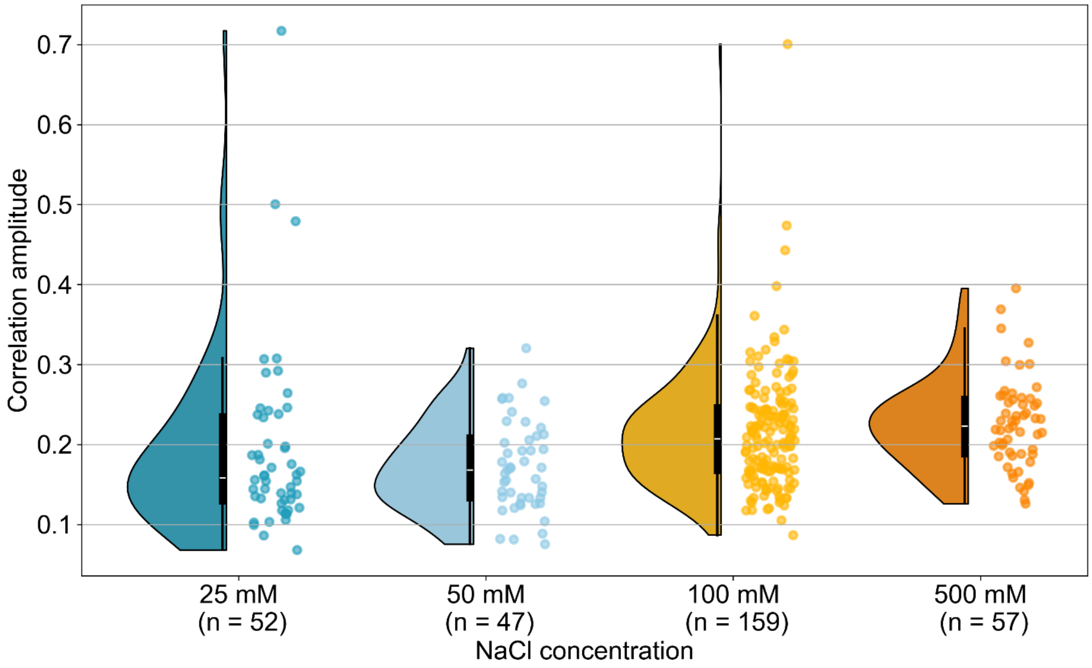

**Fig. S14. Figure 3 – Distribution of correlation amplitudes.** The plot displays all experimentally determined correlation amplitudes for a 66 bp-long dsDNA at 25 mM to 500 mM sodium chloride concentration. Each half violinplot on the left displays the distribution of the *n* datapoints as depicted by the dots on the right. The interior of the violinplot shows a box-and-whisker plot with the inter quartile range from the first to the third quartile shown as a black box separated by the median value in white. The whiskers range from minimum to maximum value.

**Fig. S15.**

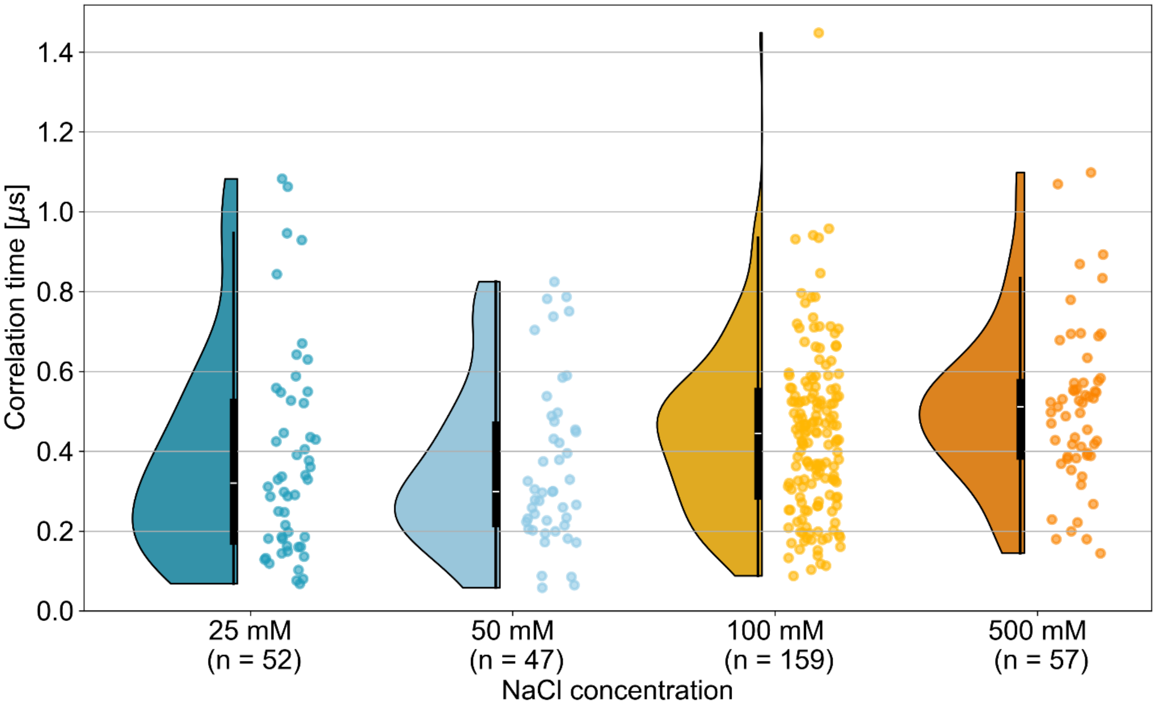

**Fig. S15. Figure 3 – Distribution of correlation times.** The plot displays all experimentally determined correlation times for a 66 bp-long dsDNA at 25 mM to 500 mM sodium chloride concentration. Each half violinplot on the left displays the distribution of the *n* datapoints as depicted by the dots on the right. The interior of the violinplot shows a box-and-whisker plot with the inter quartile range from the first to the third quartile shown as a black box separated by the median value in white. The whiskers range from minimum to maximum value.

**Fig. S16.**

a

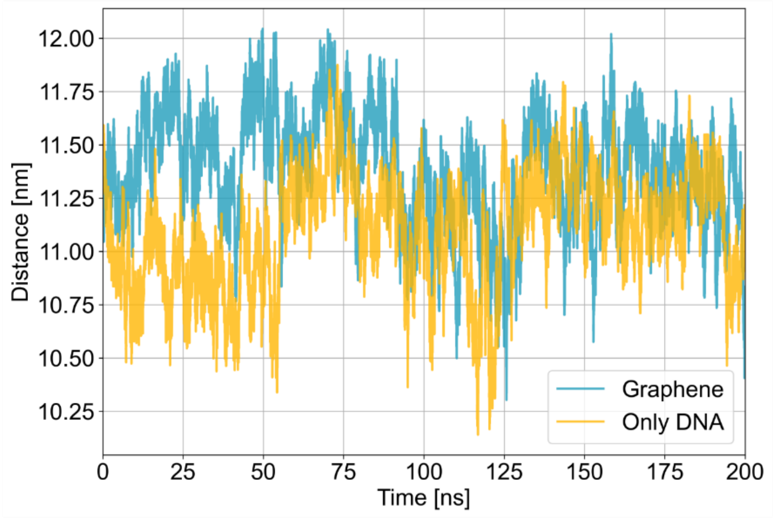

b

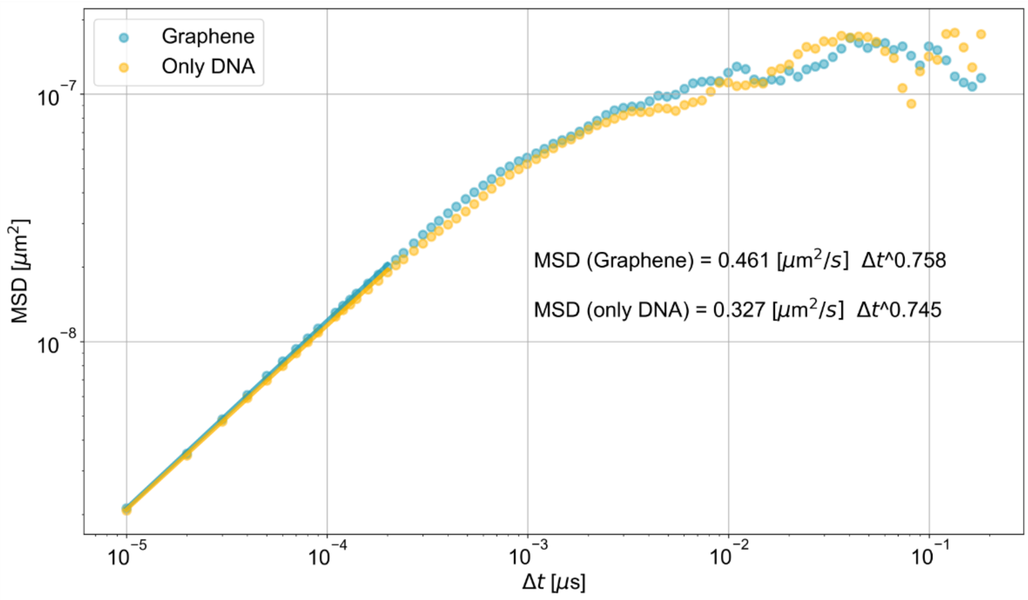

**Fig. S16. Figure 3 – MD simulation: with versus without graphene.** The MD simulation was performed twice for the same sequence according to Supplementary Table S3, with and without graphene at 50 mM sodium concentration. (a) The resulting data for the projected z-distance of the 36 bp dsDNA are compared. (b) Histogram of the simulated distances. (c) Plot of the mean squared displacement (MSD) versus time lag $\Delta t$ with the respective linear fit. The comparison reveals no significant influence of the presence on graphene on the conformation or the dynamics of the dsDNA.

**Fig. S17.**

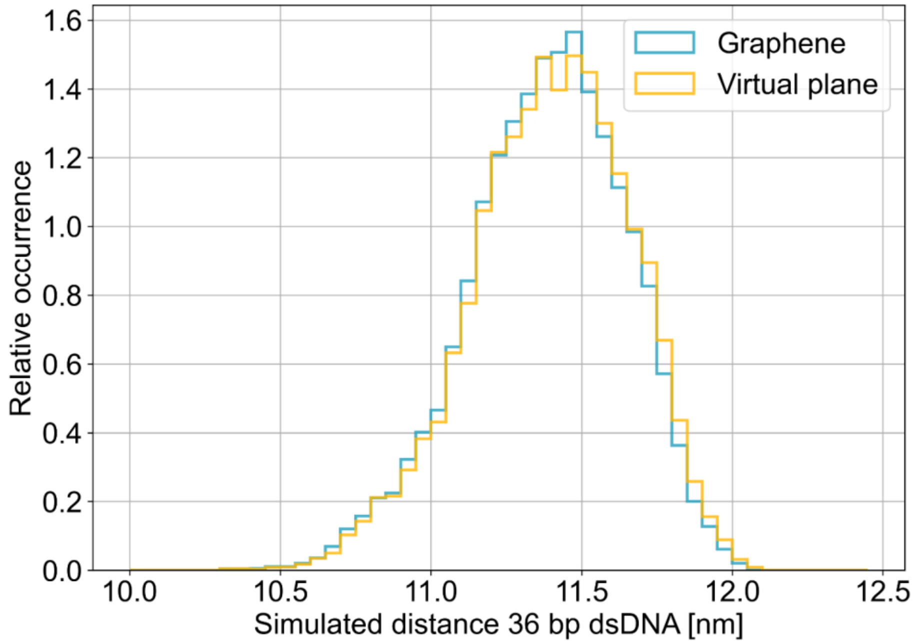

**Fig. S17. Figure 3 – MD simulation: virtual plane parallel to graphene.** The MD simulation with and without graphene (Supplementary Figure S16) revealed that the three layers of carbon atoms have a negligible influence on the conformation and dynamics of the dsDNA. Hence, and in the interest of computational resources, the graphene is not simulated, yet a virtual plane that, in principle, always extends parallel to graphene needs to be found. See the discussion in Supplementary Discussion's section 1.1 for details. Here, the simulated distance from the same simulation run measured from graphene is compared to the distance from the virtual plane revealing an offset value of 0.4 nm. Here, for better visual clarity, the distances for 'Virtual plane' are already corrected by this offset value. Subsequently, this value is added in all simulations without graphene.

**Fig. S18.**

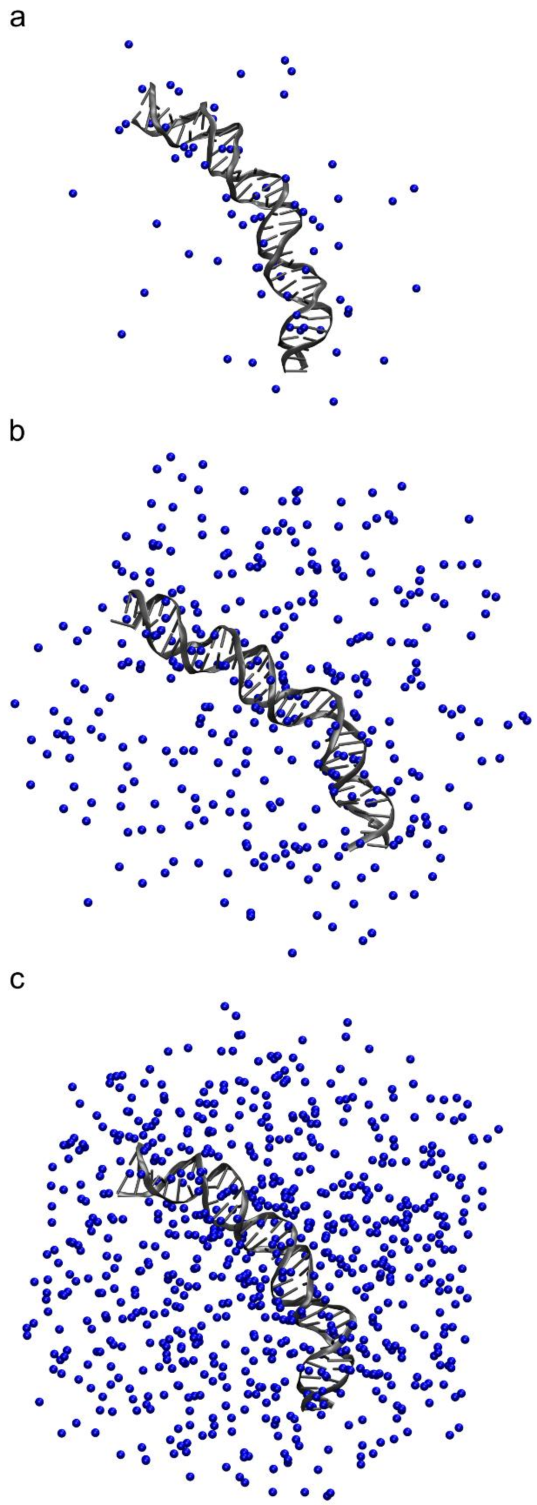

**Fig. S18. Figure 3 – MD simulation: snapshots.** Exemplary snapshots taken at time points of 1,715 ns at 50 mM sodium concentration (a), 1,327 ns at 250 mM sodium concentration (b), as well as 1,798 ns at 550 mM sodium concentration (c). The blue spheres represent sodium ions.

**Fig. S19.**

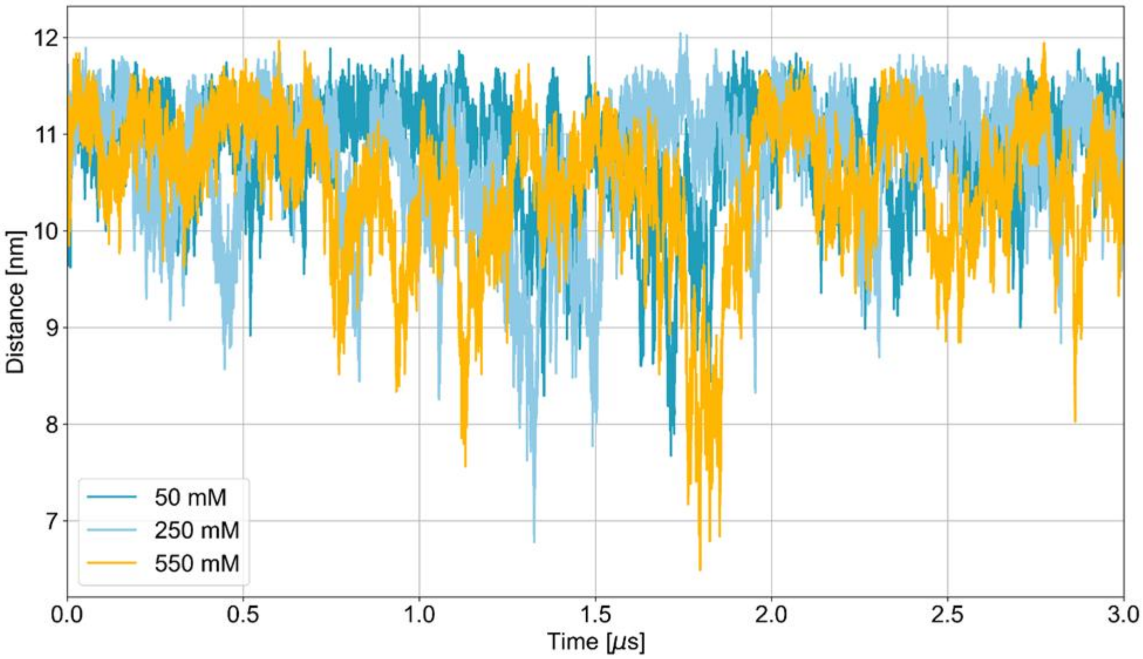

**Fig. S19. Figure 3 – MD simulation: time-distance traces.** The plot displays the raw simulated data for the projected z-distance obtained in the MD simulation for different sodium concentrations and with sequences according to Supplementary Table S1. For clarity, the raw data for the end-to-end distances are not shown. The respective histogram values can be found in Supplementary Figure S22.

**Fig. S20.**

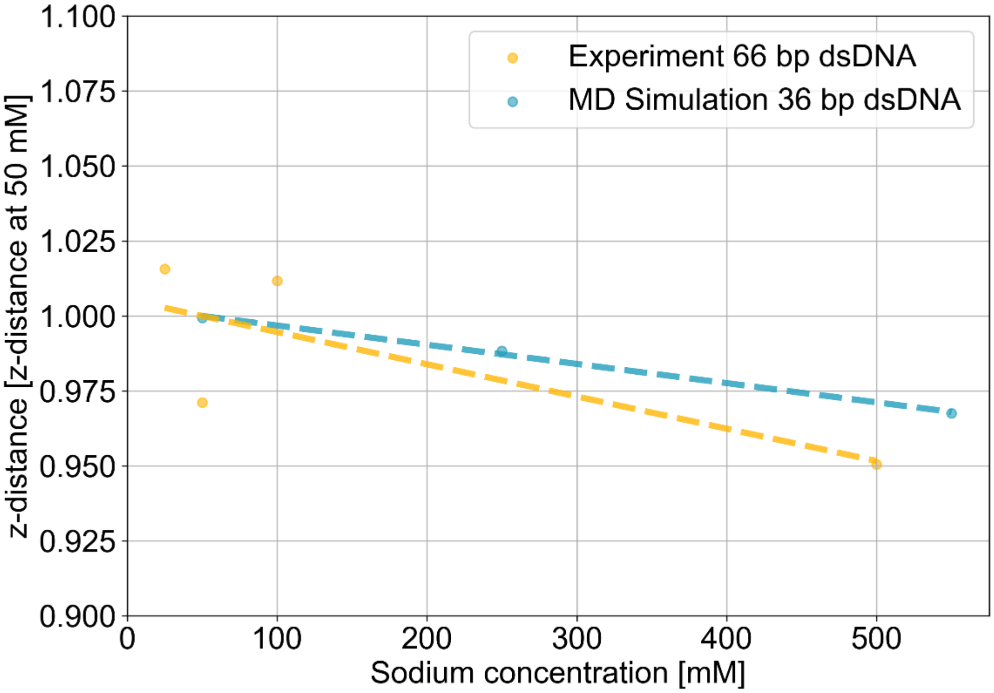

**Fig. S20. Figure 3 – Graphene distances from experiment versus MD simulation.** The plot displays the individual data points for the mean z-distance from experiments discussed in the main text (66 bp dsDNA) and the MD simulation (36 bp dsDNA) at the indicated sodium concentrations. The dashed lines provide the fit of the respective data points by a line. In order to compare the results, the values are normalized to the fitted value at 50 mM highlighting that the MD simulation results in a very similar dependency for the z-distance on the sodium concentration as the experiments.

**Fig. S21.**

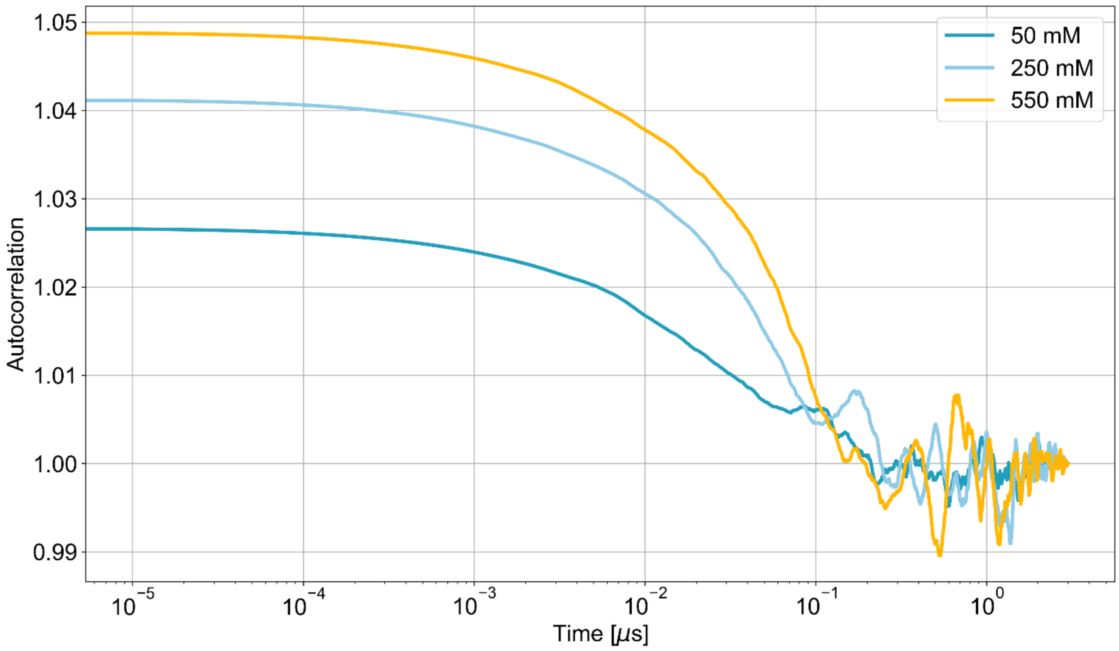

**Fig. S21. Figure 3 – MD simulation: correlation of simulated signal.** Based on the simulated distances, we calculated the respective normalized intensity on graphene and computed the corresponding second-order autocorrelation function. The resulting values are incremented by 1 for clarity. We note an increase in correlation amplitude as well as correlation time for higher sodium concentrations.

**Fig. S22.**

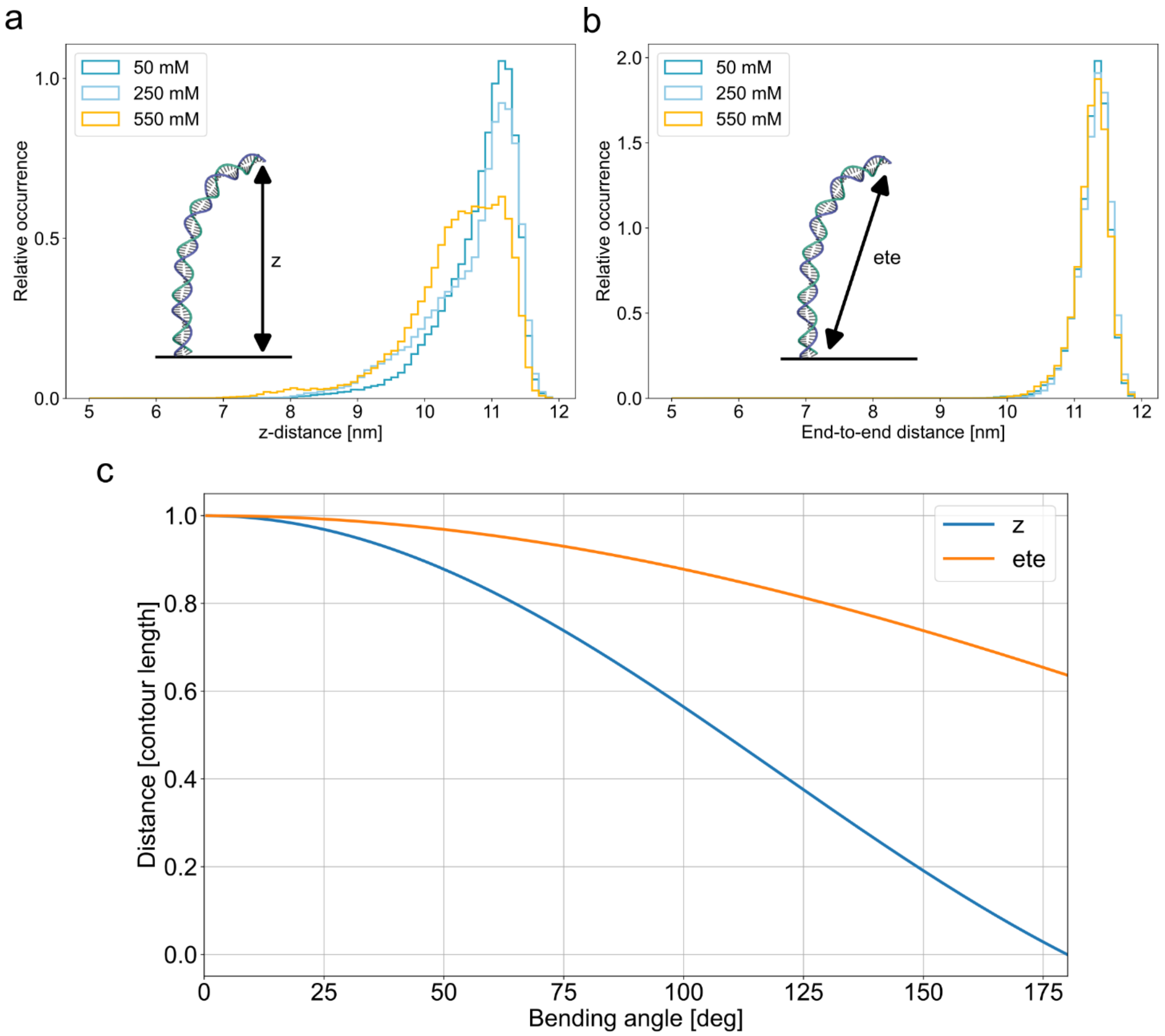

**Fig. S22. Figure 3 – MD simulation: end-to-end vs. projected z-distance.** Based on the raw data from the MD simulation of a 36 bp dsDNA at the three different sodium concentrations, we calculated the projected z-distance as well as the end-to-end (ete) distance. The histograms in (a) show the resulting *z*-distance whereas (b) displays the corresponding end-to-end distances. The comparison demonstrates that only the *z*-projection is sensitive to the minute fluctuations in distance caused by the different ionic environment. (c) Analytical calculation of the respective distance normalized to the contour length as a function of the bending angle. Similar to the model underlying the kMC simulation, we assume the Worm-like chain model and a quadratic approximation for the bending angle. This analysis supports the data from the MD simulation and underlines that the bending of the dsDNA does not cause a significant change in the ete distance (*e.g.* measured by FRET) compared to the change in distance for the *z*-projection.

**Fig. S23.**

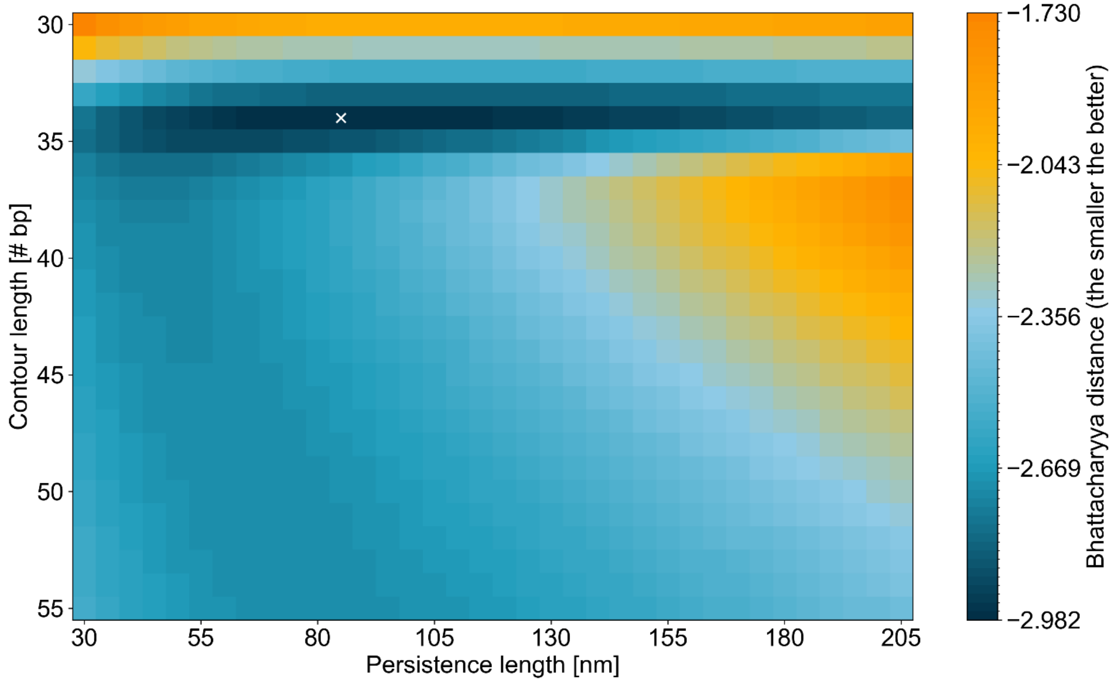

**Fig. S23. Figure 3 – MD simulation: comparison to kMC simulation**. In order to compare the distance distributions derived analytically (and underlying the kMC simulation) and from the MD simulation, we compute the respective Bhattacharyya distance (*3*). Whereas the MD simulation of the 36 bp dsDNA provides a fixed distance distribution for the defined dsDNA sequence and ion concentration, in our analytical model the contour length as well as the persistence length are two free parameters. The above graph shows the resulting Bhattacharyya distance for the corresponding combination of parameters. Each value is color-coded according to the color bar on the right. As indicated, smaller values show a higher agreement of the two distance distributions. Consequently, the optimum is marked by a white cross, corresponding to a persistence length of 85 nm and a contour length that corresponds to 34 bp dsDNA. The respective distance distributions are displayed in Figure S24.

The result justifies the choice of parameters in our kMC simulation. The longer persistence length will be mostly because of the limited sampling of extreme bending events in the 3 µs-long MD simulation compared to our experimental setting acquiring second-long signals. The optimized contour length results in a difference in distance that is smaller than the potential uncertainty in length due to the motion of the dye molecule. Furthermore, the MD simulations and the experiment were performed at two slightly distinct ion concentrations (50 mM vs. 100 mM) and the respective sequences were different by 2 nucleotides. Hence, we conclude that the MD simulation supports well the analytical model underlying the kMC simulation.

**Fig. S24.**

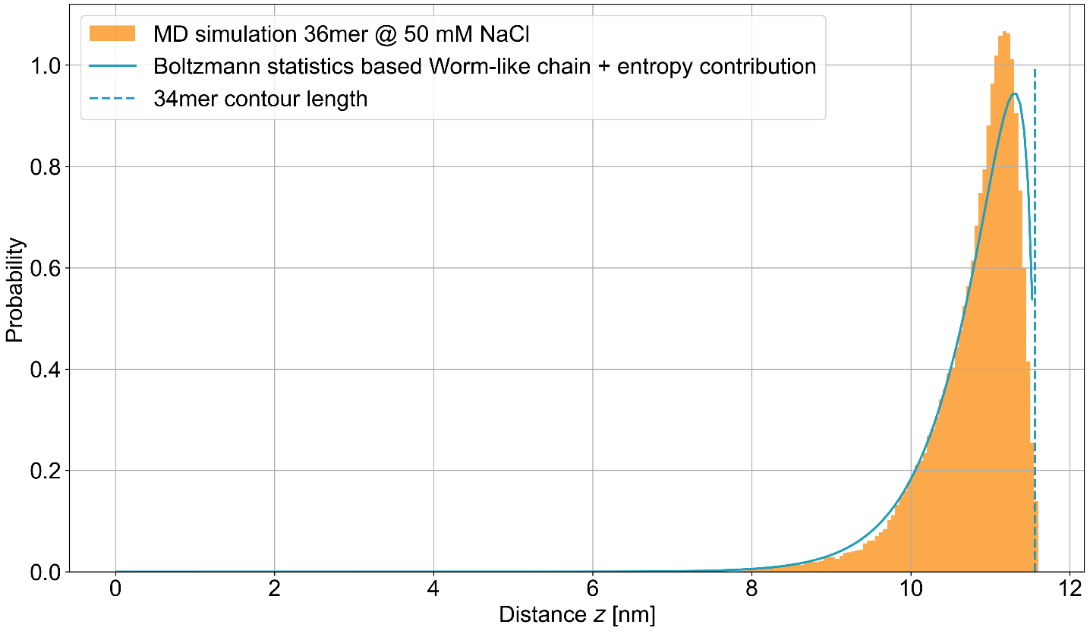

**Fig. S24. Figure 3 – MD simulation: probability distributions for optimized parameters.** See Supplementary Figure S23 for details.

**Fig. S25.**

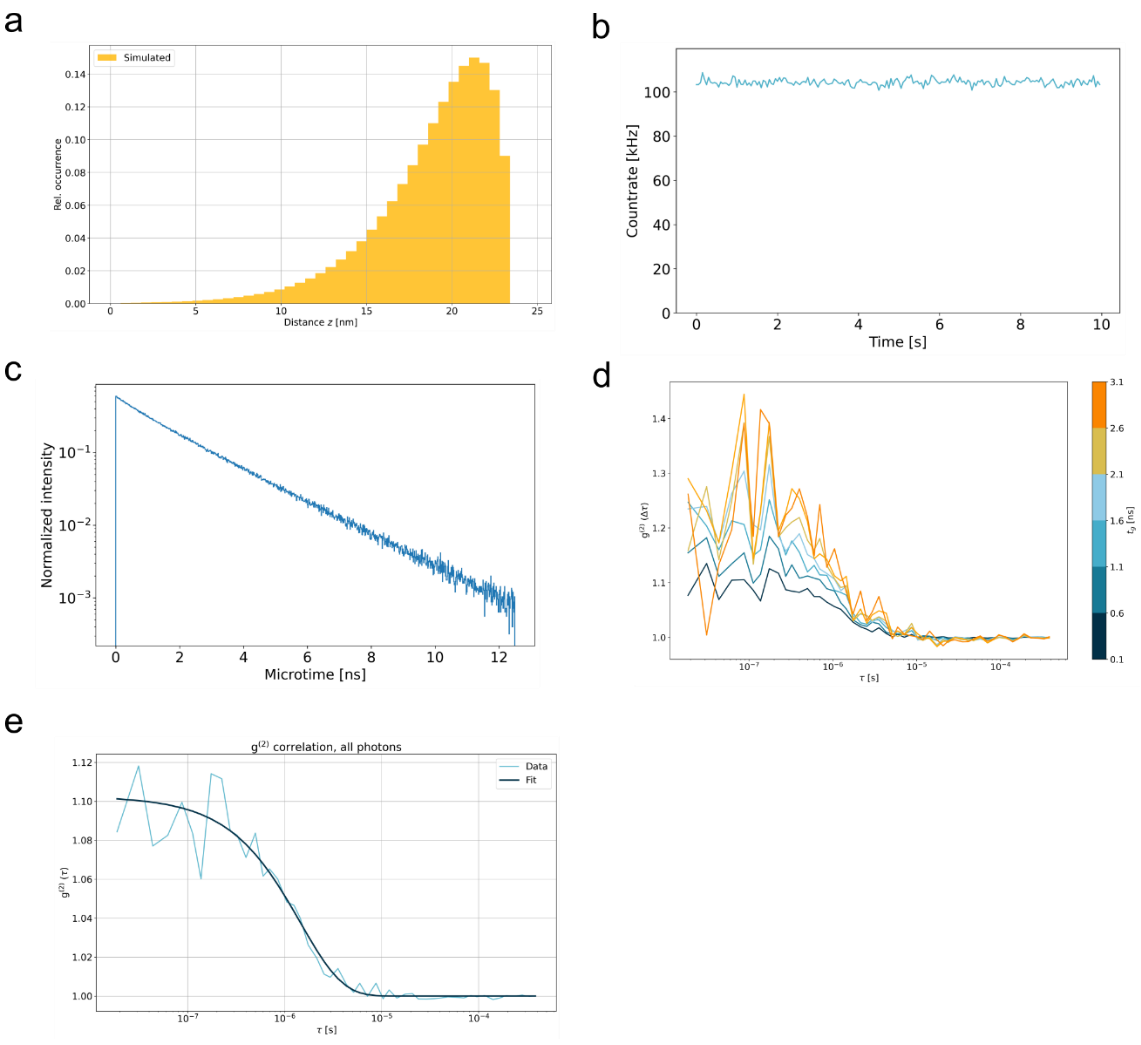

**Fig. S25. Figure 3 – kMC simulation: exemplary run.** Results from an exemplary run of the kinetic Monte Carlo simulation. The contour length of the dsDNA was set to equal a 66 bp dsDNA with a persistence length of 50 nm. Additionally, we accounted for 1.5 nm for the dye molecule and its DNA-linker, as well as another 0.4 nm for the distance to graphene. The photon emitter (*i.e.* dye molecule) was assumed to diffuse normally along the predefined probability distribution with a diffusion constant of 10 $\mu m^2/s$. (A) Resulting distribution showing the simulated motion and the discrete step size that was set to 0.6 nm. (B) The diffusion of the emitter was simulated for 10 s resulting in an intensity trace of roughly 100 kHz per 50 ms-time bin. (C) Histogram of the resulting microtimes. (D) Second-order autocorrelation of the simulated fluorescence intensity analysed using sgFCS, revealing dynamics. (E) Autocorrelation using all photon macrotimes. The respective monoexponential fit resulted in a value for the correlation amplitude of 0.10 and a correlation time of 1.45 µs.

**Fig. S26.**

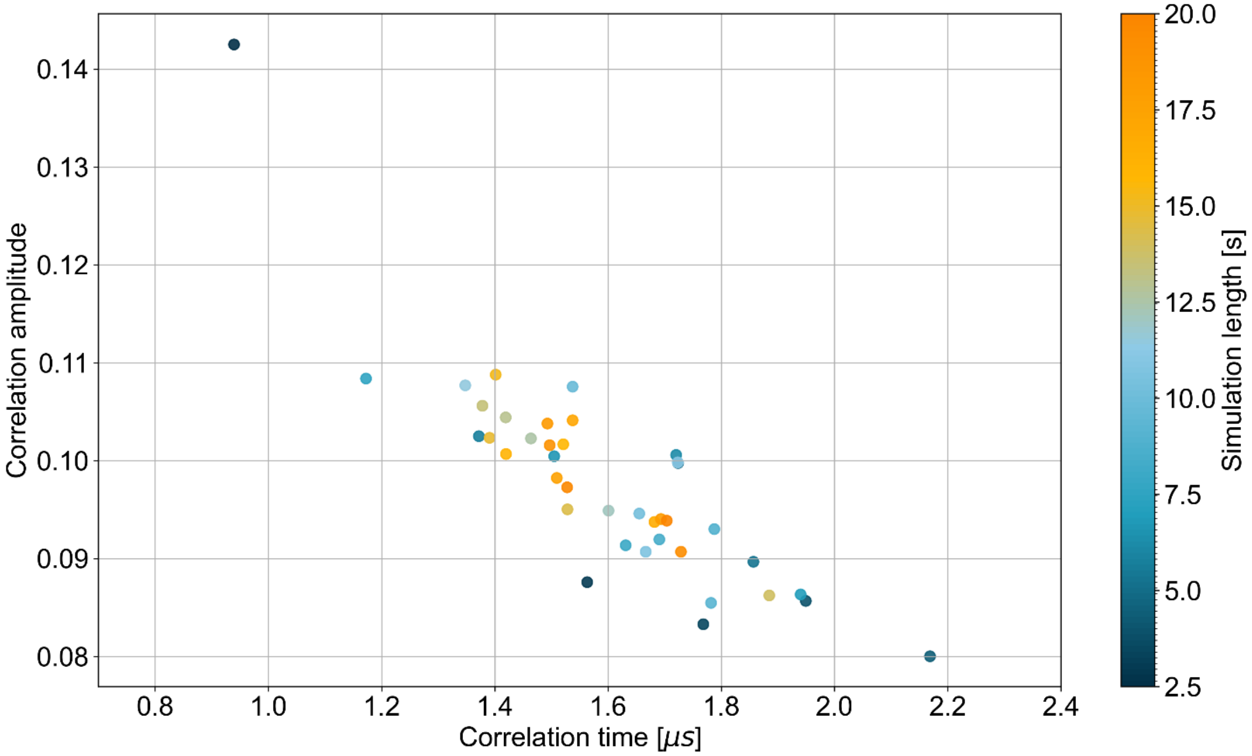

**Fig. S26. Figure 2 – kMC simulation: simulation results.** The described simulations for a 66 bp-long dsDNA (as exemplified in Supplementary Figure S25) were repeatedly run for different simulation lengths, that is experiment durations, assuming that the photon emitter is active within this duration. The resulting correlation amplitude as well as the corresponding correlation time from the fitted correlation function are displayed as colored dot. The color encodes the respective simulation length.

**Fig. S27.**

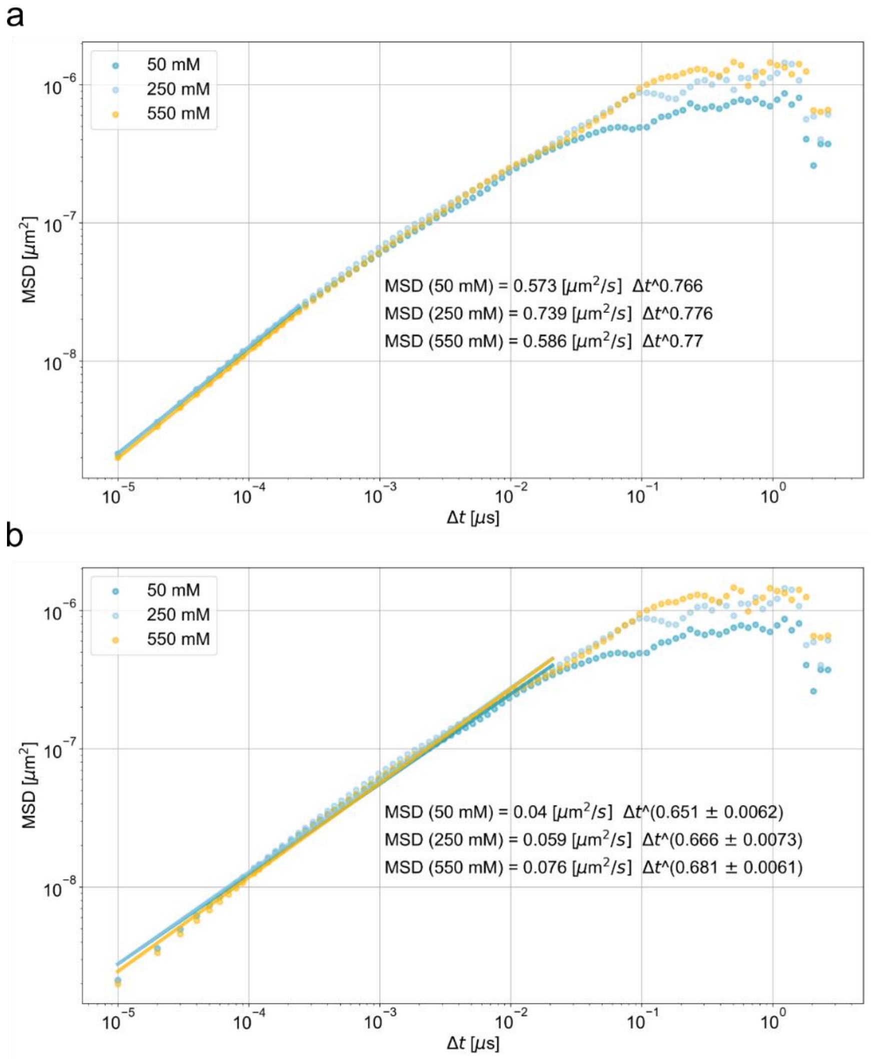

**Fig. S27. Figure 3 – MD simulation: diffusion at short and long timescales.** The mean square displacement (MSD) versus time lag *Δt* is analysed for 10 - 200 ps (a) as well as 10 ps - 20 ns (b). The respective time interval is fitted by a linear function, and the resulting dependency is annotated in each graph.

**Fig. S28.**

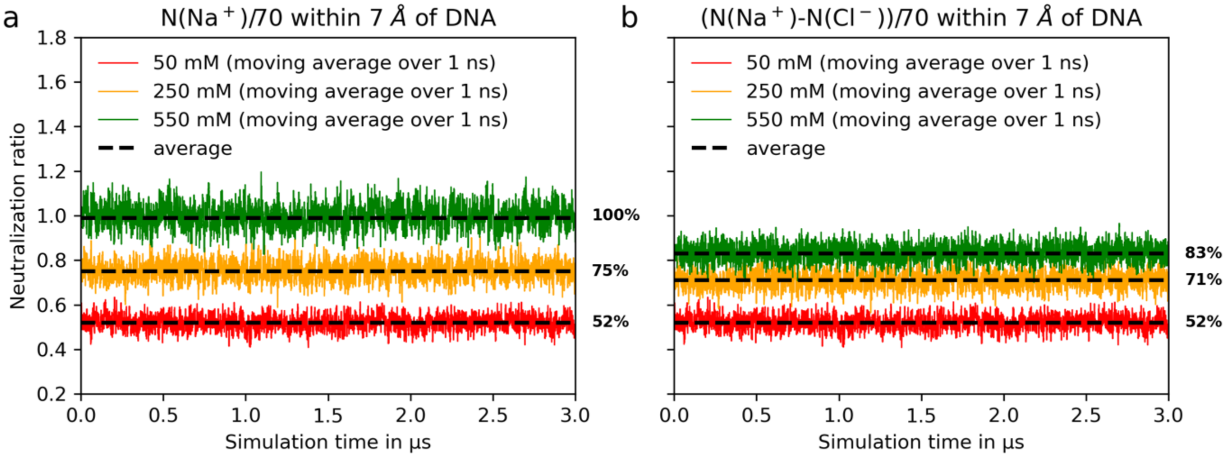

**Fig. S28. Figure 3 – MD simulation: analysis of ion concentrations.** Time evolution and average of the neutralization ratios within 7 Å to the surface of the DNA regarding only $Na^+$ cations (a) and corrected by the negative charge of present $Cl^-$ anions (b). See Figure S29 for the corresponding visualization of the ion shielding.

**Fig. S29.**

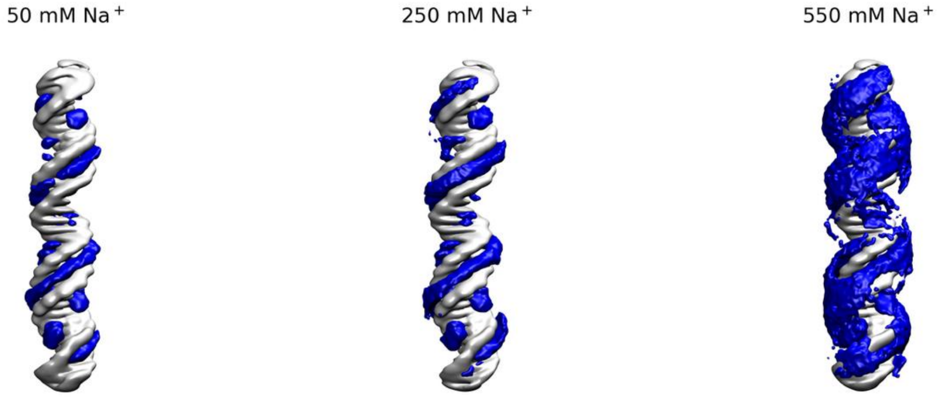

**Fig. S29. Figure 3 – MD simulation: ion density maps.** Average density maps of DNA (white) and $Na^+$ (blue) for the three investigated $Na^+$ concentrations in the DNA-water systems. The density maps were generated using the Volmap plugin of VMD and averaged over 3 µs with a step size of 0.1 ns. Isovalues of 0.5 and 0.02 $Å^{-3}$ were used for the visualization of DNA and $Na^+$ respectively.

**Fig. S30.**

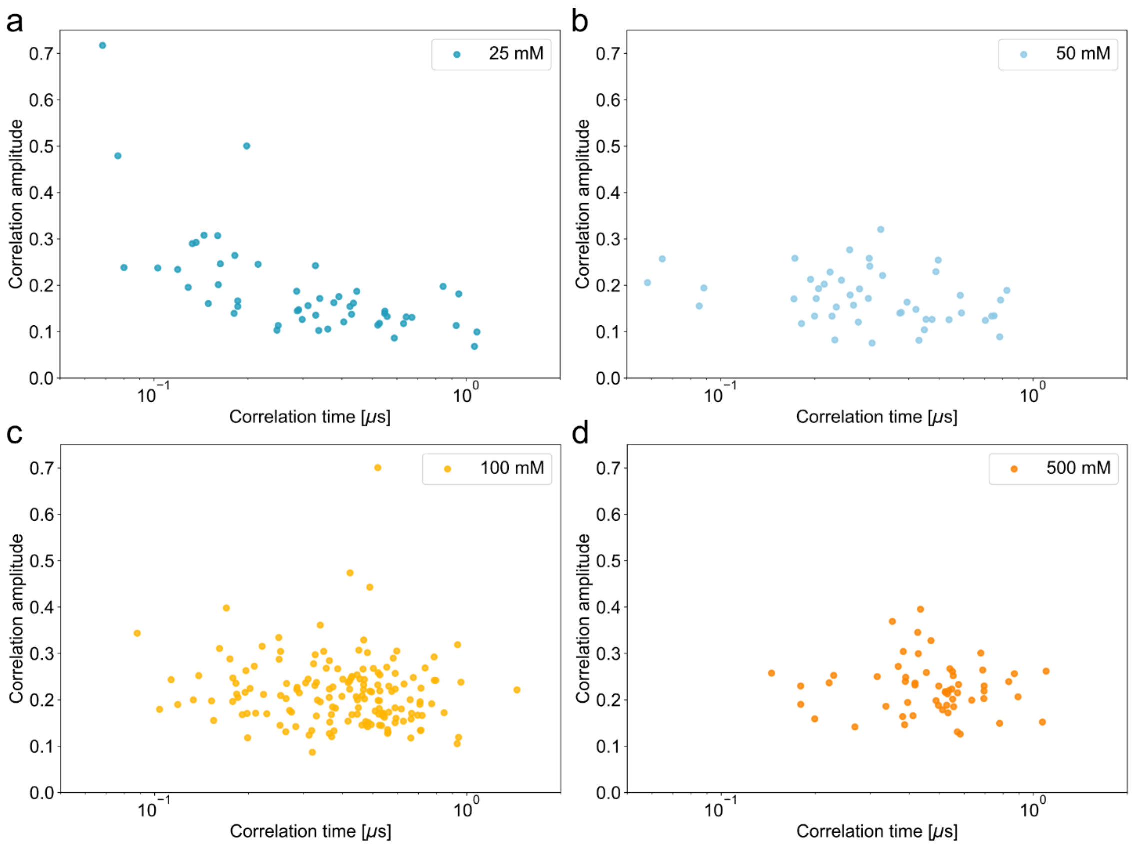

**Fig. S30. Figure 3 – Interdependency of correlation amplitude and correlation time.** Scatter plots displaying the experimentally determined correlation amplitudes and correlation times for individual data points at the indicated sodium chloride concentration.

**Fig. S31.**

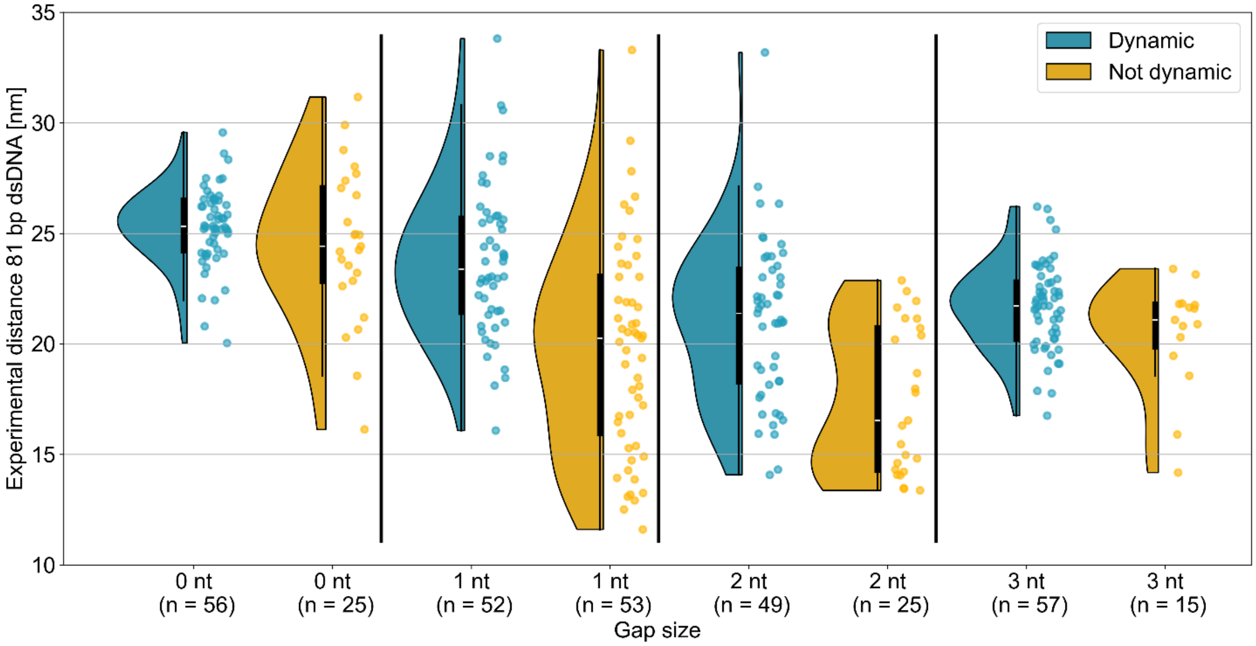

**Fig. S31. Figure 4 – Distances grouped by dynamics.** The plot displays all experimentally determined graphene distances for a 81 bp-long dsDNA at 0 nt to 3 nt gap size. The classification whether dynamic or not was done based on the respective shrinking-gate correlation. Each half violinplot on the left displays the distribution of the *n* datapoints as depicted by the dots on the right. The interior of the violinplot shows a box-and-whisker plot with the inter quartile range from the first to the third quartile shown as a black box separated by the median value in white. The whiskers range from minimum to maximum value. The results for each condition are separated by a vertical black line.

**Fig. S32.**

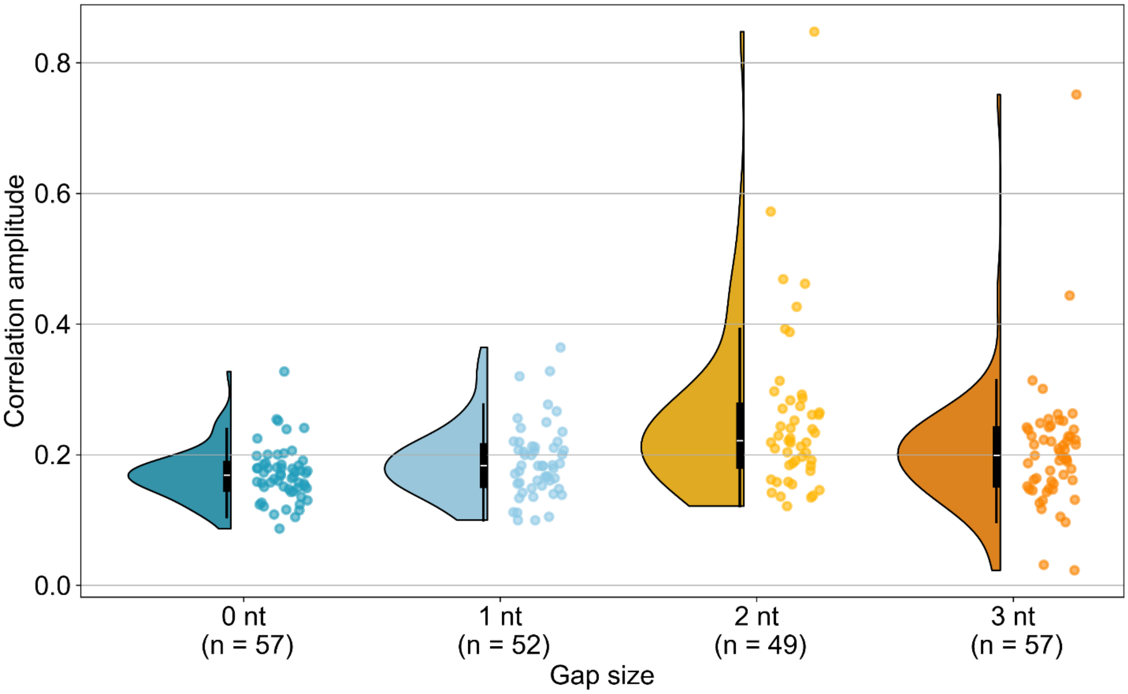

**Fig. S32. Figure 4 – Distribution of correlation amplitudes.** The plot displays all experimentally determined correlation amplitudes for a 81 bp-long dsDNA with 0 to 3 nt gap size. Each half violinplot on the left displays the distribution of the *n* datapoints as depicted by the dots on the right. The interior of the violinplot shows a box-and-whisker plot with the inter quartile range from the first to the third quartile shown as a black box separated by the median value in white. The whiskers range from minimum to maximum value.

**Fig. S33.**

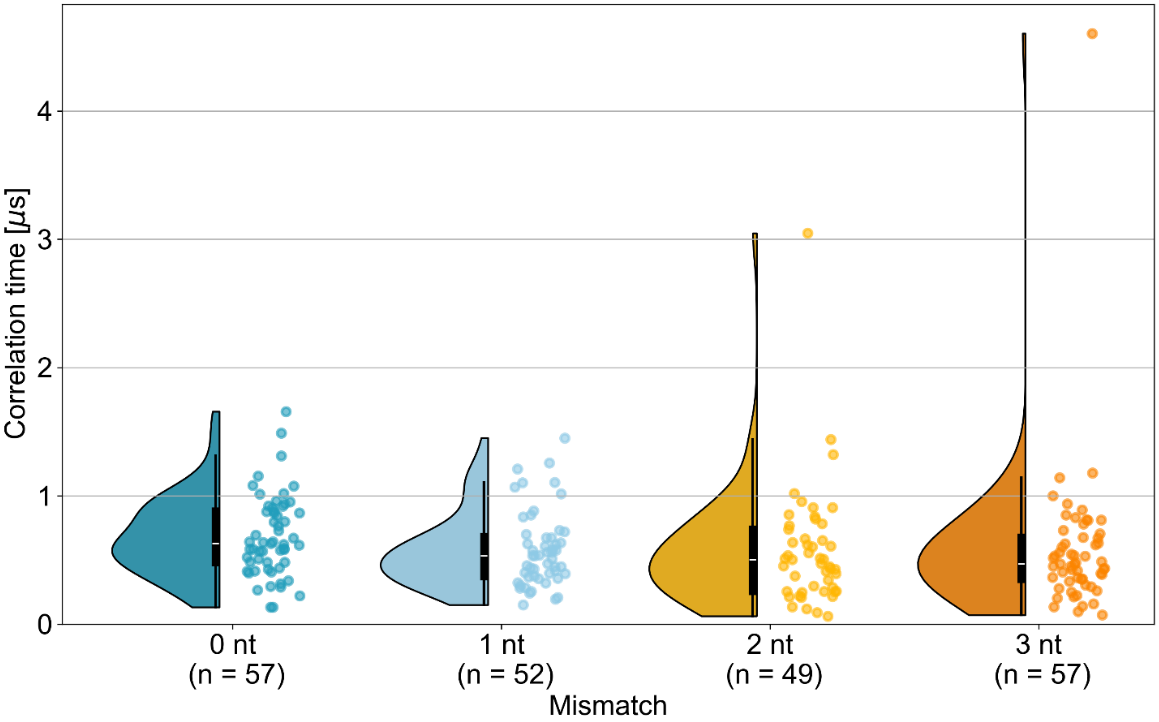

**Fig. S33. Figure 4 – Distribution of correlation times.** The plot displays all experimentally determined correlation times for a 81 bp-long dsDNA with 0 to 3 nt gap size. Each half violinplot on the left displays the distribution of the *n* datapoints as depicted by the dots on the right. The interior of the violinplot shows a box-and-whisker plot with the inter quartile range from the first to the third quartile shown as a black box separated by the median value in white. The whiskers range from minimum to maximum value.

**Fig. S34.**

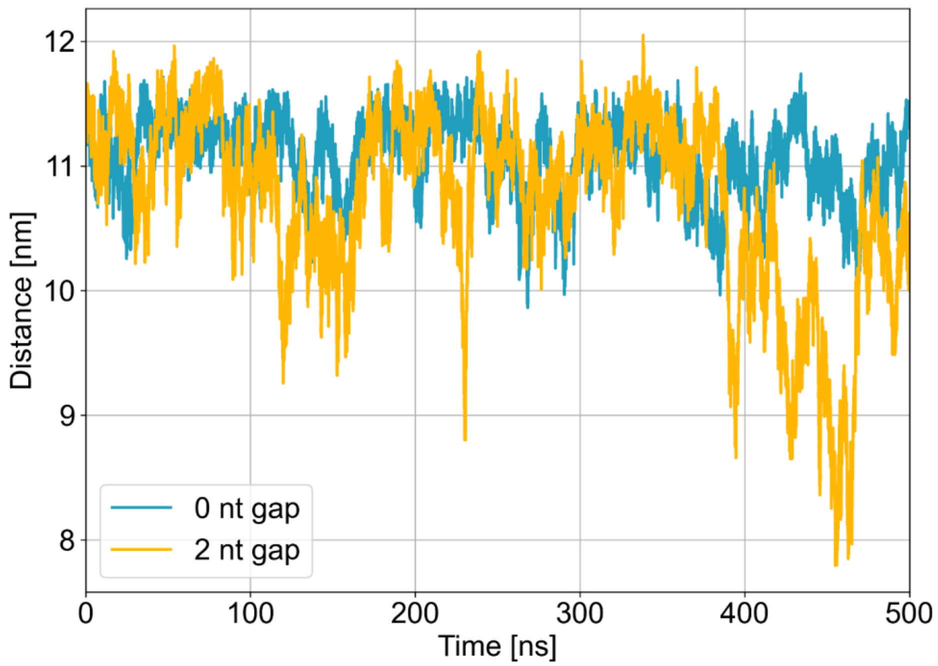

**Fig. S34. Figure 4 – MD Simulation: raw data with and without gap.** The plot displays the projected z-distance for 36 bp dsDNA with sequences according to Supplementary Table S1.

**Fig. S35.**

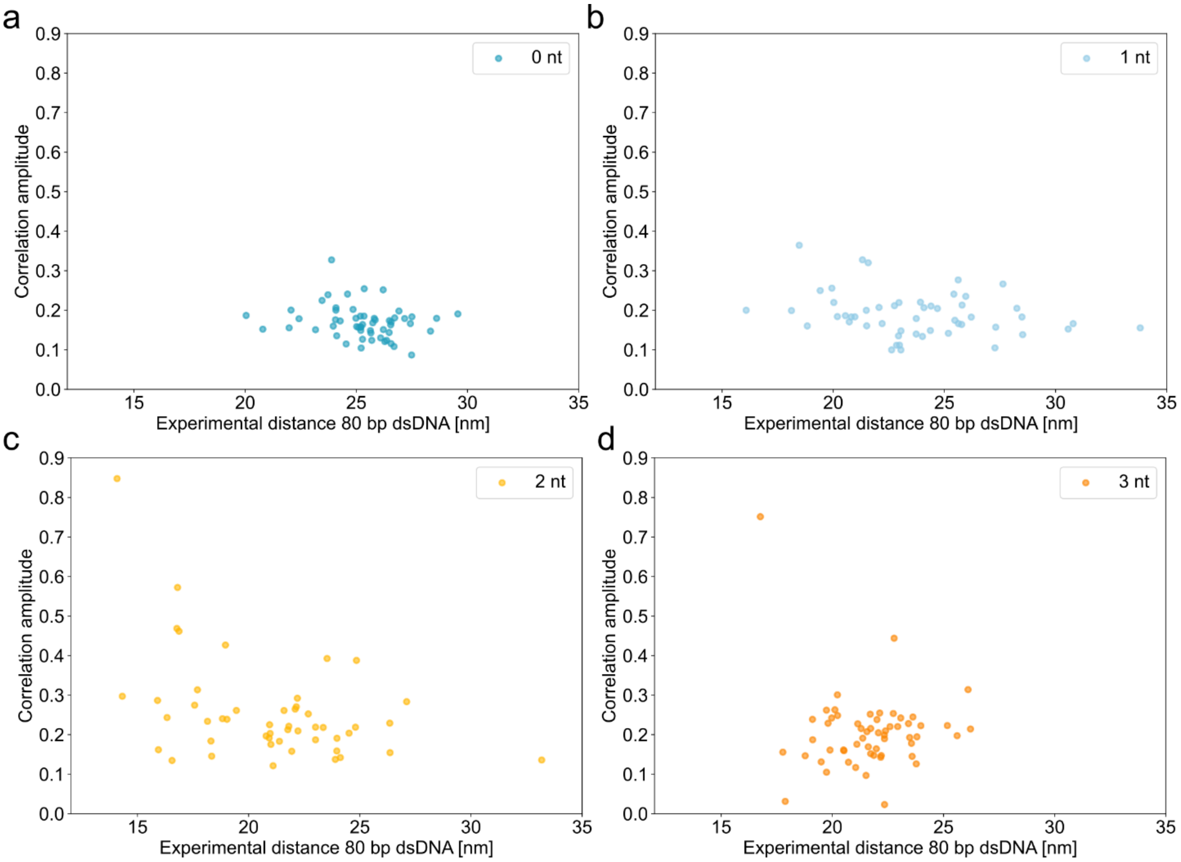

**Fig. S35. Figure 4 – Anticorrelation of correlation amplitude and distance.** Scatter plots displaying the experimentally determined correlation amplitudes versus the graphene distance for individual data points at the indicated gap size.

**Fig. S36.**

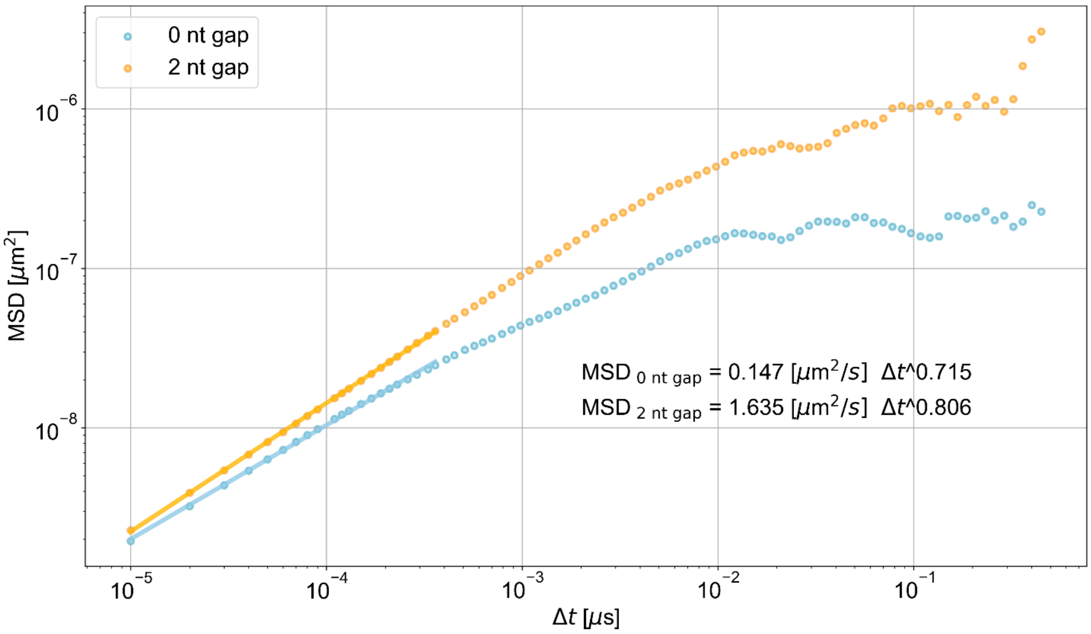

**Fig. S36. Figure 4 – MD Simulation: analysis of diffusion.** The mean square displacement (MSD) is plotted for different lag times Δ*t* for a 36 bp-long dsDNA bearing a 2 nt gap or no gap (0 nt). The respective data are fitted by a linear function resulting in the annotated diffusion equations.

**Fig. S37.**

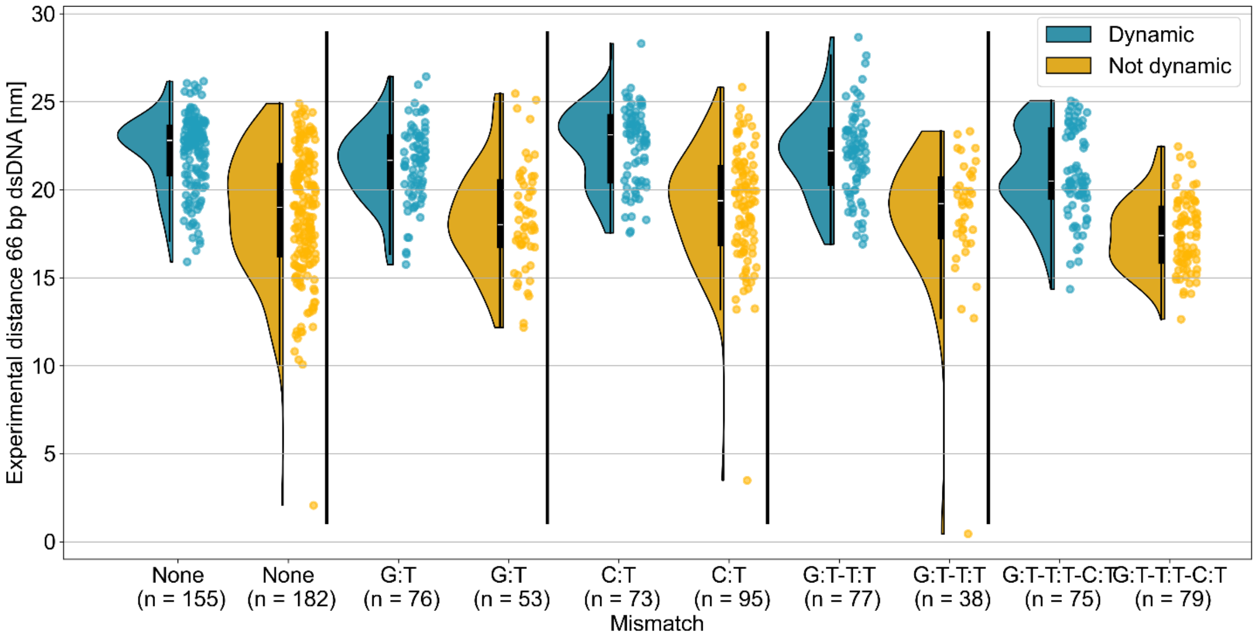

**Fig. S37. Figure 5 – Distances grouped by dynamics.** The plot displays all experimentally determined graphene distances for a 66 bp-long dsDNA for the indicated mismatch type. The classification whether dynamic or not was done based on the respective shrinking-gate correlation. Each half violinplot on the left displays the distribution of the *n* datapoints as depicted by the dots on the right. The interior of the violinplot shows a box-and-whisker plot with the inter quartile range from the first to the third quartile shown as a black box separated by the median value in white. The whiskers range from minimum to maximum value. The results for each condition are separated by a vertical black line.

**Fig. S38.**

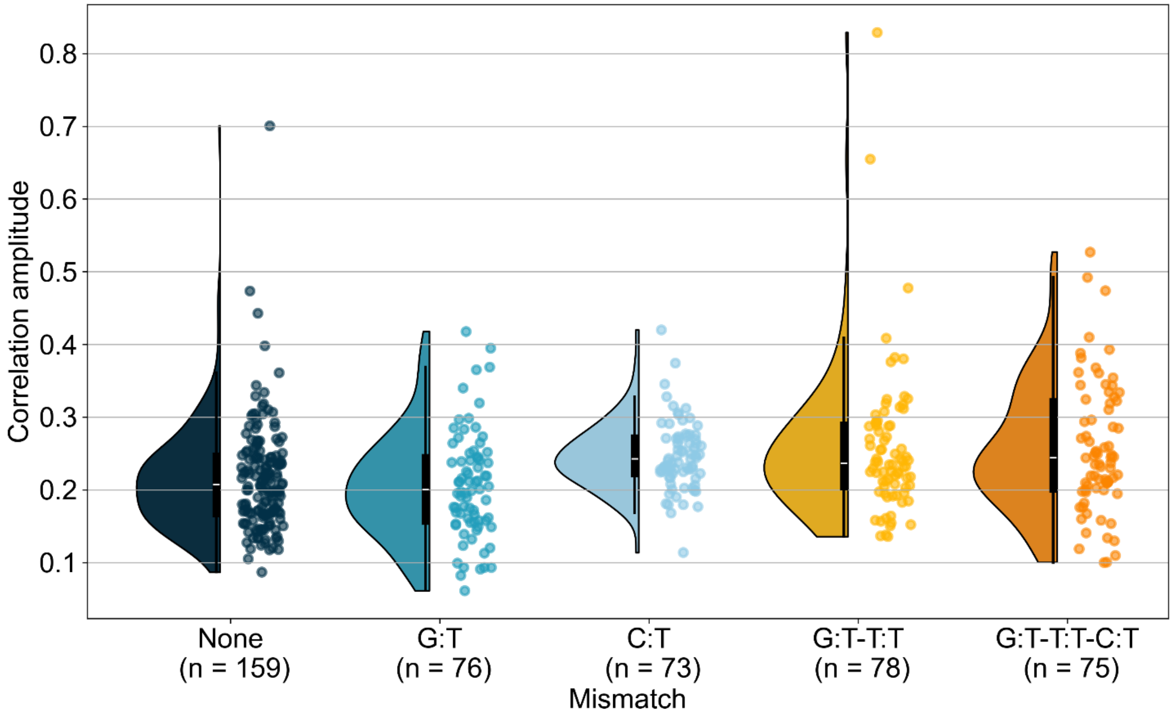

**Fig. S38. Figure 5 – Distribution of correlation amplitudes.** The plot displays all experimentally determined correlation amplitudes for a 66 bp-long dsDNA for the indicated mismatch types. Each half violinplot on the left displays the distribution of the *n* datapoints as depicted by the dots on the right. The interior of the violinplot shows a box-and-whisker plot with the inter quartile range from the first to the third quartile shown as a black box separated by the median value in white. The whiskers range from minimum to maximum value.

**Fig. S39.**

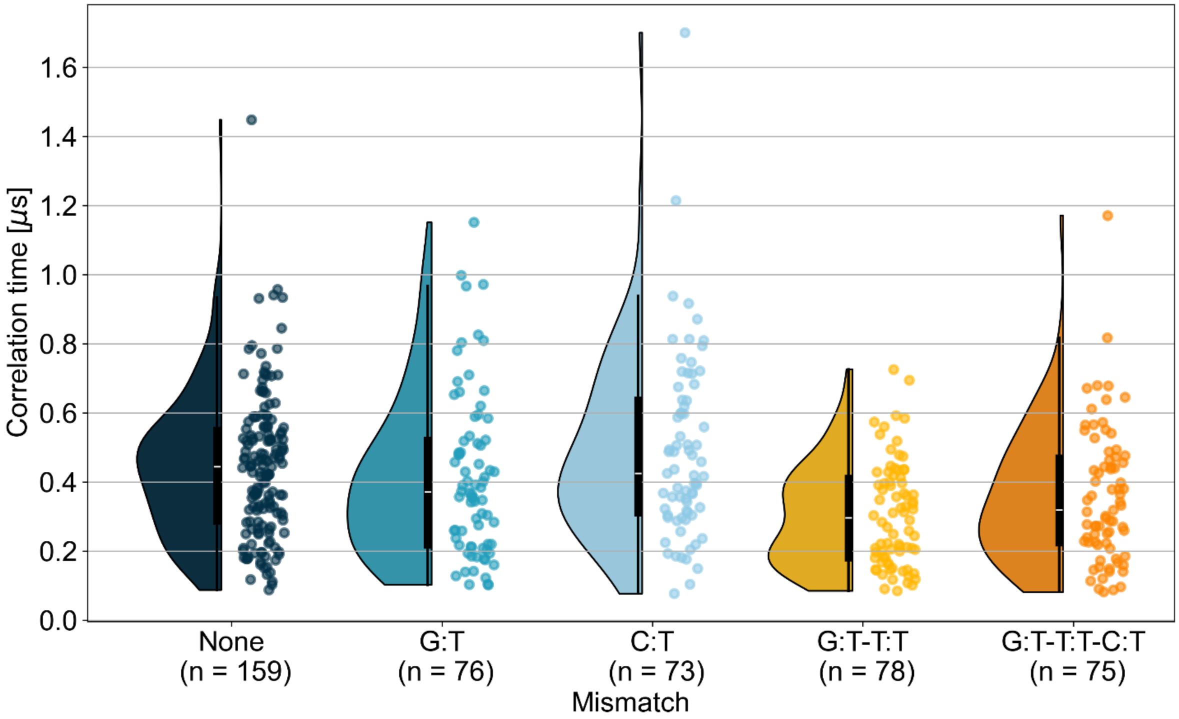

**Fig. S39. Figure 5 – Distribution of correlation times.** The plot displays all experimentally determined correlation times for a 66 bp-long dsDNA for the indicated mismatch type. Each half violinplot on the left displays the distribution of the *n* datapoints as depicted by the dots on the right. The interior of the violinplot shows a box-and-whisker plot with the inter quartile range from the first to the third quartile shown as a black box separated by the median value in white. The whiskers range from minimum to maximum value.

**Fig. S40.**

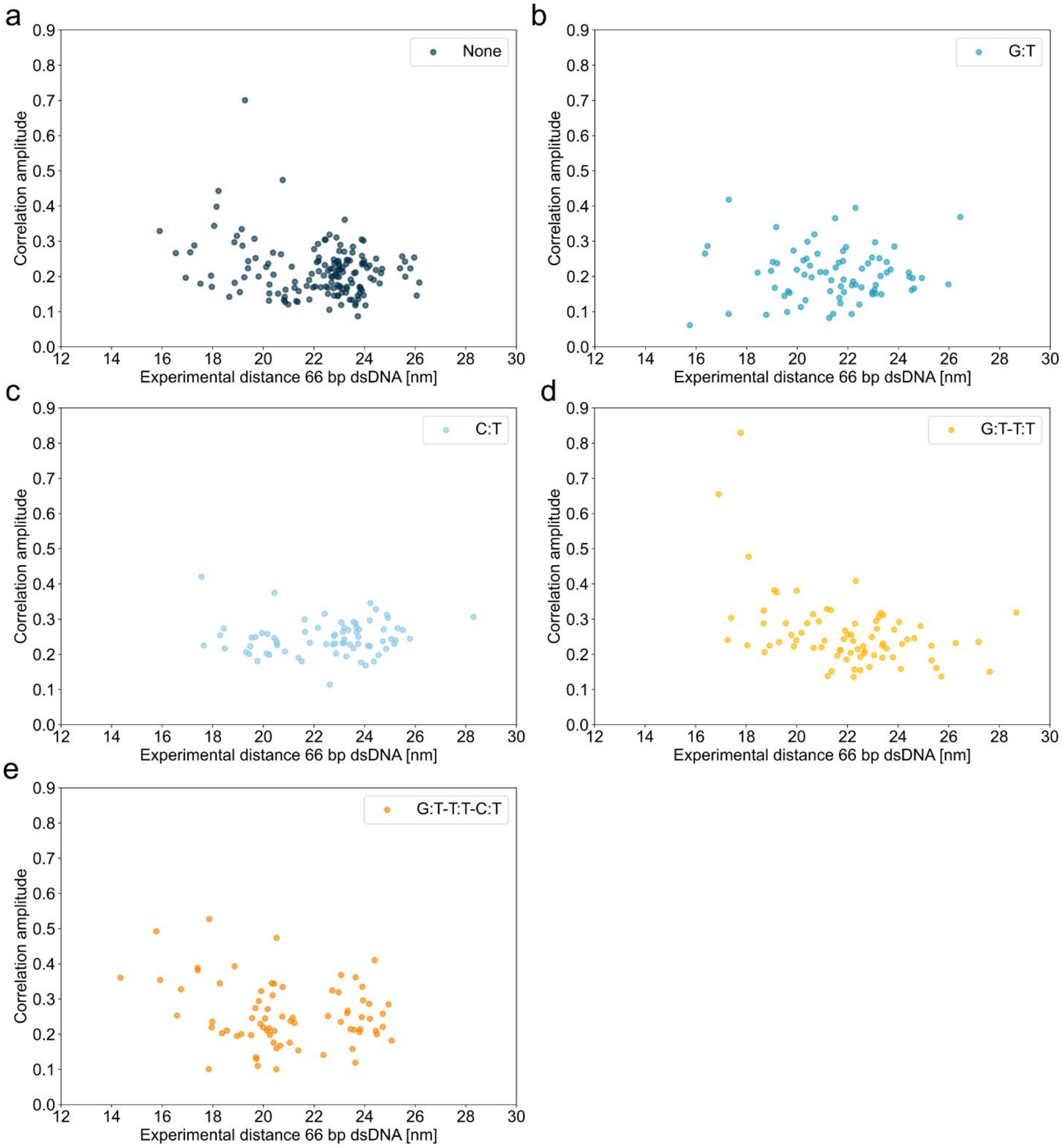

**Fig. S40. Figure 5 – Anticorrelation of correlation amplitude and distance.** Scatter plots displaying the experimentally determined correlation amplitudes versus the graphene distance for individual data points at the indicated gap size.

### 3 Supplementary Tables

**Table S1.**

List of DNA oligonucleotides for GETvNA experiments and MD simulations.

| Short description | Sequence (5' to 3' end) | Supplier |
|---|---|---|
| 96mer_A643 | GGATCTGTCGTGGTAACTTTGGGGTTGGATCGCGCTACAGTCATAAGAACTGGCTACGACGGACTGATCGGATCGAATAACGAGTACACGGTGATT[ATTO643] | Ella Biotech GmbH |
| 66mer | AATCACCGTGTACTCGTTATTCGATCCGATCAGTCCGTCGTAGCCAGTTCTTATGACTGTAGCGCG | Ella Biotech GmbH |
| 66mer_GT | AATCACCGTGTACTCGTTATTCGATCCGATCAGTCCGTCGTAGCCAGTTCTTATGATTGTAGCGCG | Ella Biotech GmbH |
| 66mer_CT | AATCACCGTGTACTCGTTATTCGATCCGATCAGTCCGTCGTAGCCAGTTCTTATGCCTGTAGCGCG | Ella Biotech GmbH |
| 66mer_GT_TT | AATCACCGTGTACTCGTTATTCGATCCGATCAGTCCGTCGTAGCCAGTTCTTATGTTTGTAGCGCG | Ella Biotech GmbH |
| 66mer_GT_TT_CT | AATCACCGTGTACTCGTTATTCGATCCGATCAGTCCGTCGTAGCCAGTTCTTATTTTTGTAGCGCG | Ella Biotech GmbH |
| 48mer_for_66_top | AATCACCGTGTACTCGTTATTCGATCCGATCAGTCCGTCGTAGCCAGT | IDT |
| 18mer_for_66/80_nick_bot | TCTTATGACTGTAGCGCG | IDT |
| 17mer_for_66/80_1gap_bot | CTTATGACTGTAGCGCG | IDT |
| 16mer_for_66/80_2gap_bot | TTATGACTGTAGCGCG | IDT |
| 110mer_A643 | GGATCTGTCGTGGTAACTTTGGGGTTGGATCGCGCTACAGTCATAAGAACTGGCTACGACGGACTGATCGGATCGAATAACGAGTACCAGGTGATTAGTACACGGTGATT[ATTO643] | Ella Biotech GmbH |
| 80mer | AATCACCGTGTACTAATCACCTGGTACTCGTTATTCGATCCGATCAGTCCGTCGTAGCCAGTTCTTATGACTGTAGCGCG | IDT |

| Short description | Sequence (5’ to 3’ end) | Supplier |
|---|---|---|
| 62mer_for_80_top | AATCACCGTGTACTAATCACCTGGTACTCGTTATTCGATCCGATCAGTCCGTCGTAGCCAGT | IDT |
| A | TAACCACCGTGTACTCGTTATTCGATCCGATCAGTC | - |
| B1 | CAGTCCGTCGTAGCCAGTTCTTATGACTGTAGCGCG | - |
| B2.1 (top) | CAGTCCGTCGTAGCCAGT | |
| B2.2 (bottom) | TTATGACTGTAGCGCG | - |

**Table S2.**

Average helical parameters, end-to-end and z-distances for the tree investigated DNA-water systems (3 µs). Major groove width, minor groove width and rise parameters were averaged over all bases, excluding the terminating two base pairs.

| System | Minor groove width / Å | Major groove width / Å | Rise / Å | End to end distance / Å | z-distance / Å |
|---|---|---|---|---|---|
| 50 mM DNA-water | 12.3 ± 0.5 | 19.5 ± 0.8 | 3.3 ± 0.3 | 112.7 ± 2.5 | 108 ± 6 |
| 250 mM DNA-water | 12.3 ± 0.5 | 19.5 ± 0.8 | 3.3 ± 0.3 | 112.9 ± 2.3 | 107 ± 7 |
| 550 mM DNA-water | 12.3 ± 0.5 | 19.5 ± 0.8 | 3.3 ± 0.3 | 112.6 ± 2.6 | 105 ± 8 |

**Table S3.**

Summary of the MD system setups. The total simulation time is indicated, and results from the sum of equilibration time and simulation time. Sequences as in Supplementary Table S1.

| System | Sequence 5’-3’ | $Na^+$ Concentration / mM | $Na^+$ Equivalents | Total Simulation Time / ns |
|---|---|---|---|---|
| 50 mM DNA-water | A | 50 | 1 | 5425 |
| 250 mM DNA-water | A | 250 | 5 | 4911 |
| 550 mM DNA-water | A | 550 | 11 | 4641 |
| DNA-graphene-water | A | 25 | 1 | 1085 |
| Healthy DNA-water | B1 | 100 | 2 | 1515 |
| 2 nt-gap DNA-water | B2.1, B2.2 (+ B1 as reverse complement) | 100 | 2 | 1515 |

**Table S4.**

Average helical parameters, end to end and z-distances for the 2nt-gap DNA-water and corresponding healthy DNA-water systems (500 ns). Major groove width, minor groove width and rise parameters were averaged over all bases, excluding the terminating two base pairs and unpaired bases.

| System | Minor groove width / Å | Major groove width / Å | Rise / Å | End to end distance / Å | z-distance / Å |
|---|---|---|---|---|---|
| Healthy DNA-water | 12.3 ± 0.5 | 19.5 ± 0.8 | 3.3 ± 3.3 | 113.1 ± 1.8 | 111 ± 3 |
| 2 nt gap-DNA-water | 12.3 ± 0.6 | 19.4 ± 0.9 | 3.3 ± 3.3 | 113 ± 3 | 107 ± 7 |

**Table S5.**

List of oligonucleotides for DNA origami nanostructure.

| Name | Sequence (5' to 3' end) | 5'-end | 3'-end |
|---|---|---|---|
| S1 | ATCGATGCTGAGAGTCTACAAGGAGAGGGAACGCCAAAAGGA [ATTO643] | 26[76] | 10[63] |
| S2 | ACCAACAAACCAAAATTAACAATTTCATTTGAATTACCGAGG | 2[65] | 4[80] |
| S3 | CCTAATTTAACAAACCCTCAATCAATATCTGATTCGCTAATC | 0[128] | 4[115] |
| S4 | ACGAGCGGCGCGGTCAGGCAAGGCGATTAAGTTGGGTAAAAC | 4[184] | 0[199] |
| S5 | CAAGCCCAATAGGAACCACCCTCACCCGGAA | 33[18] | 43[34] |
| S6 | ACTAAAGAGCAACGTGAAAATCTCCACCCACAACTAAAGGAA | 44[48] | 33[48] |
| S7 | AGAGAAAATCCAGAGAGTTGCAGCAAATC | 0[325] | 3[318] |
| S8 | ACATAAGTAGAAAAATCAAGAAGCAAAAGAAGATGTCAT | 5[70] | 47[66] |
| S9 | CGCTTTCCAGTTAGCTGTTTAAAGAACGT | 4[318] | 0[304] |
| S10 | AGGGAGCCGCCACGGGAACGGATAGGCGAAAGCATCAGCACTCTG | 0[268] | 2[280] |
| S11 | GATTAGAGAGTACCTTAACTCCAACAGG | 15[12] | 16[16] |
| S12 | AGTTTCCAACATTATTACATTATAC | 31[56] | 47[52] |
| S13 | TGAGTGTTCCGAAAGCCCTTCACCGCCTAGGCGGTATTA | 0[349] | 4[339] |
| S14 | AATTTCTTAAACCCGCTTAATTGTATCGTTGCGGGCGATATA | 37[49] | 40[49] |
| S15 | AGAACTTAGCCTAATTATCCCAAGCCCCCTTATTAGCGTTTGCCA | 63[49] | 51[79] |
| S16 | TCACAGCGTACTCCGTGGTGAAGGGATAGCTAAGAGACGAGG | 2[226] | 0[213] |
| S17 | CATCGAGATAACGTCAAACATAAAAGAGCAAAAGAATT | 53[25] | 58[35] |
| S18 | GACAATTACGCAGAGGCATTTTCGAG | 61[21] | 62[9] |
| S19 | TTAACTCGGAATTAGAGTAAATCAATATATGTGAGTGATTCT | 2[107] | 4[122] |
| S20 | GAACCGCCACCCTCCATATCATACC | 48[76] | 2[66] |
| S21 | GGATGTGGTTTGCCCCAGCAG | 0[303] | 1[318] |
| S22 | ACATAAAGCCCTTACACTGGTCGGGTTAAATTTGT | 7[252] | 6[245] |

| Name | Sequence (5' to 3' end) | 5'-end | 3'-end |
|---|---|---|---|
| S23 | CGAGGGTACTTTTTCATGAACGGGGTCATAATGCCGAGCCACCACC | 43[35] | 34[17] |
| S24 | TTGCGAATAATATTTACAGCGGAGTGAGGTAAAATTTTGAGG | 33[49] | 44[49] |
| S25 | AAGACAAATCAGCTGCTCATTCAGTCTGACCA | 38[80] | 41[76] |
| S26 | GAGTTAAAAGGGTAATTGAGCGCTAATATCAGAGGAACTGAACACC | 58[34] | 56[45] |
| S27 | TTTTCCAGCATCAGCGGGGCTAAAGAACCTCGTAGCACGCCA | 5[235] | 3[220] |
| S28 | AATAAAACGAACTATGACCCCACCAAGC | 30[83] | 46[70] |
| S29 | GAGTCTGGATTTGTTATAATTACTACATACACCAC | 2[58] | 48[52] |
| S30 | AATACCCCAACATTCATCAAAAATAATTCGCGTCT | 23[56] | 19[76] |
| S31 | CCGACTTGTTGCTAAAATTTATTTAGTTCGCGAGAGTCGTCTTTCCAGA | 48[27] | 9[34] |
| S32 | GAGCATTTATCCTGAATCAAACGTGACTCCT | 49[46] | 64[35] |
| S33 | CTAGTCAGTTGGCAAATCAACAGTCTTTAGGTAGATAACAAA | 5[109] | 3[94] |
| S34 | TTTACCAGTCCCGGCCTGCAGCCCACTACGGGCGCACCAGCT | 6[265] | 10[252] |
| S35 | TTTCCATGGCACCAACCTACGTCATACA | 45[35] | 46[21] |
| S36 | TATTACGAATAATAAACAAATCAGATATGCGT | 64[34] | 49[24] |
| S37 | AACAACAGGAAGCACGTCCTTGCTGGTAATATCCAGAAACGC | 0[177] | 2[192] |
| S38 | CCCAGCTACAATGACAGCATTTGAGGCAAGTTGAGAAATGAA | 7[35] | 9[48] |
| S39 | CGGAATAGAAAGGAATGCCTTGCTAAACAACTTTCAAC | 45[46] | 31[55] |
| S40 | TAGCCTCAGAGCATACCCTGT | 22[62] | 23[55] |
| S41 | CAGCAGCGCCGCTTGTTTATCAGCTTCACGAAAAA | 42[48] | 35[41] |
| S42 | AAATGCGGAAACATCGGTTTTCAGGTTTAACGTCAGATTAAC | 0[114] | 2[129] |
| S43 | CCTCGTCTTTCCACCACCGGAACCGCCTCCCTCA | 48[51] | 49[79] |
| S44 | CCTTAAATCAAGATTAGCGGGAGGCTCAAC | 7[5] | 65[20] |
| S45 | TAACATCCAATAAATGCAAAGGTGGCATCAACATTATGAAAG | 19[39] | 22[45] |
| S46 | AGAATTTTAGAGGAAAACAATATTACCGCCAGCTGCTCATTT | 0[198] | 2[213] |
| S47 | GCTGTAGTTAGAGCTTAATTG | 11[21] | 12[14] |

| Name | Sequence (5' to 3' end) | 5'-end | 3'-end |
|---|---|---|---|
| S48 | TTAGCCCTGACGAGAAACACCAGAAATTGGGGTGAATTATTTTAA | 34[65] | 32[77] |
| S49 | AGGCTTGCGAGACTCCTCAAGAGAAAAGTATTCGGAAC | 40[37] | 42[14] |
| S50 | CTGGCATTAGGAGAATAAAATGAAGAAACGATTTTTTGAGTA | 56[44] | 61[38] |
| S51 | CCAGCCTCCGATCCTCATGCCGGA | 2[212] | 9[216] |
| S52 | TGCTGATTGCCGTTGTCATAAACATCGGGCGG | 8[258] | 4[255] |
| S53 | AGAAAACGAGAATGACCATAAATCTACGCCCCTCAAATGCTTTA | 23[9] | 24[17] |
| S54 | AGGACAGATGAACGGTGTAACATAAGGGAACCGAAGAAT | 40[76] | 36[66] |
| S55 | TAAGTTTACACTGAGTTTCGT | 45[21] | 32[14] |
| S56 | AACAAGAGCCTAATGCAGAACGCGC | 57[21] | 58[10] |
| S57 | GAATTCGTCTCGTCGCTGGGTCTGCAATCCATTGCAACACGG | 3[221] | 1[206] |
| S58 | CTTGTAGAACGTCAGCGGCTGATTGCAGAGTTTTTCGACGTT | 9[217] | 11[230] |
| S59 | AATTGTGTCGAAATCCGCGGCACACAACGGAGATTTGTATCA | 44[76] | 45[76] |
| S60 | AACCGTGTCATTGCAACGGTAATATATTTTAAATGAAAGGGT | 14[69] | 27[55] |
| S61 | AGAAATCGTTAGACTACCTTTTTAAGGCGTTCTGACCTTTTTGCA | 4[37] | 7[34] |
| S62 | TACGGCTGGAGGTGCGCACTCGTCACTGTTTGCTCCCGGCAA | 10[251] | 8[238] |
| S63 | GCCCGAGTACGAGCCGGAAGC | 0[363] | 5[370] |
| S64 | ATAGCGAGAGGCTATCATAACCAAATCCCAAAGAAAATTTCATCCTCAT | 29[21] | 8[28] |
| S65 | TGCATTAATGAGCGGTCCACGCTCACTGCGCCACGTGCCAGC | 3[305] | 3[304] |
| S66 | TGGCTTTTTACCGTAGAATGGAAAGCG | 46[20] | 8[8] |
| S67 | CCGTAATCAGTAGCGACAGAATCTAATTATTCATTAAAAAGG | 54[79] | 60[66] |
| S68 | GAGAACAATATACAAAATCGCGCAGAGGCGATTCGACAAATCCTTTAAC | 4[114] | 1[101] |
| S69 | TAAAGCCTCCAGTACCTCATAGTTAGCG | 8[27] | 30[14] |

| Name | Sequence (5' to 3' end) | 5'-end | 3'-end |
|---|---|---|---|
| S70 | AGTACCGCATTCCACAACATGTTCAGCCTTAAGGTAAAGTAATTC | 53[14] | 60[11] |
| S71 | TGGTGGTTGTTCCAGTTTGGAACA | 1[336] | 0[326] |
| S72 | ATTACGAGATAAATGCCAGCTTTGAGGGGACGACGACAG | 10[62] | 13[73] |
| S73 | CCCCGCTAGGGCAACAGCTGGCGAAAGGGGGATGTGCTTATT | 4[163] | 0[178] |
| S74 | AAATGACGCTAAATGGATTATTTACATTGGCGAATACCTGGA | 4[198] | 2[185] |
| S75 | ACCGCCACCCTCAGAACCCGTACTCTAGGGA | 35[18] | 41[34] |
| S76 | GTTAAAGGAAAGACAGCATCTGCCTATTTAAGAGGCAGGAGGTTTA | 41[35] | 36[17] |
| S77 | TTGGTAGAACATTTAATTAAGCAAC | 9[49] | 5[59] |
| S78 | GAGAAGGCATCTGCAATGGGATAGGTCAAAAC | 27[56] | 16[66] |
| S79 | CCTCATCACCCCAGCAGGCCTCTTCGCTATTACGCCAGTGCC | 4[142] | 0[157] |
| S80 | AGCTTTCAGAGGTGGCGATGGCCAGCGGGAAT | 11[252] | 2[262] |
| S81 | TCGTGCCGGAGTCAATAGTGAATTTGCAGAT | 12[72] | 5[69] |
| S82 | CTGTATGGGATTACCGTTAGTATCA | 30[48] | 4[38] |
| S83 | GCGAAACAAAGTGTAAAACACATGGCCTCGATTGAACCA | 46[69] | 2[59] |
| S84 | TGCTAAATCGGGGAGCCCCCGATTTAGAGCTAGCAGAACATT | 0[212] | 4[199] |
| S85 | TATCAGCAACCGCAAGAATGCCAATGAGCCTGAGGATCTATC | 4[254] | 0[269] |
| S86 | ATTGCGTTGCTGTTATCCGCTCACAATTCCAAACTCACTTGCGTA | 4[338] | 3[349] |
| S87 | TAGCCAGCTTTCATCCAAAAATAAACGT | 18[76] | 22[63] |
| S88 | TATGACTTTATACATTTTTTTTTAATGGAAACAGTACACCGT | 2[86] | 4[101] |
| S89 | TTCATCGGCATTTTCGGTCATATCAAAA | 52[79] | 61[59] |
| S90 | TAAAACCGTTAAAGAGTCTGTCCATCCAGAAACCACACAATC | 2[142] | 0[129] |
| S91 | CCTCGTTTACCAGAAACCAAA | 10[27] | 29[20] |
| S92 | CCAATGTTTAAGTACGGTGTCCAAC | 13[49] | 29[45] |
| S93 | CAAAATCACCGGAACCAGAGCCAGATTTTGTCACAATCACAC | 50[79] | 64[66] |

| Name | Sequence (5’ to 3’ end) | 5'-end | 3'-end |
|---|---|---|---|
| S94 | TCATACATTTAATACCGATAGCCCTAAAACATCGAACGTAAC | 2[191] | 4[206] |
| S95 | GCTGGCATAGCCACATTATTC | 29[53] | 4[52] |
| S96 | ATTTCCTGATTATCAGATGATGGCTTTAAAAAGACGCTAAAA | 1[102] | 5[87] |
| S97 | AAACTCACAGGAACGGTACGCCAGTAAAGGGGGTGAGGAAC C | 1[207] | 5[192] |
| S98 | AATATGCAACTACCATCATAGACCGGAACCGCCGCGCCGCCA | 11[42] | 7[55] |
| S99 | TTTAGCGATACCAACGCGTTA | 49[25] | 50[38] |
| S100 | AGTTTATTGTCCATATAACAGTTGATTC | 14[34] | 13[48] |
| S101 | AGCTCTTACCGAAGCCCAATA | 61[39] | 59[31] |
| S102 | GCGAATCAGTGAGGCCACCGAGTAGTAGCAACTGAGAGTTGA | 1[165] | 5[150] |
| S103 | AAGAAAGCGCTGAACCTCAAATATTCTAAAGGAAAGCGTTCA | 5[151] | 3[136] |
| S104 | GAAGGAGCGGAATTATCATCATATATCATTTACATAGCACAA | 1[123] | 5[108] |
| S105 | AAGGGATATTCATTACCGTAATCTATAGGCT | 36[65] | 39[76] |
| S106 | CAAAGCACTAGATAGCTCCATTCAGGCTGCGCAACTGTCTTG | 4[100] | 0[115] |
| S107 | ATAACTATATGTAAATGCTTAGGATATAAT | 5[5] | 11[20] |
| S108 | ATTTCAACCAAAAATTCTACTAATAGTTAGTTTCATTTGGGGCG CGAGC | 24[41] | 17[48] |
| S109 | TAATATCAAAGGCACCGCTTCTGGCACT | 5[60] | 12[66] |
| S110 | CTCATCGGGATTGAGTGAGCGAGTAACAACCCGTC | 25[56] | 17[76] |
| S111 | TGAGCAAATTTATACAGGAATAACATCACTTGCCTGAGTCTT | 0[135] | 2[150] |
| S112 | GGAACCATACAGGCAAGGCAAATCAAAAAGACGTAGTAGCAT | 20[58] | 19[38] |
| S113 | AAAGATTACAGAACGGGAGAAGGAAACGTCACCAATGAAACC A | 59[51] | 55[79] |
| S114 | CGTACAGGCCCCCTAACCGTCCCCGGGTACCGAGCGTTC | 6[279] | 4[283] |
| S115 | CACGGCAACAATCCTGATATACTT | 64[65] | 1[69] |
| S116 | AGGAATCATTACCGCGTTTTTATAAGTACC | 51[5] | 61[20] |
| S117 | CATTTCGCAAATGTCATCTGCGAACGAGAGATTCACAATGCC | 15[42] | 26[42] |
| S118 | TGAAAATCCGGTCAATAACCTAAATTTTAGCCTTT | 17[49] | 24[42] |

| Name | Sequence (5’ to 3’ end) | 5'-end | 3'-end |
|---|---|---|---|
| S119 | ACGGGCCGATAATCCTGAGAAGTGTTTTTATGGAGCTAACCG | 2[184] | 0[171] |
| S120 | TGCCCGTATAAACAGTGTGCCTTCTGGTAA | 43[5] | 45[20] |
| S121 | CGTACTATGGTAACCACTAGTCTTTAATGCGCGAACTGAATC | 2[170] | 4[185] |
| S122 | TATTGAAAGGAATTGAGGTAG | 5[88] | 29[80] |
| S123 | TTCGGTCCCATCGCATAGTTGCGCCGACATGCTTTCGAGGTG | 40[48] | 37[48] |
| S124 | TTACCATTAGCAAGGCCTTGAATTAGAGCCAGCCCGACTTGAGC | 56[74] | 58[59] |
| S125 | TTTTTGCGGATGCTCCTAAAATGTTTAGATGAATTTTGCAAAAGAAGTT | 13[18] | 28[17] |
| S126 | TTTAGATTCACCAGTCACACGACCGGCGCGTGCTTTCCCAGA | 3[179] | 1[164] |
| S127 | TTGCACGTAAAACAACGT | 7[77] | 2[87] |
| S128 | GGGATATTGACGTAGCAATAGCTAAGATAGC | 61[60] | 61[48] |
| S129 | AGCGCAGCTCCAACCGTAATCATGGTCACGGGAAACCT | 1[294] | 4[298] |
| S130 | GCTGGTCTGGTCAGGAGCCGGAATCCGCCGTGAACAGTGCCA | 9[238] | 11[251] |
| S131 | TTGGGCGGCTGATTTCGGCAAAATCCCT | 3[350] | 1[363] |
| S132 | GCATGTAGAAACCAATCCATCCTAGTCCTG | 55[5] | 57[20] |
| S133 | CAAATTATTCATTTCAATTACCTGAGTA | 3[95] | 3[80] |
| S134 | ACAACGCCTGTAGCATTTACCGTATAGGAAG | 31[18] | 45[34] |
| S135 | TAGCCCGGAATAGGTGTAAGGATAAGTGCCGTCGA | 37[18] | 38[17] |
| S136 | GTCGCAGAAAAACTTAAATTTGCC | 0[219] | 7[223] |
| S137 | CCAGAACAGAGCCATAAAGGTGGAA | 7[56] | 64[59] |
| S138 | GAACTGGCTCATTACAACTTTAATCATTCTTGAGATTACTTA | 32[76] | 43[76] |
| S139 | TTAGTTTGAGTGCCCGAGAAATAAAGAAATTGCGTAGAGATA | 0[93] | 2[108] |
| S140 | ATTTGGAAGTTTCATGCCTCAACATGTTTTA | 29[39] | 11[41] |
| S141 | GGCGCAGACGGTCAATCATCGAGACCTGCTCCATGTGGT | 42[76] | 34[66] |
| S142 | TGACCGCGCCTTAATTTACAATATTTTTGAATGGCTATCACA | 2[149] | 4[164] |
| S143 | AATATCGTTAAGAGAGCAAAGCGGATTGTGAAAAATCAGGTCTTT | 19[21] | 22[18] |

| Name | Sequence (5’ to 3’ end) | 5'-end | 3'-end |
|---|---|---|---|
| S144 | CATTTGAGATAACCCACGAAACAATG | 58[58] | 59[50] |
| S145 | CAAGCCGCCCAATAGCAAGTAAACAGCCATATTATTTTGCCATAAC | 52[27] | 49[45] |
| S146 | TAAGTTGGCATGATTAAAGAA | 64[58] | 65[45] |
| S147 | TAACGACATTTTTACCAGCGCCAAAGAAAGTTACCAGAACCCAAA | 60[65] | 63[48] |
| S148 | CTGAATATAGAACCAAATTAT | 1[70] | 7[76] |
| S149 | AATATTCATTGAATCCATGCTGGATAGCGTCCAAT | 25[18] | 26[17] |
| S150 | ACCTGACGGGGAAAGCCGGCGAACCAAGTGTCTGCGCGTTGC | 5[193] | 3[178] |
| S151 | AAGGCTCCAAAAGGAGCCTTTATATTTTTTCACGTGCTACAGTCACCCT | 35[42] | 42[49] |
| S152 | CGAACACCAAATAAAATAGCAGCCAAGTTTGCCTTTAGCGTCAGA | 61[49] | 53[79] |
| S153 | TTCGGGGTTTCTGCCAGGCCTGTGACGATCC | 4[282] | 1[293] |
| S154 | GCGTCCACTATTCCTGTGTGAAATGCTCACTGCC | 1[319] | 4[319] |
| S155 | AAGGCCTGTTTAGTATCATGTTAGCTACCTC | 1[39] | 48[28] |
| S156 | CCATAATGCCAGGCTATCAAGGCCGGAGACATCTA | 12[65] | 29[52] |
| S157 | GTAATTAATTTAGAATCTGGGAAGGGCGATCGGTGCGGCAAA | 4[121] | 0[136] |
| S158 | GTCGCGTGCCTTCGAATTGTCAAAG | 4[297] | 0[287] |
| S159 | ACTAATGCCACTACGAATAAA | 47[53] | 45[45] |
| S160 | CCTGCGCTGGGTGGCGAGAAAGGAAGGGAAGGAGCGGGGCCG | 0[170] | 4[157] |
| S161 | GAGGCCAAGCTTTGAATACCAAGTACGGATTACCTTTTCAAA | 3[137] | 1[122] |
| S162 | AAATCAGCTCATTTTTTAACCATTTTGTTAAAATTCGCATTA | 20[83] | 21[83] |
| S163 | GCGAAAATCCCGTAAAAAAAGCCGTGGTGCTCATACCGGCGTCCG | 2[261] | 4[273] |
| S164 | GGCAACACCAGGGTCTAATGAGTGAGCTCACAACAATAGGGT | 2[363] | 0[350] |
| S165 | TGCCATCCCACGCAGGCAGTTCCTCATTGCCGTTTTAAACGA | 10[272] | 8[259] |

| Name | Sequence (5’ to 3’ end) | 5'-end | 3'-end |
|---|---|---|---|
| S166 | CTAAATCGGTCAGAATTAGCAAAATTAAGCAATAAAATAATA | 22[44] | 20[59] |
| S167 | GCCAGCAGTTGGGCGCAAATCAGGTTTCTTGCCCTGCGTGGT | 7[224] | 9[237] |
| S168 | TGCGTGTTCAGGTTGTGTACATCG | 1[242] | 7[251] |
| S169 | TATTTAAATTGCAGGAAGATTG | 22[80] | 23[80] |
| S170 | GGCCAACGCGCGGGGAGGCCCTGTGTTTGA | 3[319] | 1[335] |
| S171 | ATAAAGTCTTTCCTTATCACT | 59[32] | 53[24] |
| S172 | TTATAAGGGTATGGAATAATTCATCAATATA | 3[81] | 6[70] |
| S173 | ATTAGCGGGGTTTTGCTCAGTACCAGGCTGACAACAAGCTG | 39[5] | 40[38] |
| S174 | AGTAGGTATATGCGTTATACA | 65[21] | 6[14] |
| S175 | ACGCGGTCCGTTTTTGGGTAAGTGA | 8[237] | 2[227] |
| S176 | ATTGTTATCTGAGAAGAAACCAGGCAAAGCGCCATTCGTAGA | 4[79] | 0[94] |
| S177 | GGCGAAGCACCGTAATAACGCCAGGGTTTTCCCAGTCATGGG | 4[205] | 0[220] |
| S178 | CAAACGGAATAGGAAACCGAGGAATAAGAAATTACAAG | 50[37] | 52[28] |
| S179 | ATCGGTCAGATGATATTCACAAACCAAAAGA | 4[51] | 47[48] |
| S180 | CTTACGGAACAGTCAGGACGTTGGGAAGAAA | 47[67] | 31[83] |
| S181 | AATAGAAAAAAATAAACGTCTGAGAGGAATATAAGAGCAACAC TATGAT | 65[46] | 29[38] |
| S182 | ATGAAGGGTAAAGTTCACGGTGCGGCCATGCCGGTCGCCAT G | 8[279] | 6[266] |
| S183 | TGAGTAAAGGATAAGTTTAGCTATATCATAGACCATTAGATA | 26[41] | 15[41] |
| S184 | GTAAAACGACGGCCCATCACCCAAATCAGCGC | 11[23 1] | 5[234] |
| S185 | GAGAGATAGACTTTACGGCATCAGA | 6[244] | 1[241] |
| S186 | AAGAAAGCTTGATACCGCCACGCATACAGACCAGGCGCTGAC | 38[65] | 38[66] |
| S187 | AGCAACAAAGTCAGAAATAATATCCAATAATCGGCTCAGGGA | 55[39] | 55[38] |
| S188 | CCGTGTGATAAATAACCTCCGGCTGATG | 1[14] | 5[27] |
| S189 | GGCTAAAACTTCAGAAAAGTTTTGCGGGAGATAGAACC | 16[65] | 25[55] |
| S190 | ACGTAAGAATTCGTTCTTAGAAGAACTCAAACTATCGGATAA | 0[156] | 2[171] |

| Name | Sequence (5’ to 3’ end) | 5'-end | 3'-end |
|---|---|---|---|
| S191 | CGTGTCAAATCACCATCTAGGTAATAGATTT | 29[46] | 14[35] |
| S192 | AGTCGCCTGATACTTGCATAACAGAATACGTGGCACAGCTGA | 2[128] | 4[143] |
| S193 | CGCGCTACAGAGTAATAAAAGGGACATTCTGATAGAACTTAG | 4[156] | 2[143] |
| S194 | CCCGGTTGATAAAGCATGTCAATC | 24[81] | 25[81] |
| S195 | ACGCGAGAGAAGGCCATGTAATTTAGGCCAGGCTTAATTGAGAATCGC | 49[14] | 64[8] |
| S196 | AGACAGCAGAAACGAAAGAGGAAATAAATCGAGGTGACAGTTAAAT | 30[34] | 1[38] |
| S197 | ACCAGACCGGATTAATTCGAGC | 17[17] | 18[17] |

**Table S6.**

| Temperature [°C] | Time [s] |
|---|---|
| 65 | 300 |
| 64 | 50 |
| 63 | 95 |
| 62 | 95 |
| 61 | 95 |
| 60 | 540 |
| 59 | 540 |
| 58 | 1140 |
| 57 | 1740 |
| 56 | 2340 |
| 55 | 2940 |
| 54 | 3540 |
| 53 | 3540 |
| 52 | 3540 |
| 51 | 3540 |
| 50 | 3540 |
| 49 | 3540 |
| 48 | 3540 |
| 47 | 3540 |
| 46 | 3540 |
| 45 | 3540 |
| 44 | 2940 |
| 43 | 2340 |
| 42 | 1740 |
| 41 | 1140 |
| 40 | 1140 |
| 39 | 1140 |
| 38 | 540 |
| 37 | 540 |
| 36 | 290 |
| 35 | 290 |
| 34 | 290 |
| 33 | 290 |
| 32 | 290 |
| 31 | 290 |
| 30 | 290 |
| 29 | 50 |
| 28 | 50 |
| 27 | 50 |
| 26 | 50 |
| 25 | 50 |

**Table S7.**

Average helical parameters, end-to-end and z-distances for the 50mM-DNA-water (3 µs) and DNA-graphene-water (200 ns) systems. Major groove width, minor groove width and rise parameters were averaged over all bases, excluding the terminating two base pairs. The z-distance of the 50 mM DNA-water system was corrected by the average DNA-graphene distance obtained from the DNA-graphene-water simulation (4 Å).

| System | Minor groove width / Å | Major groove width / Å | Rise / Å | End to end distance / Å | z-distance / Å |
|---|---|---|---|---|---|
| 50 mM DNA-water | 12.3 ± 0.5 | 19.5 ± 0.8 | 3.3 ± 0.3 | 112.7 ± 2.5 | 112 ± 6 |
| DNA-graphene-water | 12.3 ± 0.5 | 19.5 ± 0.8 | 3.3 ± 0.3 | 113.0 ± 2.5 | 113.8 ± 2.6 |
